# Low baseline pulmonary levels of cytotoxic lymphocytes as a predisposing risk factor for severe COVID-19

**DOI:** 10.1101/2020.05.04.075291

**Authors:** Pascal H.G. Duijf

**Affiliations:** Institute of Health and Biomedical Innovation, Queensland University of Technology (QUT), Faculty of Health, School of Biomedical Sciences, Brisbane QLD, Australia; Centre for Data Science, Queensland University of Technology (QUT), Brisbane QLD, Australia; University of Queensland Diamantina Institute, The University of Queensland, Translational Research Institute, Brisbane QLD, Australia

**Keywords:** COVID-19, SARS-CoV-2, ACE2, TMPRSS2, T cells, NK cells

## Abstract

COVID-19 is caused by the coronavirus SARS-CoV-2 and currently has detrimental human health, community and economic impacts around the world. It is unclear why some SARS-CoV-2-positive individuals remain asymptomatic, while others develop severe symptoms. Baseline pulmonary levels of anti-viral leukocytes, already residing in the lung prior to infection, may orchestrate an effective early immune response and prevent severe symptoms. Using “*in silico* flow cytometry”, we deconvoluted the levels of all seven types of anti-viral leukocytes in 1,927 human lung tissues. Baseline levels of CD8+ T cells, resting NK cells and activated NK cells, as well as cytokines that recruit these, are significantly lower in lung tissues with high expression of the SARS-CoV-2 entry receptor ACE2. We observe this in univariate analyses, in multivariate analyses, and in two independent datasets. Relevantly, ACE2 mRNA and protein levels very strongly correlate in human cells and tissues. Above findings also largely apply to the SARS-CoV-2 entry protease TMPRSS2. Both SARS-CoV-2-infected lung cells and COVID-19 lung tissues show upregulation of CD8+ T cell- and NK cell-recruiting cytokines. Moreover, tissue-resident CD8+ T cells and inflammatory NK cells are significantly more abundant in bronchoalveolar lavages from mildly affected COVID-19 patients, compared to severe cases. This suggests that these lymphocytes are important for preventing severe symptoms. Elevated ACE2 expression increases sensitivity to coronavirus infection. Thus, our results suggest that some individuals may be exceedingly susceptible to develop severe COVID-19 due to concomitant high pre-existing ACE2 and TMPRSS expression and low baseline cytotoxic lymphocyte levels in the lung.

## Introduction

Coronaviruses are viruses belonging to the family of *Coronaviridae* (1). They are large, single-stranded RNA viruses that often originate from bats and commonly infect mammals. While the majority of coronavirus infections cause mild symptoms, some can cause severe symptoms, such as pneumonia, respiratory failure and sepsis, which may lead to death (2, 3).

Coronavirus zoonosis constitutes a serious health risk for humans. Indeed, in recent history, transmissions of three types of coronaviruses to humans have led to varying numbers of deaths. The outbreak of the Severe Acute Respiratory Syndrome (SARS) epidemic, which is caused by the SARS coronavirus (SARS-CoV), originated in Guangdong, China in 2002 and led to nearly 800 deaths (4). The Middle East Respiratory Syndrome coronavirus (MERS-CoV) outbreak, which emerged in Saudi Arabia in 2012, similarly caused about 800 deaths, although with over 8,000 cases, nearly four times as many cases were reported (4). Finally, Coronavirus Disease 19 (COVID-19), caused by Severe Acute Respiratory Syndrome Coronavirus 2 (SARS-CoV-2), is currently causing a pandemic. On 1 May 2020, the World Health Organization reported over 3 million confirmed cases and over 220,000 patients to have succumbed to COVID-19 around the world (5). However, the factual number of deaths is probably considerably higher (6). In addition, this figure is still soaring, on 1 May 2020 at a rate exceeding 6,400 deaths per day (5).

To infect target cells, coronaviruses use their spike (S) glycoprotein to bind to receptor molecules on the host cell membrane. Angiotensin-converting enzyme 2 (ACE2) has been identified as the main SARS-CoV-2 entry receptor on human cells (7, 8), while the serine protease TMPRSS2, or potentially cathepsin B and L, are used for S-protein priming to facilitate host cell entry (7). SARS-CoV-2 S-protein has a 10 to 20-fold higher affinity to human ACE2 than SARS-CoV S-protein (9). Moreover, ACE2 expression proportionally increases the susceptibility to S protein-mediated coronavirus infection (10–12). Hence, increased expression of ACE2 is thought to increase susceptibility to COVID-19 (13–15).

Epithelial cells of the respiratory tract, including the lung, are primary SARS-CoV-2 target cells (16–18). These cells can sense viral infection via pattern recognition receptors (PRRs). PRRs, including Toll-like receptors and NOD-like receptors, recognize pathogen-associated molecular patterns (PAMPs) (19). Upon PRR activation, a range of pro-inflammatory cytokines and chemokines are produced and released in order to activate the host’s immune system. Interferons (IFNs), in particular type I and type III IFN, are among the principal cytokines to recruit immune cells (19, 20).

Six types of leukocytes have been implicated in detecting and responding to viral infections in the lung, a major site of SARS-CoV-2 infection, which also presents with severe COVID-19 symptoms. The cytotoxic activities of CD8+ T cells and NK cells can facilitate early control of viral infections by clearing infected cells and avoiding additional viral dissemination (21, 22). Dendritic cells specialize in sensing infections, including by viruses, and inducing an immune response (23). CD4+ T cells contribute to viral clearance by promoting production of cytokines and interactions between CD8+ T cells and dendritic cells (24). M1 macrophages interact with pulmonary epithelial cells to fight viral infections in the lung (25). Finally, neutrophils may contribute to clearance of viral infections through phagocytosis of virions and viral particles. However, their precise role is uncertain (26).

SARS-CoV-2 is considerably more efficient in infection, replication and production of infectious virus particles in human lung tissue than SARS-CoV (17). Strikingly, despite this, SARS-CoV-2 initially does not significantly induce type I, II or III IFNs in infected human lung cells and tissue (17, 27). When this does occur, it may in fact promote further SARS-CoV-2 infection, as IFNs directly upregulate expression of the SARS-CoV-2 receptor ACE2 (28). These observations suggest that baseline levels of leukocytes, which already reside in the lung prior to infection, may be important in mounting a rapid immune response against SARS-CoV-2 infection and prevent severe COVID-19 symptoms. As stated above, ACE2 expression level may be a predictor of increased susceptibility to COVID-19 (10–15)}. Thus, we here investigated the relationship between ACE2 and TMPRSS2 expression and the levels of seven leukocyte types implicated in anti-viral immune response in human lung tissue.

## Results

We used bulk RNAseq gene expression data from the 578 human lung tissues present in the Genotype-Tissue Expression (GTEx) database (29, 30), because this is the largest publicly available dataset with clinical information. Using an established “*in silico* flow cytometry” pipeline (31), we estimated the levels of CD8+ T cells, resting and activated NK cells, M1 macrophages, dendritic cells, CD4+ T cells and neutrophils in these tissues **(Fig. S1a-c**, **Table S1)**. We compared these to ACE2 expression levels in these lung tissues. This revealed that ACE2 expression is negatively correlated with the levels of CD8+ T cells, resting and activated NK cells and M1 macrophages (*p* < 8×10^−6^, Pearson correlations) **(Fig. 1a-c)**. However, there are no statistically significant correlations between ACE2 expression and the levels of CD4+ T cells, dendritic cells and neutrophils (*p* > 0.05) **(Fig. S2a-d)**. Thus, the levels of a majority of leukocytes involved in anti-viral immune responses are significantly lower in lung tissues with high ACE2 expression levels.

**Figure 1.**
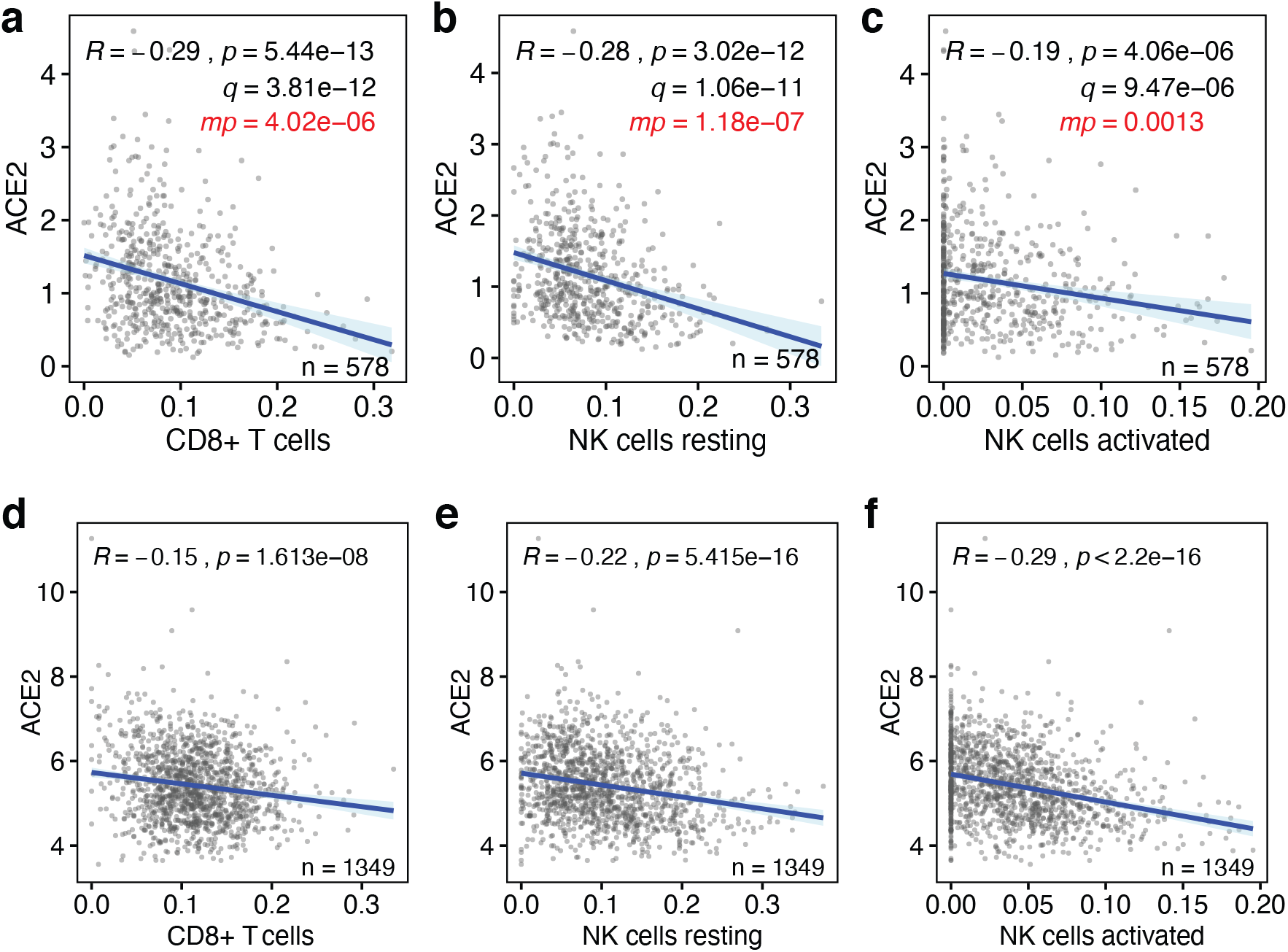
Baseline levels of cytotoxic lymphocytes inversely correlate with ACE2 expression in the lung. **(a-c)** The correlations between the baseline levels of CD8+ T cells, resting NK cells and activated NK cells (x-axes) and the those of the SARS-CoV-2 host cell receptor ACE2 (y-axes) in human lung tissue are shown. Data are from the GTEx dataset (n=578). **(d-f)** The same correlations are shown for lung tissues in the LUG dataset (n=1,349). Regression lines and 95% confidence intervals are shown. *R* and *p* values: Pearson correlations; *q* values: Benjamini-Hochberg-adjusted *p* values using a false discovery rate of 0.05; *mp* values: multivariate *p* values (see Table S3).

It is possible that some of above observations are linked to phenotypic characteristics, such as sex, age, body mass index (BMI), race or smoking status. To test the robustness of our findings, we applied multivariable regression analysis that includes these five covariates **(Table S2)**, as well as the levels of the seven above leukocyte types or states. This showed that only 4 of the 12 variables significantly contribute to predicting ACE2 expression levels, specifically the levels of CD8+ T cells, resting NK cells, activated NK cells and M1 macrophages **(Fig. 1a-c, Table S3)**. Notably, none of the five added phenotypic covariates showed statistically significant contributions. Consistently, we found limited statistically significant correlations between these variables and ACE2 expression in univariate analyses, irrespective of whether they were analyzed as continuous data or binned into discrete ordinal categories **(Fig. S4a-j)**. Thus, the levels of four types of leukocytes that respond to viral infection are low in lung tissue with high ACE2 expression levels independently of phenotypic covariates.

Next, we tested whether above observations could be validated in an independent cohort of individuals. For this, we used the, to our knowledge, largest publicly available lung tissue dataset. The Laval University, University of British-Columbia, Groningen University (LUG) dataset includes microarray gene expression data of 1,349 human lung tissues. Following determination of ACE2 expression levels and estimation of the levels of CD8+ T cells, resting NK cells, activated NK cells and M1 macrophages **(Fig. S1a-c**, **Table S1)**, we found that three of the four also negatively correlated with ACE2 expression in this independent dataset (*p*<2×10^−8^) **(Fig. 1d-f)**. With a correlation coefficient of *R*=0.096, only M1 macrophages did not correlate with ACE2 expression in this dataset **(Fig. S3)**. Thus, our observations indicate that the baseline levels of three types of cytotoxic lymphocytes, specifically CD8+ T cells, resting NK cells and activated NK cells, are robustly and consistently low in lung tissue with high expression of the SARS-CoV-2 receptor ACE2.

To more rigorously assess our observations, we employed a range of additional analyses. Although highly statistically significant (all *p* values <4.1×10^−6^, **Fig. 1**), the absolute Pearson *R* values between baseline levels of ACE2 and the three lymphocyte types were seemingly low, as they ranged between 0.2 and 0.3 **(Fig 1a-c)**. To test how strong these are in relative terms, we calculated the Pearson *R* and *p* values of 1000 randomly sampled other genes. This revealed that the ACE2 *R* and *p* values were significantly lower than expected by chance (all one-sample *t* test *p*<2.2×10^−16^; **Fig 2a,b**). In addition, these *R* and *p* values ranked in the top 0.4 to 11 percentiles of strongest and most significant correlations for each of the three leukocyte types **(Fig 2a,b)**. Thus, the seemingly low correlations between ACE2 mRNA and cytotoxic lymphocyte levels in the lung are not only highly statistically significant, but also strong in relative terms.

**Figure 2.**
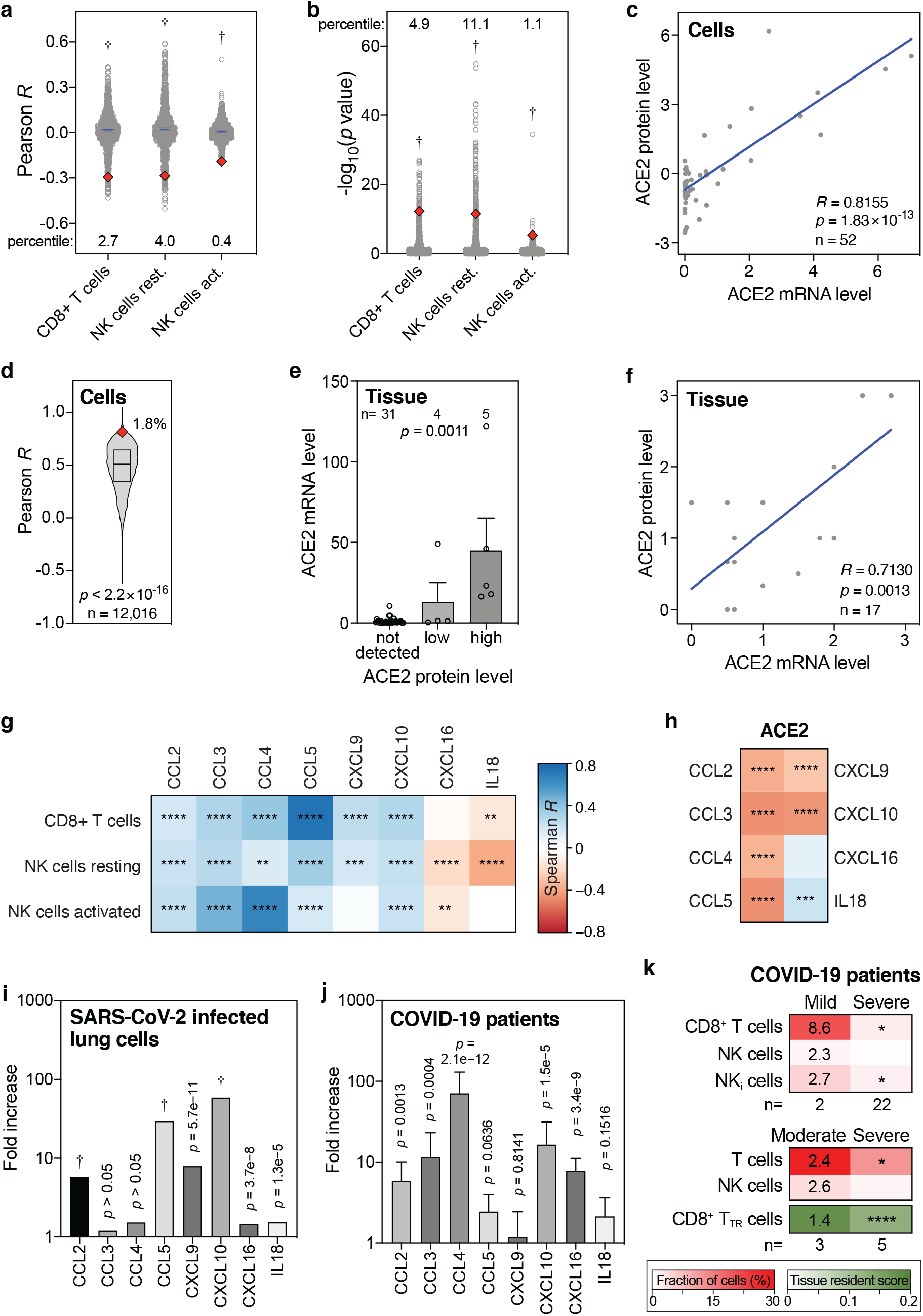
Levels of ACE2 mRNA, ACE2 protein, CD8+ T cells, NK cells and cytokines in lung cells, lung tissues and COVID-19 patient samples. **(a, b)** Pearson *R* and −log_10_(*p* values) of correlations between 1000 randomly sampled genes and the levels of indicated lymphocytes in lung tissues were determined and plotted. ACE2 Pearson *R* and *p* values are shown as red diamonds. Blue lines indicate means with 95% confidence intervals. Percentiles for ACE2 with respect to the 1000 random *R* and *p* values are shown. Data are from the GTEx dataset (n=578). *P* values: one-sample *t* tests. **(c)** Correlations between ACE2 mRNA and protein levels in 52 cell lines. *R* and *p* values: Pearson correlations. **(d)** Pearson correlation *R* values between mRNA and protein levels of 12,016 genes are compared to the ACE2 *R* coefficient (red diamond). The box represents the median and interquartile range. The ACE2 *R* percentile is also shown. *P* value: one-sample *t* test. **(e)** Bar graph showing the correlation between ACE2 mRNA and protein levels in human tissues. Means with standard errors of the means are shown. Samples are from the Human Protein Atlas. *P* value: Kruskal-Wallis test. **(f)** Meta-analysis scatter plot showing the correlation between ACE2 mRNA and protein levels in 17 human tissues. Data are from 9 different studies, as detailed in Table S4. *P* value: Pearson correlation. **(g, h)** Heatmaps showing Spearman correlations between the levels of ACE2 or indicated cytotoxic lymphocytes and eight cytokines that recruit these cells in human lung tissues from the GTEx dataset (n=578). Colors of tiles represent Spearman *R*, as per scale bar on the right. Spearman significance levels are shown by asterisks. See also Fig. S5. **(i, j)** Fold increase in expression levels of indicated cytokines in Calu-3 lung cells 24 hours after SARS-CoV-2 infection compared to uninfected Calu-3 cells (i), and in post-mortem COVID-19 lung tissues (n=2) to those in healthy, uninfected lung tissues (n=2) (j). **(k)** Comparison of indicated fractions of lymphocyte levels in bronchoalveolar lavages of mild/moderate and severe COVID-19 patients, as determined in two separate studies (46, 47). The second also determined a tissue resident (TR) score for CD8+ T cells. Numbers in the mild/moderate column on the left show fold increase compared to the respective severe cases on the right. Asterisks in the severe column on the right represent statistical significance levels, as determined my Mann-Whitney *U* tests (top panel), or *t* tests (middle and bottom panels) comparing mild/moderate to severe cases. *P* value abbreviations: *, *p*<0.05; **, *p*<0.01; ***, *p*<0.001; ****, *p*<0.0001; †, *p*<2.2×10^−16^.

Further, we tested how well ACE2 mRNA and protein levels correlate. Using the mRNA and protein levels in 52 cell lines, we find that ACE2 mRNA levels strongly correlate with ACE2 protein levels in human cells (Pearson *R*=0.8155, *p*=1.8×10^−13^; **Fig. 2c**). In fact, ACE2 ranks in the top 1.8 percentile of over 12,000 genes with the strongest mRNA-protein level correlations (*p*<2.2×10^−16^; **Fig 2d**). Using two ACE2-specific antibodies, immunochemistry on 40 human tissues also shows a strong ACE2 mRNA-protein correlation (*p*=0.0011, Kruskal-Wallis test; **Fig. 2e**) and this is additionally validated by a meta-analysis that we conducted using nine published studies (Pearson *R*=0.7130, *p*=0.0013; **Fig. 2f**, **Table S4**). Therefore, we conclude that ACE2 mRNA and protein levels very strongly correlate, both in human cells and in human tissues.

Above, we found that baseline ACE2 levels in the lung negatively correlate with CD8+ T cells and resting and activated NK cells in multivariate analyses and in an independent dataset **(Fig. 1a-f)**. Several cytokines, including CCL2-5, CXCL9, CXCL10, CXCL16 and IL18, are known to chemotactically attract CD8+ T cells and NK cells (32–37). Consistently, we find that the baseline levels of these chemokines in human lung tissue typically significantly correlate with the baseline levels of CD8+ T cells and resting and activated NK cells (**Fig. 2g, Fig. S5**). Additionally, as expected given our above results, we find significant negative correlations between the levels of ACE2 and the levels of six of these eight cytokines in the lung **(Fig. 2h)**. These findings lend further support to our previous observations, suggesting that high levels of said cytokines in the lung establish a favorable milieu for cytotoxic lymphocytes, which correlates with low ACE2 levels.

Next, we assessed the direct consequences of SARS-CoV-2 infection. *In vitro* SARS-CoV-2 infection of human lung cells invariably leads to upregulation of all eight abovementioned CD8+ T cell- and NK cell-attracting cytokines, with six of these increases showing statistical significance **(Fig. 2i)**. Similarly, compared to control lung tissues, all eight cytokines are upregulated in lung tissues from COVID-19 patients, with five showing statistical significance **(Fig. 2j)**. Moreover, the levels of CD8+ T cells and NK cells are higher in bronchoalveolar lavages of mildly affected COVID-19 patients than in severe cases, with CD8+ T cells and a subset of NK cells, inflammatory NK cells, showing a statistically significant higher level **(Fig. 2k)**. These findings are corroborated in a different cohort of patients, additionally showing a highly significant increase in a tissue-resident signature score for CD8+ T cells **(Fig. 2k)**. Thus, together, these observations suggest that SARS-CoV-2 infection of lung cells stimulates CD8+ T cell- and NK cell-attracting cytokines and that these cytotoxic lymphocytes are important for preventing severe symptoms of COVID-19.

Finally, we tested whether the levels of the SARS-CoV-2 host cell protease TMPRSS2 shows similar correlations with the levels of CD8+ T cells and NK cells in the lung. In univariate analyses, baseline TMPRSS2 levels in the lung show significant negative correlations with these lymphocyte levels, although in multivariate analyses, these are only statistically significant for CD8+ T cells and activated NK cells **(Fig. 3a-c)**. The corresponding *R* and *p* values are typically also significantly lower than expected by chance **(Fig. 3d,e)**. Furthermore, TMPRSS2 mRNA and protein levels strongly correlate (*R*=0.8048, *p*<2.2×10^−16^, **Fig. 3f**) and TMPRSS2 is in the top 2.5 percentile of genes that show the strongest mRNA-protein correlation (*p*<2.2×10^−16^, **Fig. 3g**). Additionally, TMPRSS2 expression tends to correlate negatively with CD8+ T cell- and NK cell-attracting cytokines **(Fig. 3h**). Therefore, albeit typically to a lesser extent, baseline TMPRSS2 expression levels in the lung negatively correlate with the levels of CD8+ T cells and NK cells in a manner similar to ACE2.

**Figure 3.**
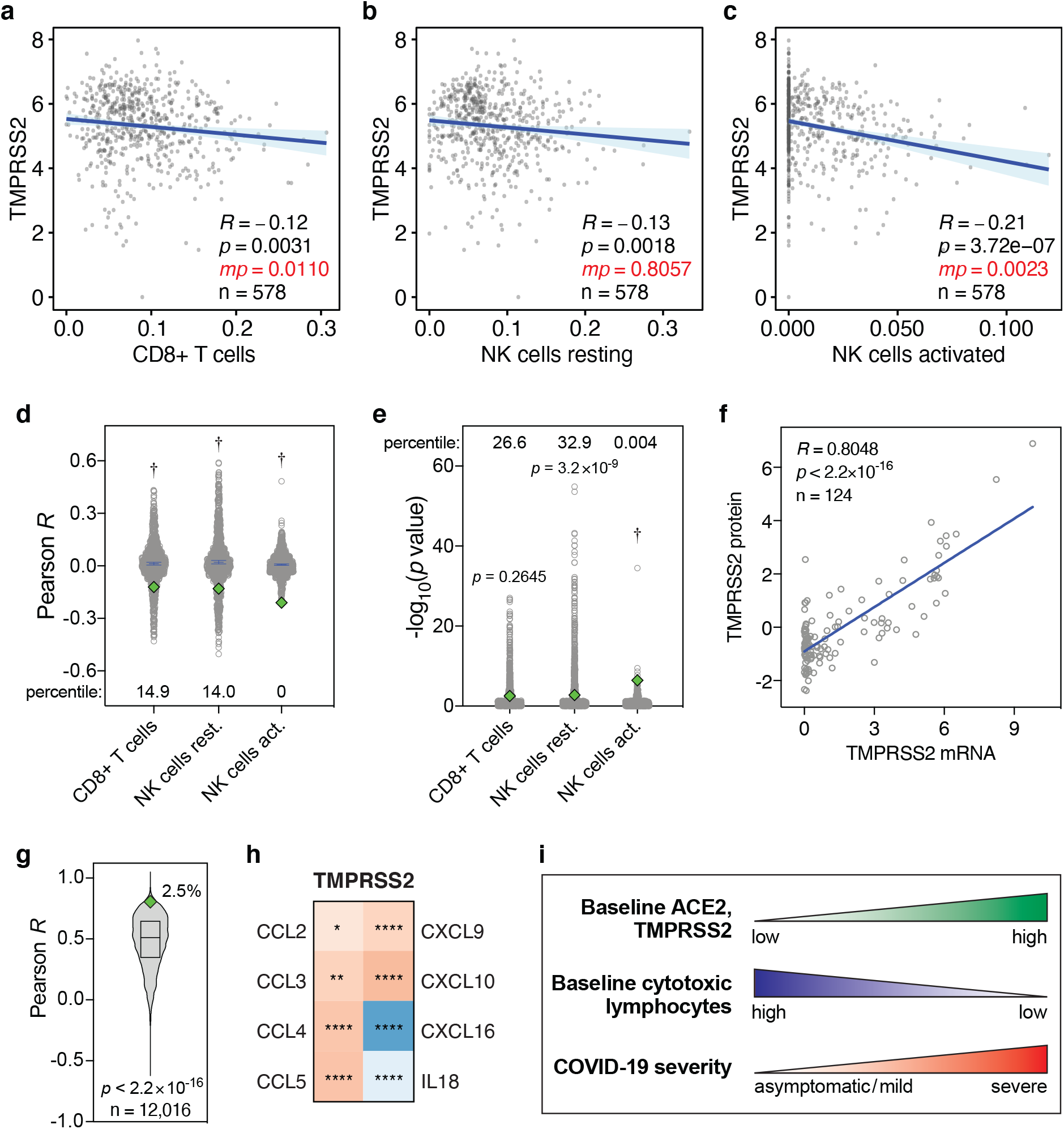
Levels of TMPRSS2 mRNA, TMPRSS2 protein and cytokines in lung cells and tissues. **(a-c)** Pearson correlations between baseline levels of indicated lymphocytes and TMPRSS2 in human lung tissue, as Figure 1a-c. Data are from the GTEx dataset (n=578). **(d, e)** Pearson *R* and −log_10_(*p* values) of correlations between 1000 randomly sampled genes and the levels of indicated lymphocytes in lung tissues, as in Figure 2a,b. TMPRSS2 Pearson *R* and *p* values are shown as green diamonds. **(f)** Correlations between TMPRSS2 mRNA and protein levels in 124 cell lines. *R* and *p* values: Pearson correlations. **(g)** Pearson correlation *R* values between mRNA and protein levels of 12,016 genes are compared to the TMPRSS2 *R* coefficient (green diamond). The box represents the median and interquartile range. The TMPRSS2 *R* percentile is also shown. *P* value: one-sample *t* test. **(h)** Heatmap showing Spearman correlations between the levels of TMPRSS2 and cytokines in human lung tissues from the GTEx dataset (n=578), as in Figure 2h. **(i)** Individuals with high baseline levels of ACE2 and TMPRSS2 show low baseline tissue-resident levels of cytotoxic lymphocytes in the lung. We propose that this may jointly predispose these individuals to development of severe COVID-19.

Taken together, our observations suggest that a subgroup of individuals may be exceedingly susceptible to developing severe COVID-19 due to concomitant high pre-existing ACE2 and TMPRSS2 expression and low baseline levels of CD8+ T cells and NK cells in the lung **(Fig. 3i)**.

## Discussion

We investigated the baseline expression levels of the SARS-CoV-2 host cell entry receptor ACE2 and the host cell entry protease TMPRSS2 and the baseline levels of all seven types of anti-viral leukocytes in 1,927 human lung tissue samples. Although SARS-CoV-2 cellular tropism is broad (16–18), we focused on lung tissue. In addition to epithelial cells elsewhere in the respiratory tract, alveolar epithelial cells are thought to be a primary SARS-CoV-2 entry point (16, 28). Consistently, the SARS-CoV-2 receptor ACE2 is expressed in these cells at the mRNA and protein levels (28, 38–40). Moreover, in severely affected COVID-19 patients, the lungs are among the few organs that present with the most life-threatening symptoms. “Cytokine storm”-induced acute respiratory distress syndrome (ARDS), widespread alveolar damage, pneumonia, and progressive respiratory failure have been observed (41, 42). These indications frequently require admission to intensive care units (ICUs) and mechanical ventilation and may ultimately be fatal.

Early after infection, rapid activation of the innate immune system is of paramount importance for the clearance of virus infections. Infected cells typically do so through release of pro-inflammatory cytokines and chemokines, in particular type I and III interferons (19, 20). Notably, however, several studies have highlighted multiple complexities related specifically to SARS-CoV2 and innate immune system activation at early stages. First, unlike SARS-CoV, SARS-CoV-2-infected lung tissue initially fails to induce a range of immune cell-recruiting molecules, including several interferons (17, 27), suggesting that leukocytes are ineffectively recruited to infected lung shortly after infection. Second, the host cell entry receptor ACE2 has been identified as an interferon target gene (28). Thus, even when interferons are upregulated in order to recruit immune cells, concomitant upregulation of ACE2 expression may in fact exacerbate SARS-CoV-2 infection (28).

These findings suggest that the levels of immune cells that already reside in the lung prior to infection are more critical for dampening SARS-CoV-2 infection at early stages than they are for fighting infections of other viruses. Cytotoxic lymphocytes, including CD8+ T cells and NK cells, are key early responders to virus infections and these are the cells whose baseline levels we here identify as significantly reduced in lung tissue with elevated ACE2 and TMPRSS2 expression. That these immune cells are important in preventing severe COVID-19 is supported by the fact that their levels are significantly higher in bronchoalveolar lavages from mild patients than from severe patients. Therefore, our results suggest that individuals with increased baseline susceptibility to SARS-CoV-2 infection in the lungs may also be less well equipped from the outset to mount a rapid anti-viral cellular immune response **(Fig. 3i)**.

Several observations indicate that these cytotoxic lymphocytes are critically important for effective control of SARS-CoV-2 infection. Recent studies showed that CD8+ T cells in peripheral blood are considerably reduced and functionally exhausted in COVID-19 patients, in particular in elderly patients and in severely affected patients that require ICU admission (43–45). Reduced CD8+ T cell counts also predict poor COVID-19 patient survival (43). Additionally, CD8+ T cell- and NK cell-attracting cytokines are upregulated in SARS-CoV-2-infected human lung cells and in lung tissues from COVID-19 patients and the levels of CD8+ T cells and NK cells are higher in bronchoalveolar lavages of mildly affected COVID-19 patients than in severe cases (46, 47).

We found that the five phenotypic parameters, sex, age, BMI, race and smoking history did not statistically significantly contribute to variation in ACE2 expression in human lung tissue, neither in univariate nor in multivariate analyses. This is consistent with some studies but inconsistent with others (42, 48–50). These paradoxical observations may be partially explained by varying gender, age and race distributions within each study cohort.

Further research will be required to elucidate the precise mechanisms of SARS-CoV-2-induced activation of the innate immune system early after infection. However, the link that we identified between high baseline ACE2 and TMPRSS2 expression and reduced cytotoxic lymphocyte levels in human lung tissue prior SARS-CoV-2 infection is striking. It suggests that increased susceptibility to SARS-CoV-2 infection in the lungs may be accompanied by a poorer ability to mount a rapid innate immune response at early stages. This may predict long-term outcome of individuals infected with COVID-19, given that the levels of CD8+ T cell and NK cell are significantly higher in bronchoalveolar lavages of mild cases compared to severe patients (46, 47). Finally, it may contribute to the substantial variation in COVID-19 clinical presentation, ranging from asymptomatic to severe respiratory and other symptoms.

## Methods

### Discovery dataset and processing

Gene expression data and corresponding phenotype data from human lung tissues (n=578) were obtained from the Genotype-Tissue Expression (GTEx) Portal (https://gtexportal.org), managed by the National Institutes of Health (NIH). Gene expression data were publicly available. Access to phenotype data required authorization. The GTEx protocol was previously described (29, 30). Briefly, total RNA was extracted from tissue. Following mRNA isolation, cDNA synthesis and library preparation, samples were subjected to HiSeq2000 or HiSeq2500 Illumina TrueSeq RNA sequencing. Gene expression levels were obtained using RNA-SeQC v1.1.9 (51) and expressed in transcripts per million (TPM). Reported expression levels were log_2_-transformed, unless otherwise indicated.

### Validation dataset and processing

For validation purposes, the Laval University, University of British-Columbia, Groningen University (LUG) lung tissue dataset (n=1,349) was used. This dataset was accessed via Gene Expression Omnibus (GEO; https://www.ncbi.nlm.nih.gov/geo), accession number GSE23546, and was previously described (52). Briefly, total RNA from human lung tissue samples was isolated, quantified, quality-checked and used to generate cDNA, which was amplified and hybridized to Affymetrix gene expression arrays. Arrays were scanned and probe-level gene expression values were normalized using robust multichip average (RMA). These normalized values were obtained from GEO. To collapse probe-level expression data to single expression levels per gene, for each gene the probe with the highest median absolute deviation (MAD) was used. The MAD for each probe *p* was calculated using equation (1).

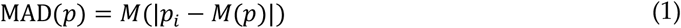

Herein, *M* is the median, *p*_*i*_ denotes probe *p*’s expression level in sample *i*, and *M*(*p*) represents the median signal of probe *p*.

### *In silico* cytometry

The levels of seven types of leukocytes involved in anti-viral cellular immune response, specifically CD8+ T cells, resting NK cells, activated NK cells, M1 macrophages, CD4+ T cells, dendritic cells and neutrophils, were estimated in the discovery and validation lung tissue samples using a previously described approach (31). Specifically, the following workflow was used. First, only non-log-transformed expression values were used. Thus, where required, expression values for all samples in the discovery and validation datasets were reverse-log_2_-transformed using equation 2.

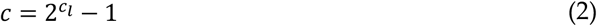

Herein, *c* denotes the calculated non-log_2_-transformed expression counts and *c*_*l*_ denotes the previously reported log_2_-transformed expression counts. Next, to compensate for potential technical differences between signatures and bulk sample gene expression values due to inter-platform variation, bulk-mode batch correction was applied. To ensure robustness, deconvolution was statistically analyzed using 100 permutations. Pearson correlation coefficients *R*, root mean squared errors (RMSA) and *p* values are reported on a per-sample level in **Fig. S1a-c** and **Table S1**.

### Univariate statistical analyses

Log_2_-transformed expression levels of ACE2 in lung tissue samples were compared to the estimated levels of seven leukocyte types or states. Pearson correlation analyses were performed to determine Pearson correlation coefficients *R* and *p* values. *P* values were adjusted at a false discovery rate of 0.05 to yield *q* values, as previously described (53). Straight lines represent the minimized sum of squares of deviations of the data points with 95% confidence intervals shown. Continuous phenotypic covariates were analyzed in the same way and, additionally, as discrete ordinal categories after binning. Discrete and binned phenotype data were statistically evaluated using Mann-Whitney *U* tests. All analyses were performed in the *R* computing environment (*R* Project for Statistical Computing, Vienna, Austria).

### Multivariate regression analyses

Multivariate analyses were performed using standard ordinary least squares regression, summarized in equation 3.

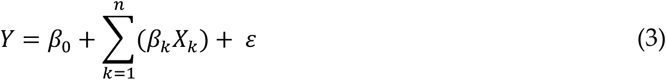

Herein, *β*_0_ denotes the intercept, while *β*_*k*_ represents the slope of each variable *X*_*k*_ in a model with *n* variables and *ε* denotes the random error component. These analyses was performed using *R*.

### Messenger RNA and protein levels in human cells

Available ACE2 and TMPRSS2 mRNA and corresponding protein levels in 52 and 124 human cell lines, respectively, were obtained from references (54, 55). The correlations between mRNA and protein levels were analyzed by linear regression analysis using Pearson correlations. Pearson coefficients *R* for mRNA-protein level correlations were also determined for 12,015 other genes. To test whether the ACE2 and TMPRSS2 coefficients and *p* values were statistically significantly lower than for other genes, one-sample *t* tests were used.

### Messenger RNA and protein levels in human tissues

Two types of analyses were performed to compare mRNA and protein levels in human tissues. First, mRNA expression levels from 40 human tissues were obtained from the Human Protein Atlas (https://www.proteinatlas.org). These represented the consensus normalized mRNA expression levels from three sources, specifically, Human Protein Atlas in-house RNAseq data, RNAseq data from the Genotype-Tissue Expression (GTEx) project and CAGE data from FANTOM5 project. Corresponding ordinal human tissue protein expression levels (‘not detected’, ‘low’, ‘high’) were also obtained from the Human Protein Atlas. These were based on immunohistochemical staining of the tissues using DAB (3,3’-diaminobenzidine)-labeled antibodies (HPA000288, CAB026174), followed by knowledge-based annotation, as described on said website. A Kruskal-Wallis test was performed to assess whether mRNA and protein levels significantly correlated. Second, a meta-analysis was performed. For this, mRNA and protein expression levels were obtained from nine different sources. Tissue mRNA levels were obtained from six sources, as determined by Northern blot (56), quantitative RT-PCR (57), microarray hybridization (58), RNA sequencing (59) and cap analysis of gene expression (60). Tissue protein levels were obtained from three sources, as determined by mass spectrometry (61) and immunohistochemistry (39, 62). For each source and tissue, mRNA and protein expression levels were scored as ‘not detected’, ‘low’, ‘intermediate’ or ‘high’ and these received scores of 0-3, respectively. For each tissue, final mRNA and protein scores were calculated by averaging and scores from the respective six and three sources **(Table S4)**. Strength and significance level of the correlation between the final scores were determined by linear regression analysis using Pearson correlation.

### Cytokine levels in SARS-CoV-2-infected lung cells

SARS-CoV-2-induced fold increases in the expression levels of eight cytotoxic lymphocyte-attracting cytokines, CCL2-5, CXCL9, CXCL10, CXCL16 and IL18, were determined from reference (27). These represent fold increase in expression in Calu-3 lung cells, 24 hours after infection with SARS-CoV-2 at a multiplicity of infection of 2, compared to uninfected Calu-3 cells.

### Cytokine levels in control and COVID-19 lung tissues

Baseline levels of above eight cytokines in lung tissues were obtained from the GTEX project, as described above, and compared to the baseline levels of ACE2, TMPRSS2, CD8+ T cells, resting and activated NK cells in these tissues, estimated as described above. Their levels were compared using Spearman’s rank correlations (*R* and *p* values). For comparison of cytokine levels in post-mortem COVID-19 lung tissues (n=2) to those in healthy, uninfected lung tissues (n=2), fold increases were determined following RNAseq analyses and previously reported (27).

### Lymphocyte levels in COVID-19 bronchoalveolar lavages

The levels of CD8+ T cells, NK cells and inflammatory NK cells in bronchoalveolar lavages from mild (n=2) and severe (n=22) COVID-19 patients were reported elsewhere and determined using single-cell RNAseq (46). Statistical significance levels were assessed using Mann-Whitney *U* tests. The levels of T cells and NK cells, as well as CD8+ T cell tissue-resident signature score, in bronchoalveolar lavages from moderate (n=3) and severe/critical (n=5) COVID-19 patients were reported in another study (47).

## Acknowledgements

I thank Dr. Marianna Datseris, M.D. for critically reading the manuscript. This work was supported by the School of Biomedical Sciences, Queensland University of Technology. This study used publicly available and controlled data from the Genotype-Tissue Expression (GTEx) Project. GTEx was supported by the Common Fund of the Office of the Director of the National Institutes of Health (commonfund.nih.gov/GTEx). Additional funds were provided by the NCI, NHGRI, NHLBI, NIDA, NIMH, and NINDS. Donors were enrolled at Biospecimen Source Sites funded by NCI\Leidos Biomedical Research, Inc. subcontracts to the National Disease Research Interchange (10XS170), Roswell Park Cancer Institute (10XS171), and Science Care, Inc. (X10S172). The Laboratory, Data Analysis, and Coordinating Center (LDACC) was funded through a contract (HHSN268201000029C) to the The Broad Institute, Inc. Biorepository operations were funded through a Leidos Biomedical Research, Inc. subcontract to Van Andel Research Institute (10ST1035). Additional data repository and project management were provided by Leidos Biomedical Research, Inc. (HHSN261200800001E). The Brain Bank was supported supplements to University of Miami grant DA006227. Statistical Methods development grants were made to the University of Geneva (MH090941 & MH101814), the University of Chicago (MH090951, MH090937, MH101825, & MH101820), the University of North Carolina - Chapel Hill (MH090936), North Carolina State University (MH101819), Harvard University (MH090948), Stanford University (MH101782), Washington University (MH101810), and to the University of Pennsylvania (MH101822). The datasets used for the analyses described in this study were obtained from dbGaP at http://www.ncbi.nlm.nih.gov/gap through dbGaP accession number phs000424.v8.p2.

## Supplementary Information

**Figure S1.**
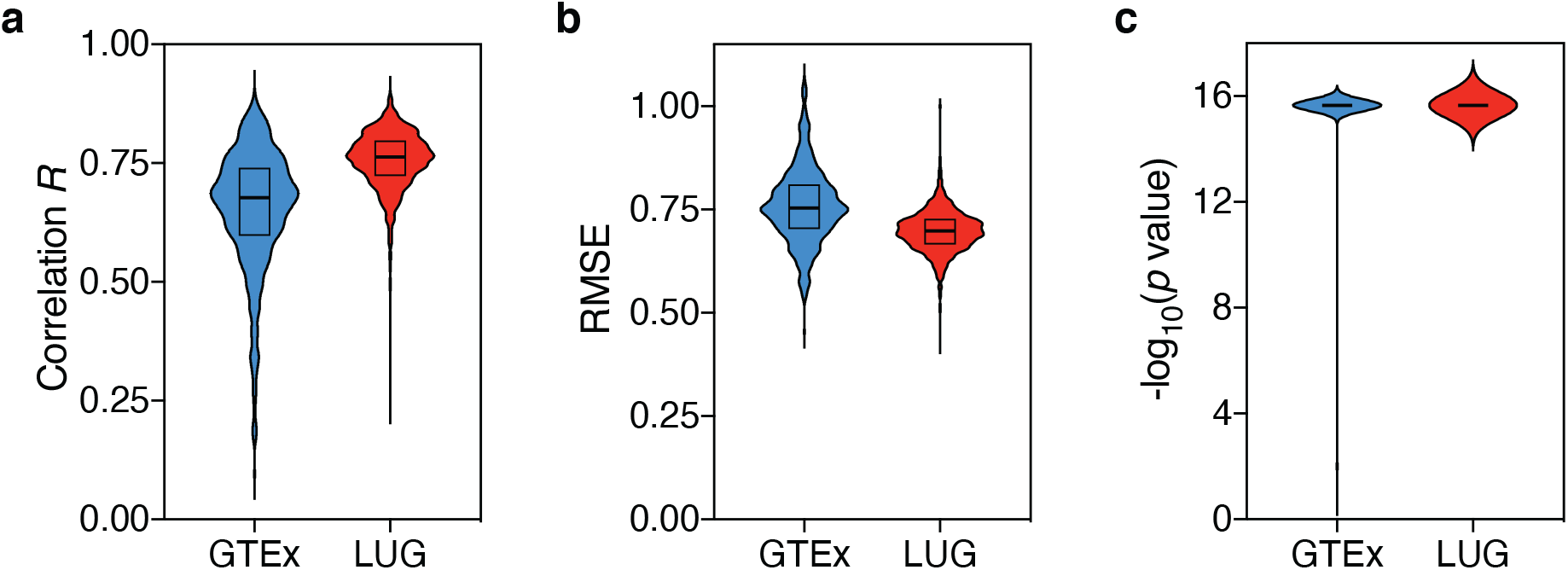
*In silico* cytometry statistics. Per-sample statistics of *in silico* cytometry on human lung tissues are shown for the GTEx (n=578) and LUG (n=1,349) datasets. **(a)** Pearson correlation coefficients *R*. **(b)** Root mean square errors (RMSE). **(c)** *p* values. *P* values < 2.2×10^−16^ were processed as equal to 2.2×10^−16^. Source data are provided in Table S1.

**Figure S2.**
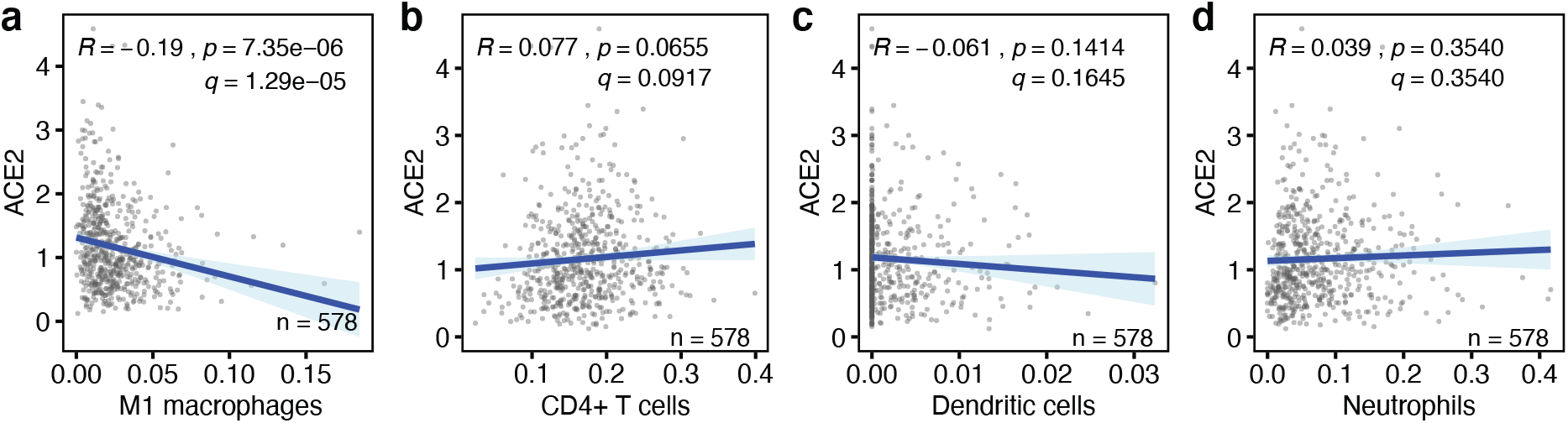
Correlations between baseline anti-viral leukocyte levels and ACE2 expression levels in human lung tissues. **(a-d)** The correlations between the baseline levels of indicated immune cell types (x-axes) and the expression level of the SARS-CoV-2 host cell receptor ACE2 (y-axis) in human lung tissue are shown. Data are from the GTEx dataset (n=578). Regression lines and 95% confidence intervals are shown. *R* and *p* values: Pearson correlations; *q* values: Benjamini-Hochberg-adjusted *p* values using a false discovery rate of 0.05. See also Figure 1.

**Figure S3.**
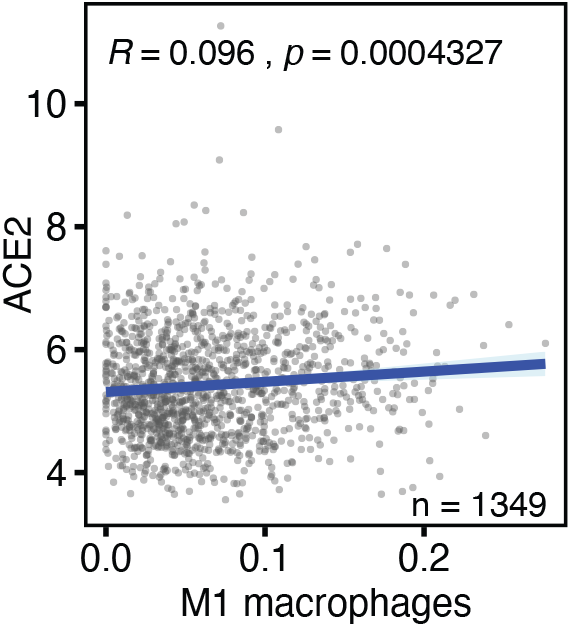
Correlation between baseline M1 macrophage levels and ACE2 expression level in human lung tissue. The correlation between these variables is shown as described in Figure S2. Data are from the LUG dataset (n=1,349).

**Figure S4.**
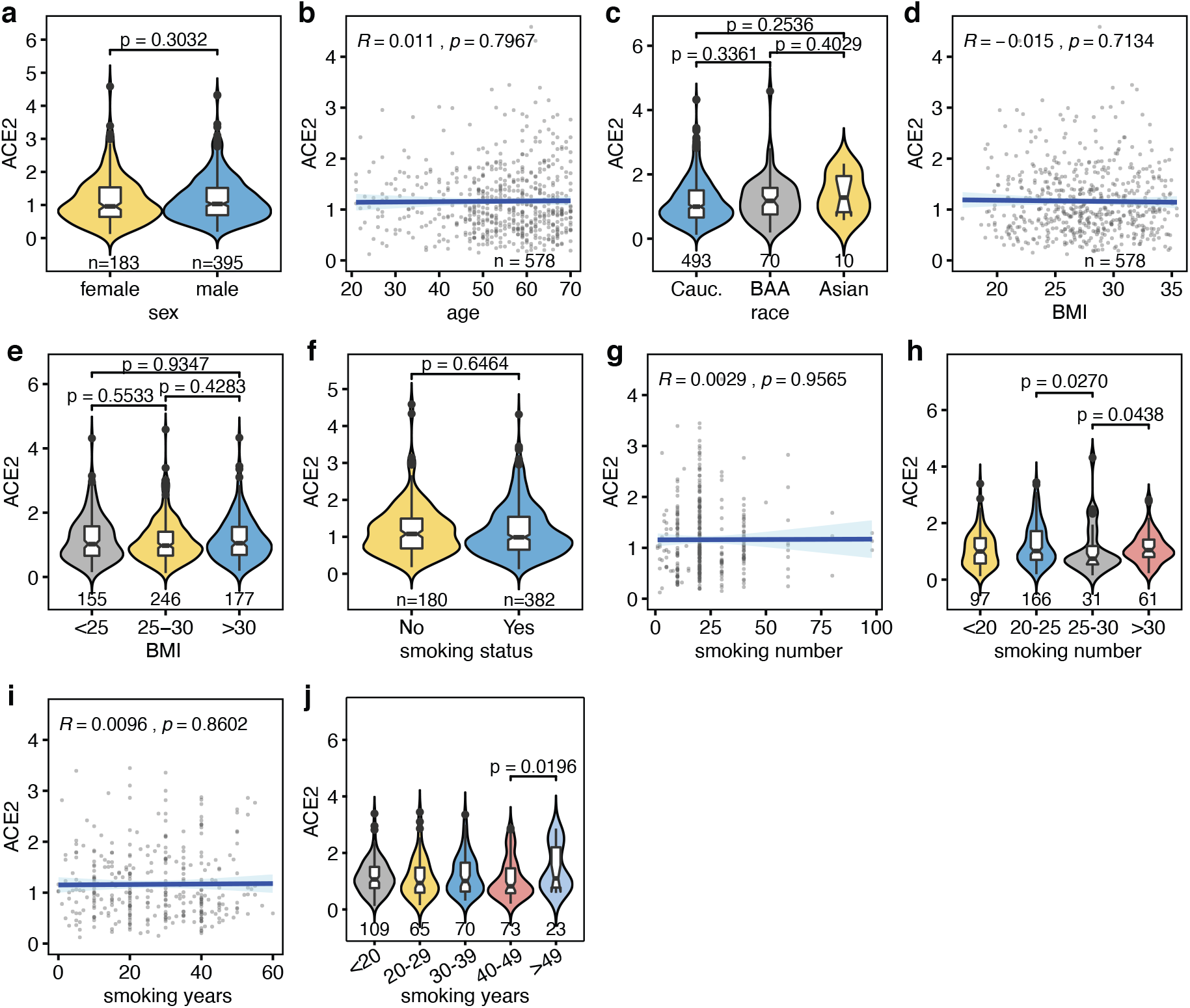
Univariate analyses of phenotypic covariates included in multivariate analyses. Univariate analyses of five covariates that were included in multivariate analyses of ACE2 and TMPRSS2 expression in human lung tissue using the GTEx dataset (Tables S2, S3). **(a)** Sex. **(b)** Age, **(c)** Race. **(d, e)** Body mass index. **(f-j)** Smoking behavior, referring to smoking status (smoker/non-smoker) (f), number of units smoked during the smoke period (g, h) and number of years smoked (i, j). *P* values in categorical analyses: Mann-Whitney *U* tests. *R* and *p* values in continuous analyses: Pearson correlations. *P* values in panels h and j are only shown if *p*<0.05. Sample numbers (n) are shown on the x-axes.

**Figure S5.**
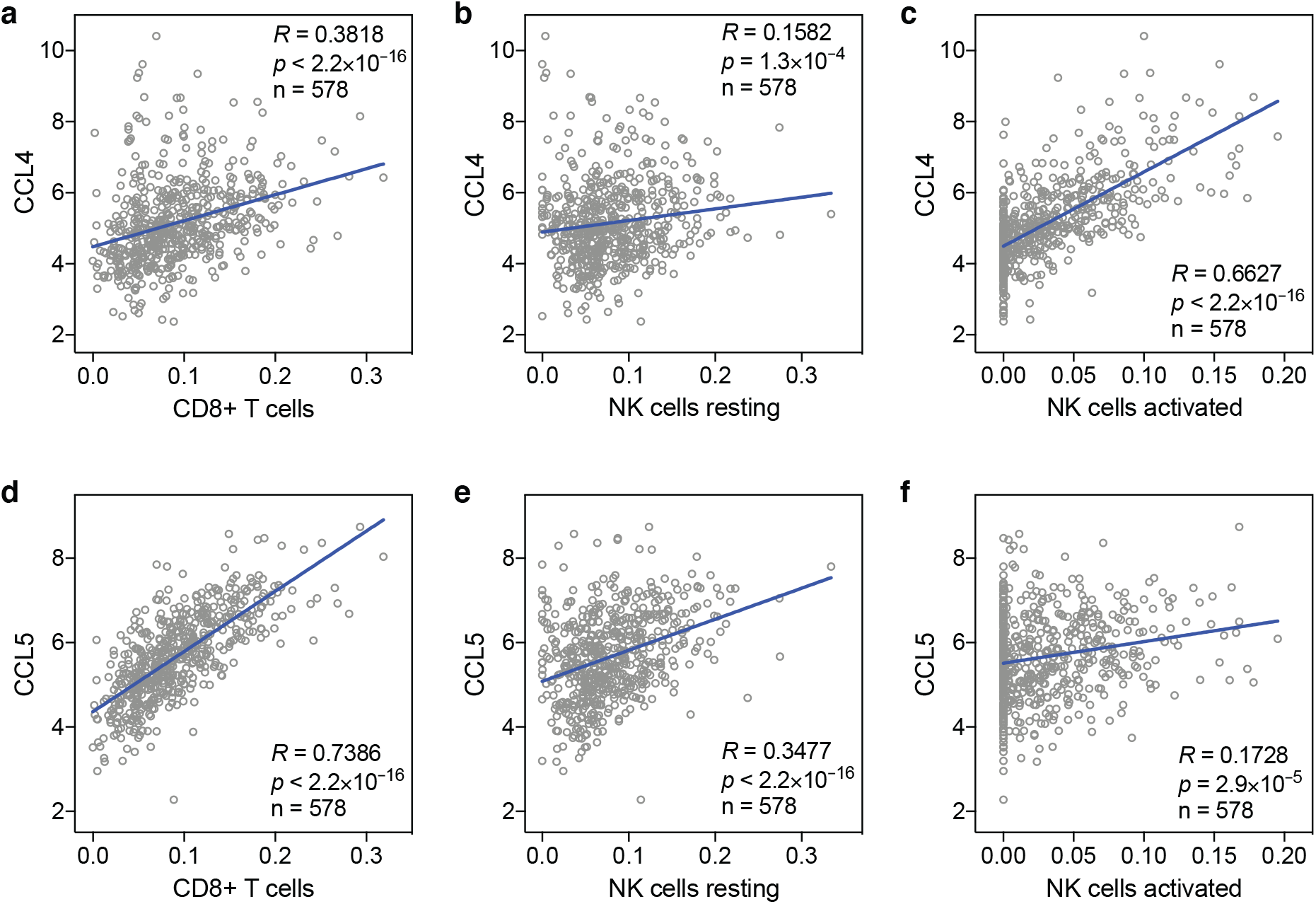
Correlations between baseline levels of cytotoxic lymphocytes and CCL4 and CCL5 in human lung tissue. The correlations between baseline levels of the chemokines CCL4, CCL5 and CD8+ T cells, resting and activated NK cells in human lung tissue are shown. Data are from the GTEx dataset (n=578). *R* and *p* values: Spearman correlations. See also Figure 2g.

**Table S1.**
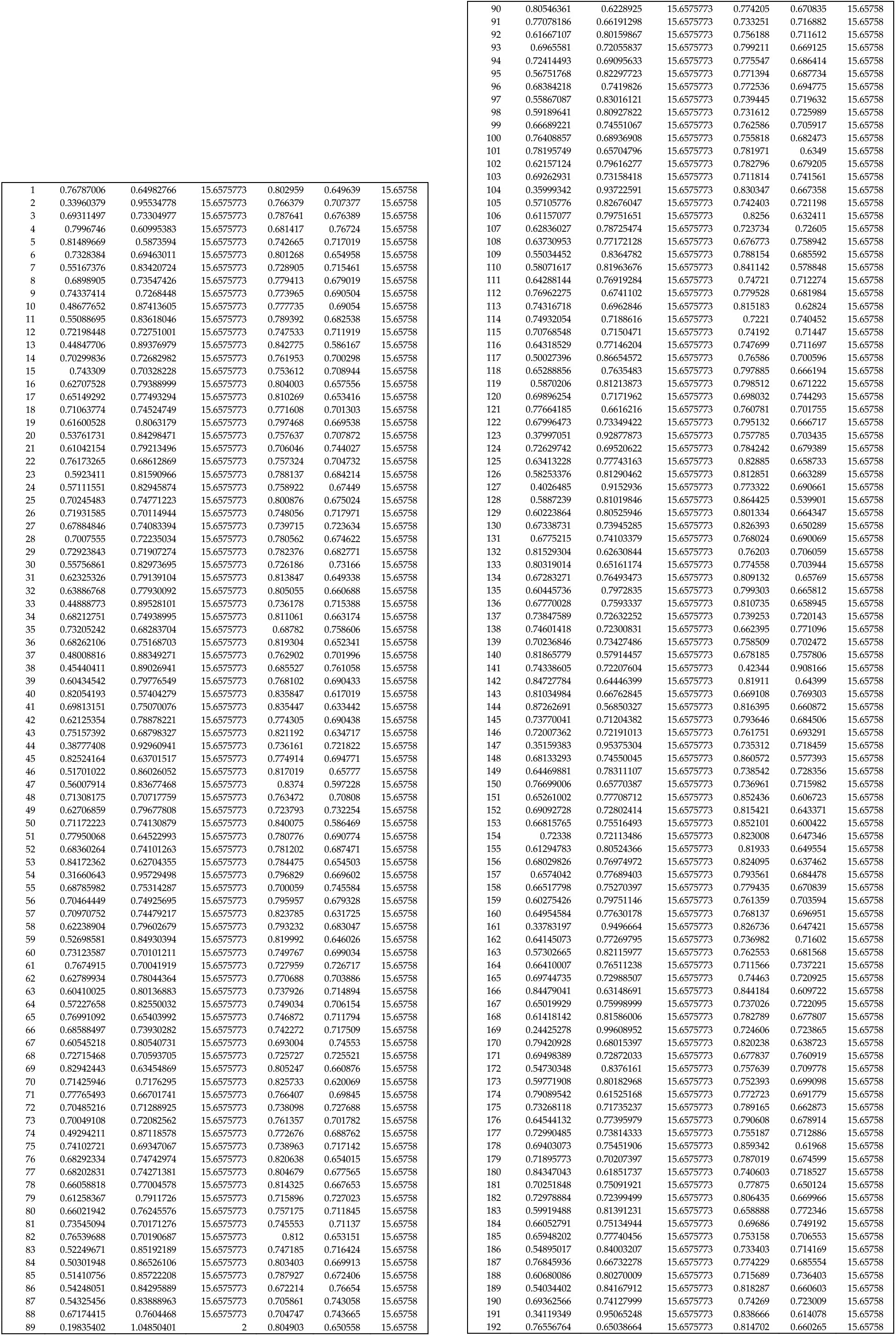
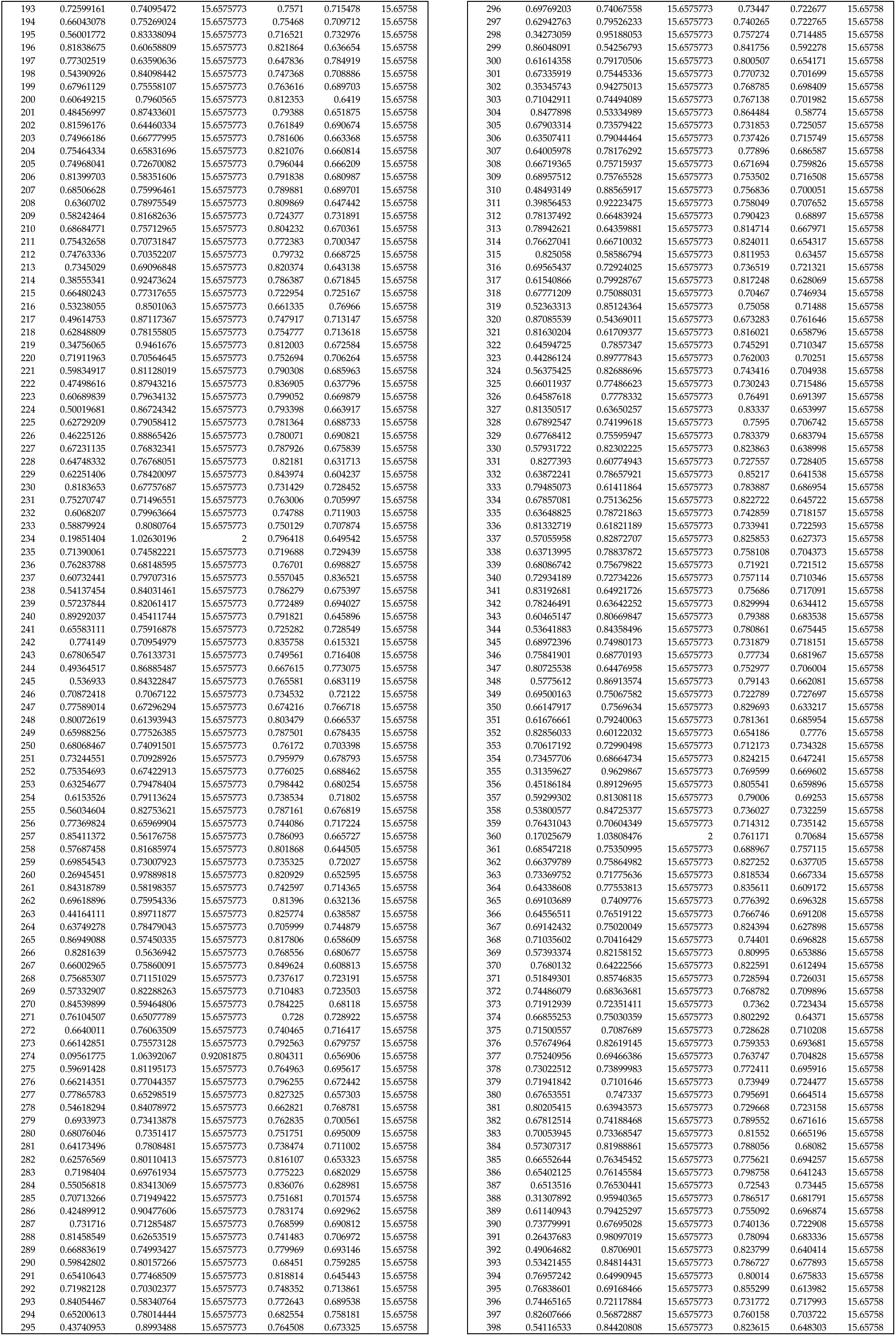
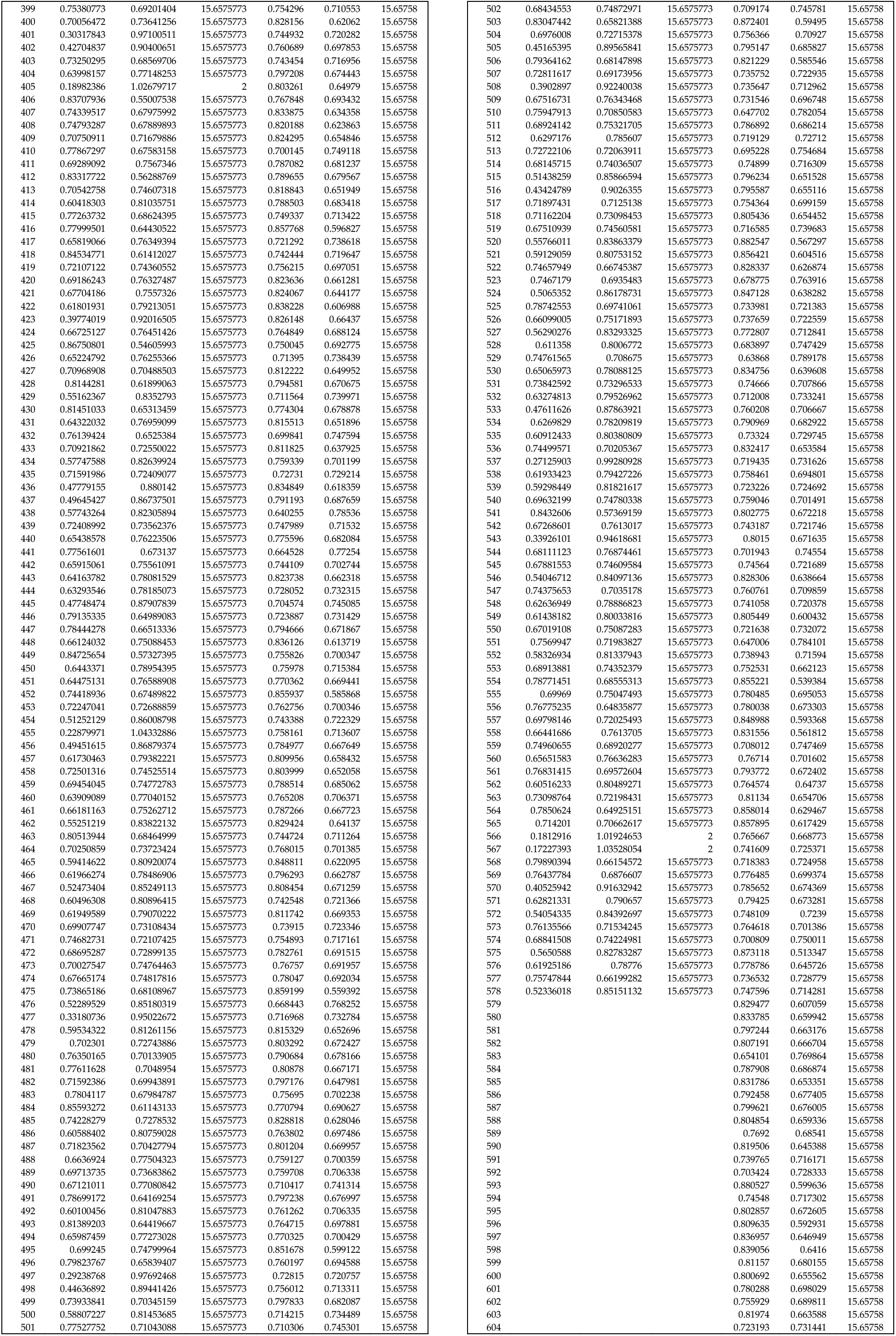
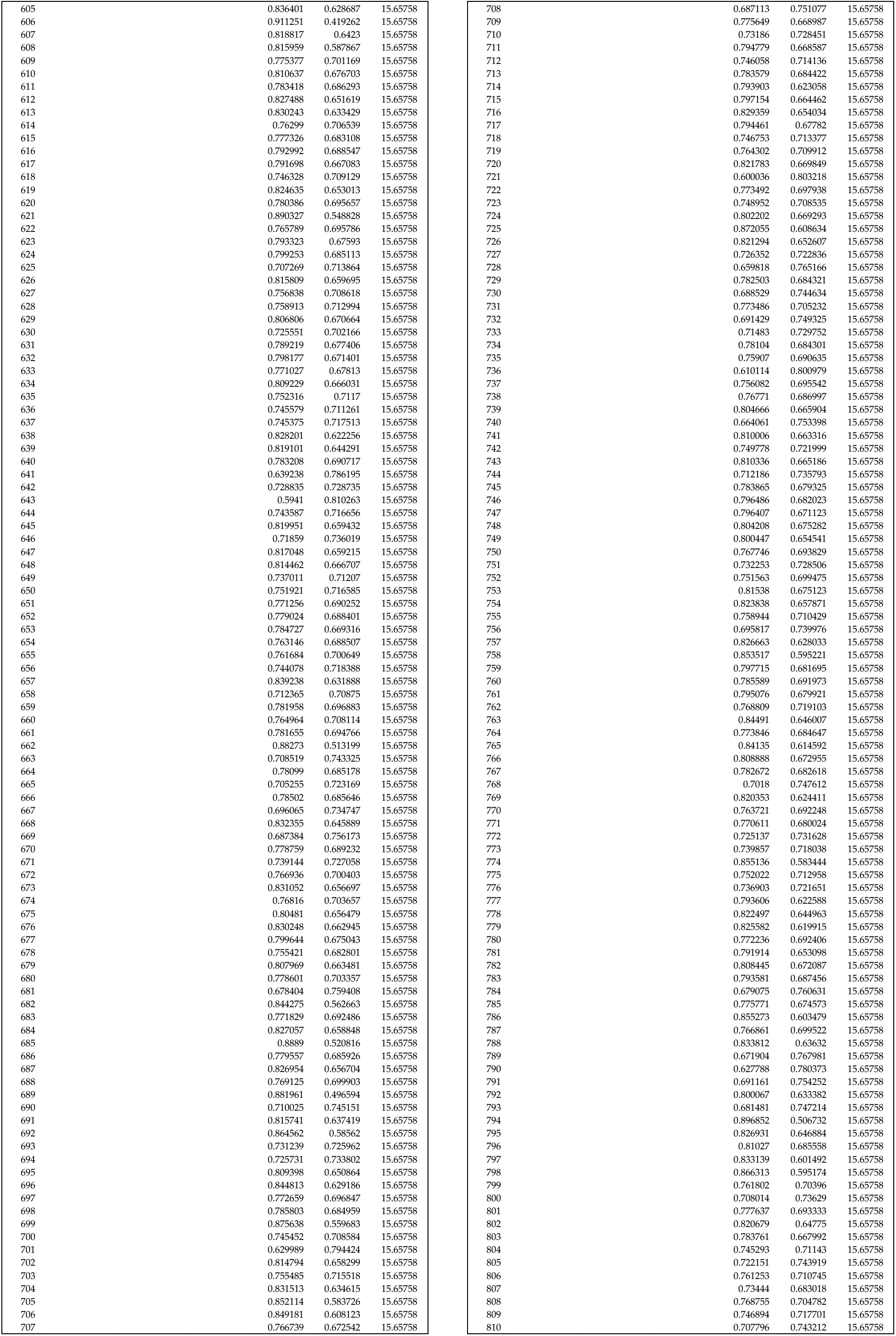
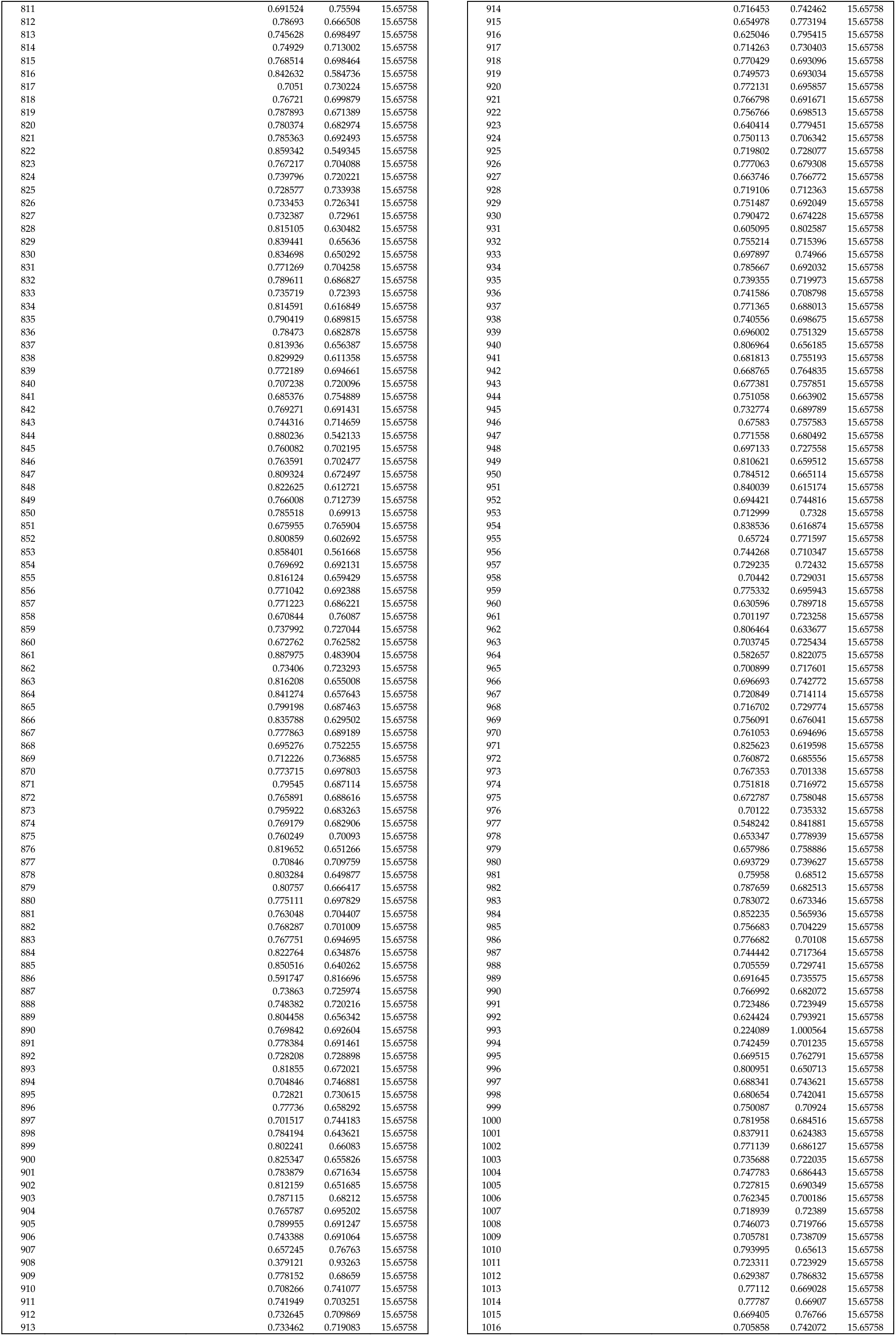
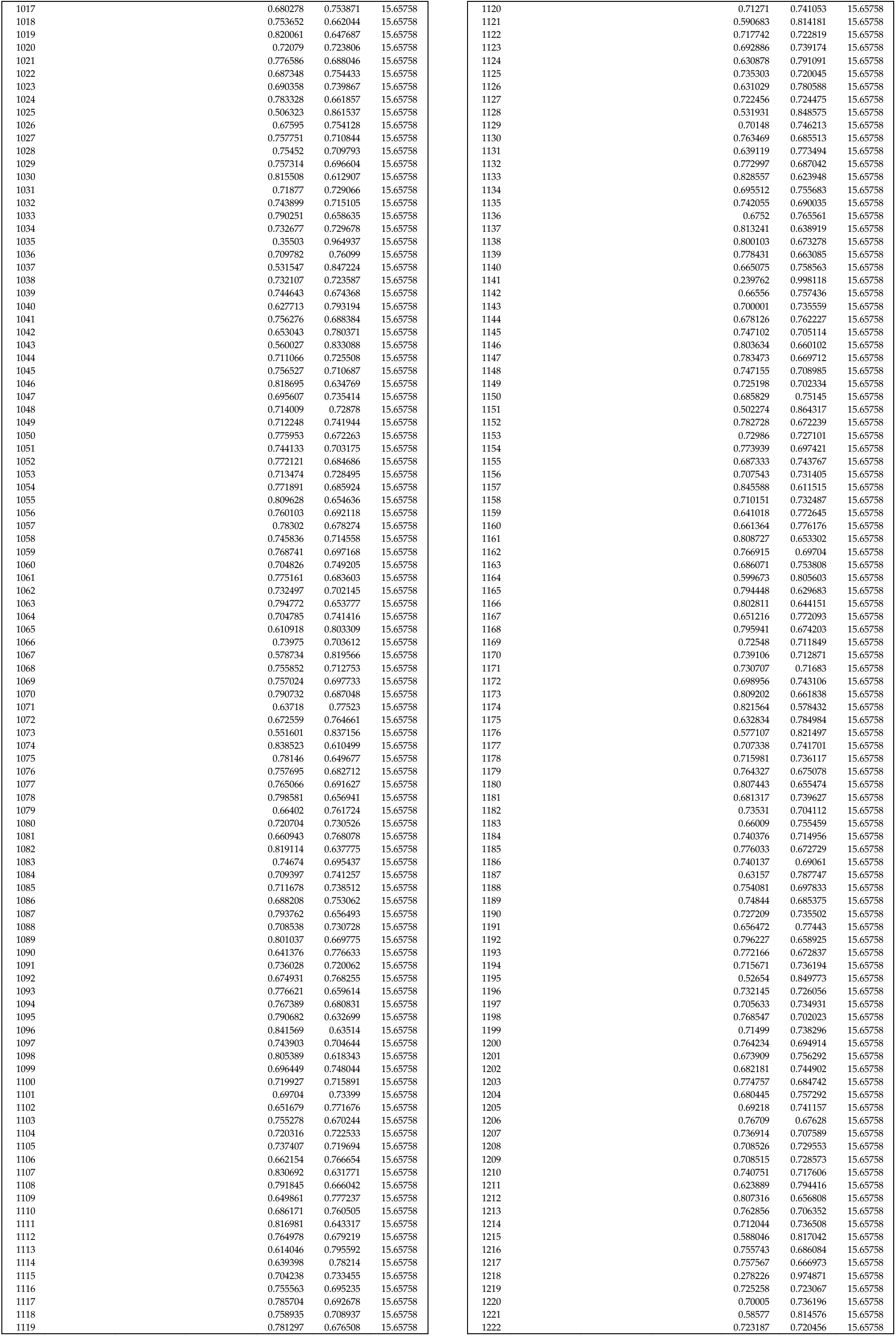
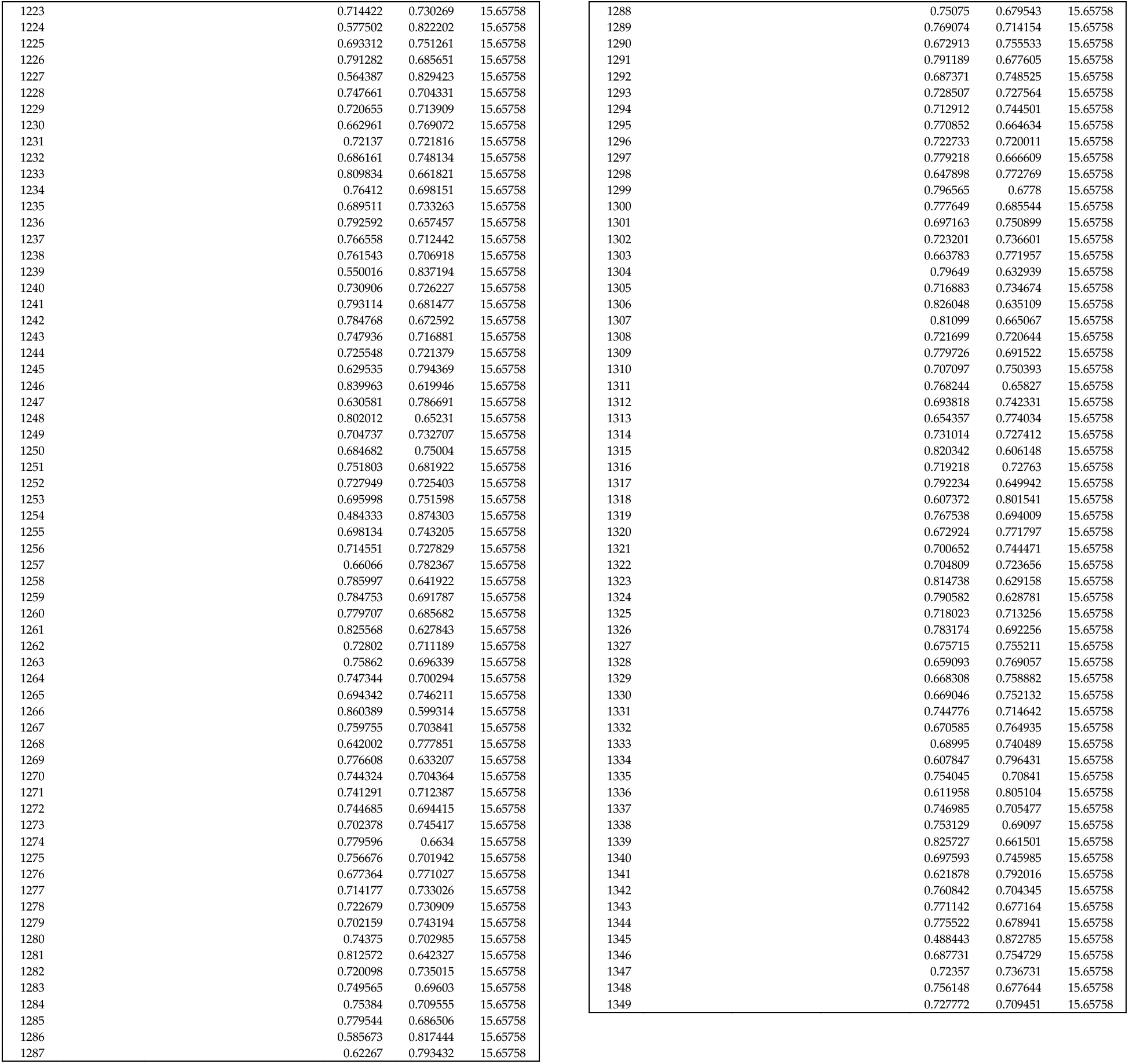
Source data of *in silico* cytometry statistics. For each sample in the GTEx (n=578) and LUG (n=1,349) datasets, Pearson correlation coefficient *R*, root mean square error (RMSE) and −log_10_(*p* value) are shown. These are the source data of Figure S1.

**Table S2.**
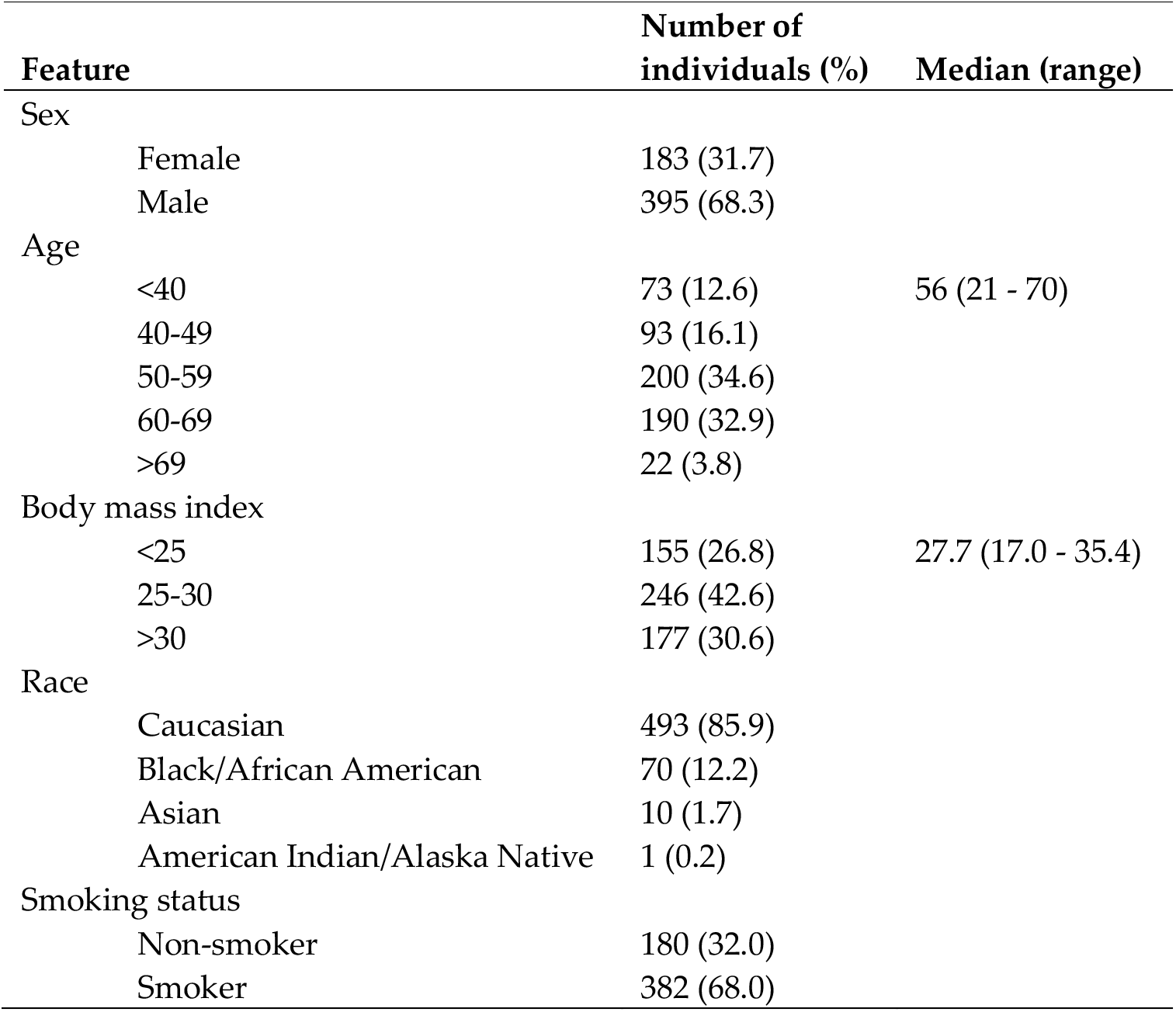
Phenotypic characteristics of GTEx lung tissue donors.

**Table S3.**
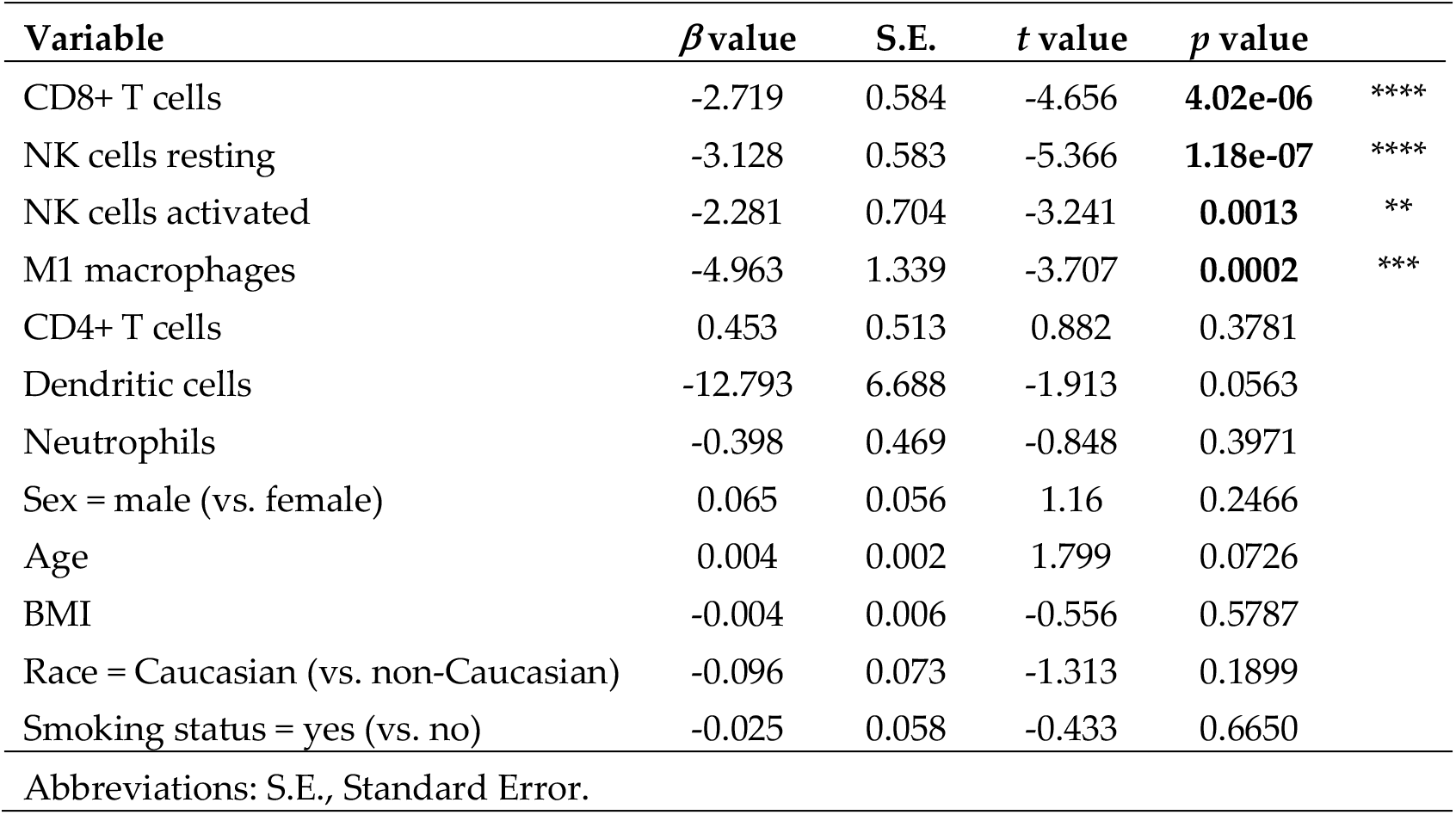
Multivariate analysis of ACE2 expression in human lung tissue.

**Table S4.**
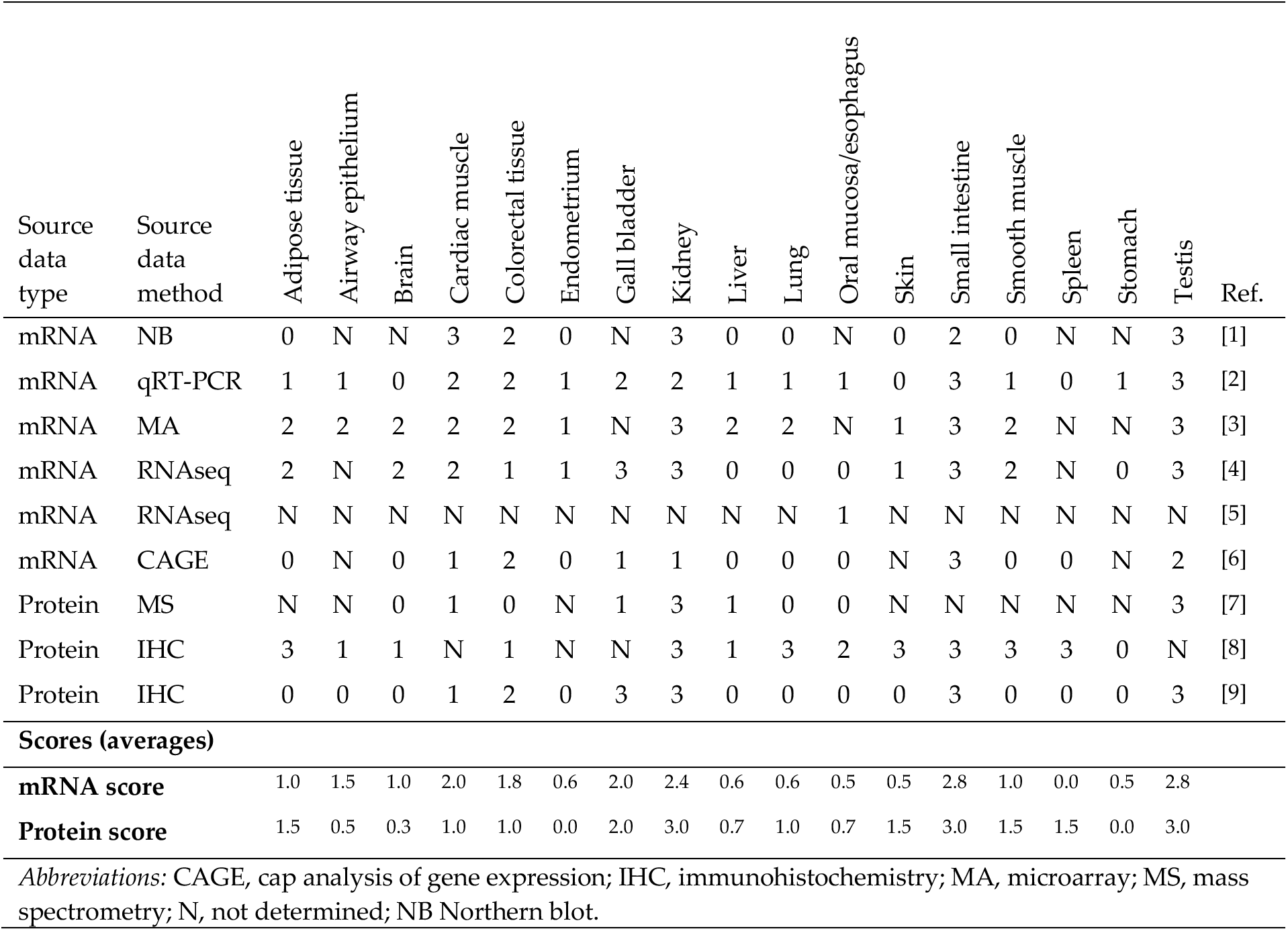
Meta-analysis of ACE2 mRNA and protein levels in 17 human tissues. *Source data of Fig. 2f.*

## Notes

https://gtexportal.org

https://www.ncbi.nlm.nih.gov/geo

### Competing Interest Statement

The authors have declared no competing interest.

