## Supplementary Information for "Low baseline pulmonary levels of cytotoxic lymphocytes as a predisposing risk factor for severe COVID-19"

Pascal H.G. Duijf

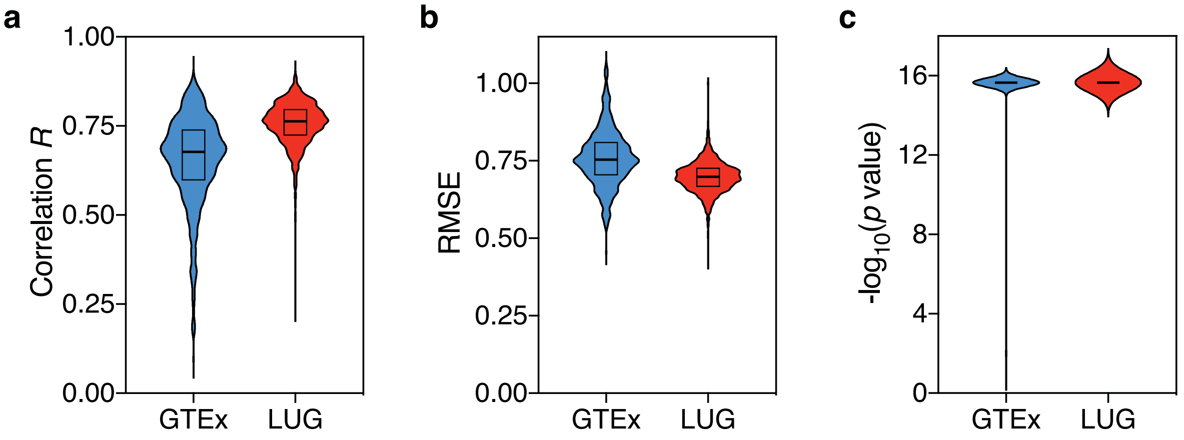


**Figure S1. *In silico* cytometry statistics.**
Per-sample statistics of *in silico* cytometry on human lung tissues are shown for the GTEx (n=578) and LUG (n=1,349) datasets. **(a)** Pearson correlation coefficients *R*. **(b)** Root mean square errors (RMSE). **(c)** *p* values. *P* values < $2.2\times10^{-16}$ were processed as equal to $2.2\times10^{-16}$. Source data are provided in Table S1.

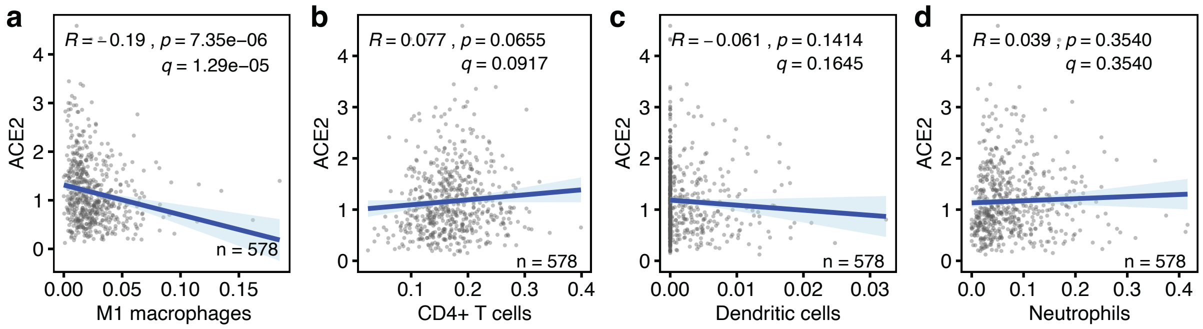


**Figure S2. Correlations between baseline anti-viral leukocyte levels and ACE2 expression levels in human lung tissues.**
**(a-d)** The correlations between the baseline levels of indicated immune cell types (x-axes) and the expression level of the SARS-CoV-2 host cell receptor ACE2 (y-axis) in human lung tissue are shown. Data are from the GTEx dataset (n=578). Regression lines and 95% confidence intervals are shown. *R* and *p* values: Pearson correlations; *q* values: Benjamini-Hochberg-adjusted *p* values using a false discovery rate of 0.05. See also Figure 1.

**Figure S3. Correlation between baseline M1 macrophage levels and ACE2 expression level in human lung tissue.** The correlation between these variables is shown as described in Figure S2. Data are from the LUG dataset (n=1,349).

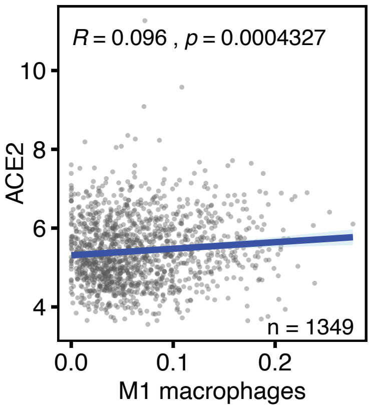


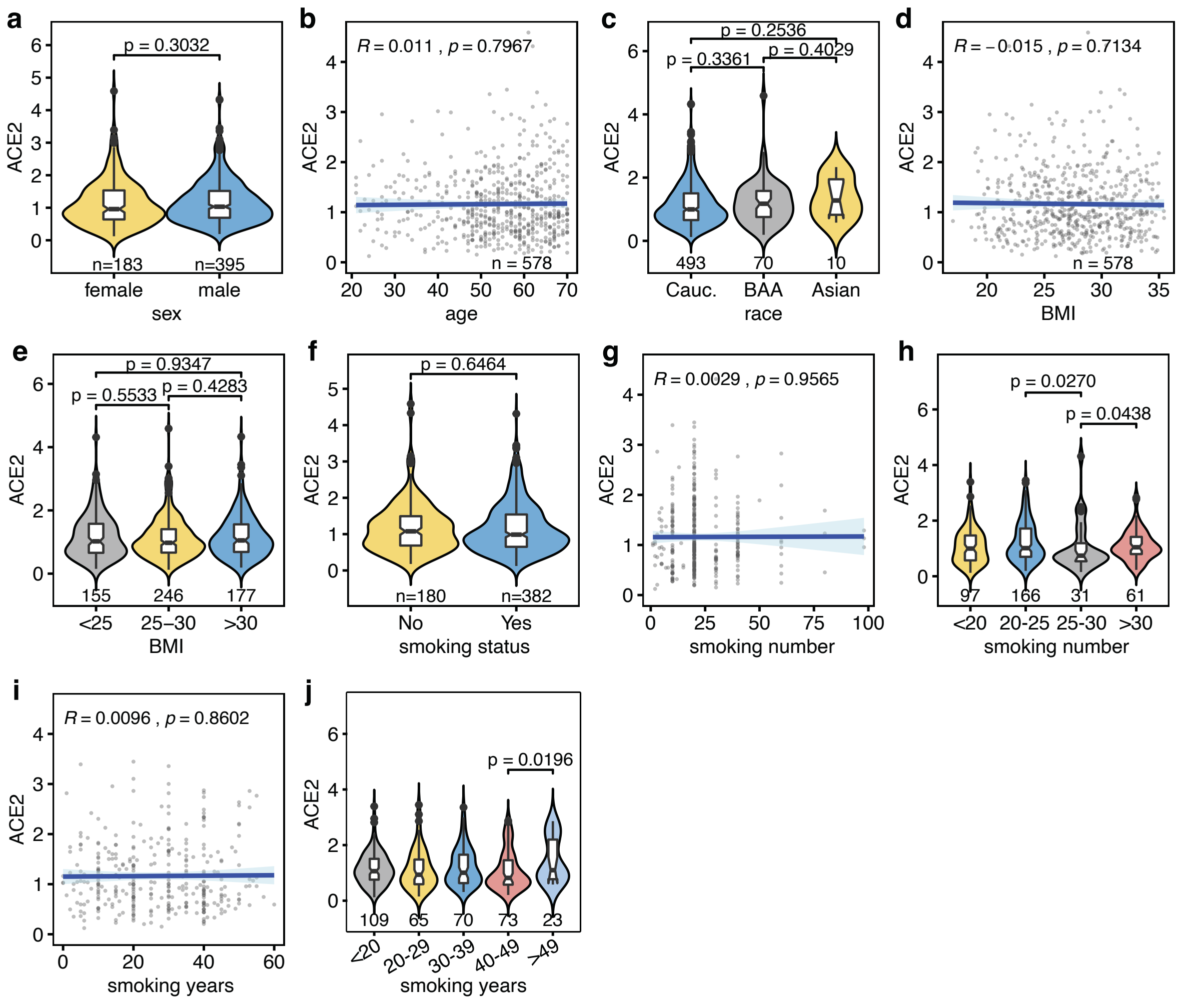


**Figure S4. Univariate analyses of phenotypic covariates included in multivariate analyses.** Univariate analyses of five covariates that were included in multivariate analyses of ACE2 and TMPRSS2 expression in human lung tissue using the GTEx dataset (Tables S2, S3). **(a)** Sex. **(b)** Age, **(c)** Race. **(d, e)** Body mass index. **(f-j)** Smoking behavior, referring to smoking status (smoker/non-smoker) (f), number of units smoked during the smoke period (g, h) and number of years smoked (i, j). *P* values in categorical analyses: Mann-Whitney *U* tests. *R* and *p* values in continuous analyses: Pearson correlations. *P* values in panels h and j are only shown if $p<0.05$. Sample numbers (n) are shown on the x-axes.

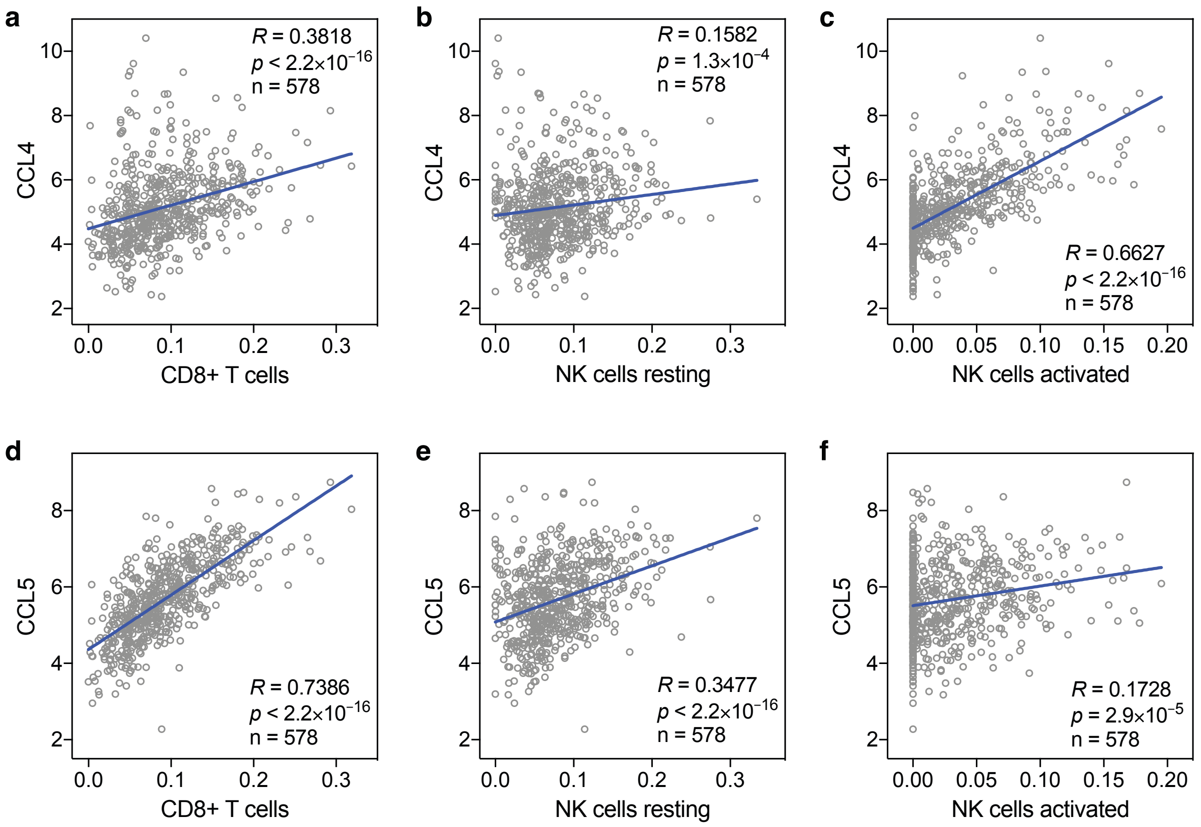


**Figure S5. Correlations between baseline levels of cytotoxic lymphocytes and CCL4 and CCL5 in human lung tissue.**

The correlations between baseline levels of the chemokines CCL4, CCL5 and CD8+ T cells, resting and activated NK cells in human lung tissue are shown. Data are from the GTEx dataset (n=578). *R* and *p* values: Spearman correlations. See also Figure 2g.

### Table S1. Source data of *in silico* cytometry statistics

For each sample in the GTEx (n=578) and LUG (n=1,349) datasets, Pearson correlation coefficient *R*, root mean square error (RMSE) and -log10(*p* value) are shown. These are the source data of Figure S1.

| Sample n | GTEx R | GTEx RMSE | GTEx -log10 pval | LUG R | LUG RMSE | LUG -log10 pval |
|---|---|---|---|---|---|---|
| 1 | 0.76787006 | 0.64982766 | 15.6575773 | 0.802959 | 0.649639 | 15.65758 |
| 2 | 0.33960379 | 0.95534778 | 15.6575773 | 0.766379 | 0.707377 | 15.65758 |
| 3 | 0.69311497 | 0.73304977 | 15.6575773 | 0.787641 | 0.676389 | 15.65758 |
| 4 | 0.7996746 | 0.60995383 | 15.6575773 | 0.681417 | 0.76724 | 15.65758 |
| 5 | 0.81489669 | 0.5873594 | 15.6575773 | 0.742665 | 0.717019 | 15.65758 |
| 6 | 0.7328384 | 0.69463011 | 15.6575773 | 0.801268 | 0.654958 | 15.65758 |
| 7 | 0.55167376 | 0.83420724 | 15.6575773 | 0.728905 | 0.715461 | 15.65758 |
| 8 | 0.6898905 | 0.73547426 | 15.6575773 | 0.779413 | 0.679019 | 15.65758 |
| 9 | 0.74337414 | 0.7268448 | 15.6575773 | 0.773965 | 0.690504 | 15.65758 |
| 10 | 0.48677652 | 0.87413605 | 15.6575773 | 0.777735 | 0.69054 | 15.65758 |
| 11 | 0.55088695 | 0.83618046 | 15.6575773 | 0.789392 | 0.682538 | 15.65758 |
| 12 | 0.72198448 | 0.72751001 | 15.6575773 | 0.747533 | 0.711919 | 15.65758 |
| 13 | 0.44847706 | 0.89376979 | 15.6575773 | 0.842775 | 0.586167 | 15.65758 |
| 14 | 0.70299836 | 0.72682982 | 15.6575773 | 0.761953 | 0.700298 | 15.65758 |
| 15 | 0.743309 | 0.70328228 | 15.6575773 | 0.753612 | 0.708944 | 15.65758 |
| 16 | 0.62707528 | 0.79388999 | 15.6575773 | 0.804003 | 0.657556 | 15.65758 |
| 17 | 0.65149292 | 0.77493294 | 15.6575773 | 0.810269 | 0.653416 | 15.65758 |
| 18 | 0.71063774 | 0.74524749 | 15.6575773 | 0.771608 | 0.701303 | 15.65758 |
| 19 | 0.61600528 | 0.8063179 | 15.6575773 | 0.797468 | 0.669538 | 15.65758 |
| 20 | 0.53761731 | 0.84298471 | 15.6575773 | 0.757637 | 0.707872 | 15.65758 |
| 21 | 0.61042154 | 0.79213496 | 15.6575773 | 0.706046 | 0.744027 | 15.65758 |
| 22 | 0.76173265 | 0.68612869 | 15.6575773 | 0.757324 | 0.704732 | 15.65758 |
| 23 | 0.5923411 | 0.81590966 | 15.6575773 | 0.788137 | 0.684214 | 15.65758 |
| 24 | 0.57111551 | 0.82945874 | 15.6575773 | 0.758922 | 0.67449 | 15.65758 |
| 25 | 0.70245483 | 0.74771223 | 15.6575773 | 0.800876 | 0.675024 | 15.65758 |
| 26 | 0.71931585 | 0.70114944 | 15.6575773 | 0.748056 | 0.717971 | 15.65758 |
| 27 | 0.67884846 | 0.74083394 | 15.6575773 | 0.739715 | 0.723634 | 15.65758 |
| 28 | 0.7007555 | 0.72235034 | 15.6575773 | 0.780562 | 0.674622 | 15.65758 |
| 29 | 0.72923843 | 0.71907274 | 15.6575773 | 0.782376 | 0.682771 | 15.65758 |
| 30 | 0.55756861 | 0.82973695 | 15.6575773 | 0.726186 | 0.73166 | 15.65758 |
| 31 | 0.62325326 | 0.79139104 | 15.6575773 | 0.813847 | 0.649338 | 15.65758 |
| 32 | 0.63886768 | 0.77930092 | 15.6575773 | 0.805055 | 0.660688 | 15.65758 |
| 33 | 0.44888773 | 0.89528101 | 15.6575773 | 0.736178 | 0.715388 | 15.65758 |
| 34 | 0.68212751 | 0.74938995 | 15.6575773 | 0.811061 | 0.663174 | 15.65758 |
| 35 | 0.73205242 | 0.68283704 | 15.6575773 | 0.68782 | 0.758606 | 15.65758 |
| 36 | 0.68262106 | 0.75168703 | 15.6575773 | 0.819304 | 0.652341 | 15.65758 |
| 37 | 0.48008816 | 0.88349271 | 15.6575773 | 0.762902 | 0.701996 | 15.65758 |
| 38 | 0.45440411 | 0.89026941 | 15.6575773 | 0.685527 | 0.761058 | 15.65758 |
| 39 | 0.60434542 | 0.79776549 | 15.6575773 | 0.768102 | 0.690433 | 15.65758 |
| 40 | 0.82054193 | 0.57404279 | 15.6575773 | 0.835847 | 0.617019 | 15.65758 |
| 41 | 0.69813151 | 0.75070076 | 15.6575773 | 0.835447 | 0.633442 | 15.65758 |
| 42 | 0.62125354 | 0.78878221 | 15.6575773 | 0.774305 | 0.690438 | 15.65758 |
| 43 | 0.75157392 | 0.68798327 | 15.6575773 | 0.821192 | 0.634717 | 15.65758 |
| 44 | 0.38777408 | 0.92960941 | 15.6575773 | 0.736161 | 0.721822 | 15.65758 |
| 45 | 0.82524164 | 0.63701517 | 15.6575773 | 0.774914 | 0.694771 | 15.65758 |
| 46 | 0.51701022 | 0.86026052 | 15.6575773 | 0.817019 | 0.65777 | 15.65758 |
| 47 | 0.56007914 | 0.83677468 | 15.6575773 | 0.8374 | 0.597228 | 15.65758 |
| 48 | 0.71308175 | 0.70717759 | 15.6575773 | 0.763472 | 0.70808 | 15.65758 |
| 49 | 0.62706859 | 0.79677808 | 15.6575773 | 0.723793 | 0.732254 | 15.65758 |
| 50 | 0.71172223 | 0.74130879 | 15.6575773 | 0.840075 | 0.586469 | 15.65758 |
| 51 | 0.77950068 | 0.64522993 | 15.6575773 | 0.780776 | 0.690774 | 15.65758 |
| 52 | 0.68360264 | 0.74101263 | 15.6575773 | 0.781202 | 0.687471 | 15.65758 |
| 53 | 0.84172362 | 0.62704355 | 15.6575773 | 0.784475 | 0.654503 | 15.65758 |
| 54 | 0.31660643 | 0.95729498 | 15.6575773 | 0.796829 | 0.669602 | 15.65758 |
| 55 | 0.68785982 | 0.75314287 | 15.6575773 | 0.700059 | 0.745584 | 15.65758 |
| 56 | 0.70464449 | 0.74925695 | 15.6575773 | 0.795957 | 0.679328 | 15.65758 |
| 57 | 0.70970752 | 0.74479217 | 15.6575773 | 0.823785 | 0.631725 | 15.65758 |
| 58 | 0.62238904 | 0.79602679 | 15.6575773 | 0.793232 | 0.683047 | 15.65758 |
| 59 | 0.52698581 | 0.84930394 | 15.6575773 | 0.819992 | 0.646026 | 15.65758 |
| 60 | 0.73123587 | 0.70101211 | 15.6575773 | 0.749767 | 0.699034 | 15.65758 |
| 61 | 0.7674915 | 0.70041919 | 15.6575773 | 0.727959 | 0.726717 | 15.65758 |
| 62 | 0.62789934 | 0.78044364 | 15.6575773 | 0.770688 | 0.703886 | 15.65758 |
| 63 | 0.60410025 | 0.80136883 | 15.6575773 | 0.737926 | 0.714894 | 15.65758 |
| 64 | 0.57227658 | 0.82550032 | 15.6575773 | 0.749034 | 0.706154 | 15.65758 |
| 65 | 0.76991092 | 0.65403992 | 15.6575773 | 0.746872 | 0.711794 | 15.65758 |
| 66 | 0.68588497 | 0.73930282 | 15.6575773 | 0.742272 | 0.717509 | 15.65758 |
| 67 | 0.60545218 | 0.80540731 | 15.6575773 | 0.693004 | 0.74553 | 15.65758 |
| 68 | 0.72715468 | 0.70593705 | 15.6575773 | 0.725727 | 0.725521 | 15.65758 |
| 69 | 0.82942443 | 0.63454869 | 15.6575773 | 0.805247 | 0.660876 | 15.65758 |
| 70 | 0.71425946 | 0.7176295 | 15.6575773 | 0.825733 | 0.620069 | 15.65758 |
| 71 | 0.77765493 | 0.66701741 | 15.6575773 | 0.766407 | 0.69845 | 15.65758 |
| 72 | 0.70485216 | 0.71288925 | 15.6575773 | 0.738098 | 0.727688 | 15.65758 |
| 73 | 0.70049108 | 0.72082562 | 15.6575773 | 0.761357 | 0.701782 | 15.65758 |
| 74 | 0.49294211 | 0.87118578 | 15.6575773 | 0.772676 | 0.688762 | 15.65758 |
| 75 | 0.74102721 | 0.69347067 | 15.6575773 | 0.738963 | 0.717142 | 15.65758 |
| 76 | 0.68292334 | 0.74742974 | 15.6575773 | 0.820638 | 0.654015 | 15.65758 |
| 77 | 0.68202831 | 0.74271381 | 15.6575773 | 0.804679 | 0.677565 | 15.65758 |
| 78 | 0.66058818 | 0.77004578 | 15.6575773 | 0.814325 | 0.667653 | 15.65758 |
| 79 | 0.61258367 | 0.7911726 | 15.6575773 | 0.715896 | 0.727023 | 15.65758 |
| 80 | 0.66021942 | 0.76245576 | 15.6575773 | 0.757175 | 0.711845 | 15.65758 |
| 81 | 0.73545094 | 0.70171276 | 15.6575773 | 0.745553 | 0.71137 | 15.65758 |
| 82 | 0.76539688 | 0.70190687 | 15.6575773 | 0.812 | 0.653151 | 15.65758 |
| 83 | 0.52249671 | 0.85192189 | 15.6575773 | 0.747185 | 0.716424 | 15.65758 |
| 84 | 0.50301948 | 0.86526106 | 15.6575773 | 0.803403 | 0.669913 | 15.65758 |
| 85 | 0.51410756 | 0.85722208 | 15.6575773 | 0.787927 | 0.672406 | 15.65758 |
| 86 | 0.54248051 | 0.84295889 | 15.6575773 | 0.672214 | 0.76654 | 15.65758 |
| 87 | 0.54325456 | 0.83888963 | 15.6575773 | 0.705861 | 0.743058 | 15.65758 |
| 88 | 0.67174415 | 0.7604468 | 15.6575773 | 0.704747 | 0.743665 | 15.65758 |
| 89 | 0.19835402 | 1.04850401 | 2 | 0.804903 | 0.650558 | 15.65758 |
| 90 | 0.80546361 | 0.6228925 | 15.6575773 | 0.774205 | 0.670835 | 15.65758 |
| 91 | 0.77078186 | 0.66191298 | 15.6575773 | 0.733251 | 0.716882 | 15.65758 |
| 92 | 0.61667107 | 0.80159867 | 15.6575773 | 0.756188 | 0.711612 | 15.65758 |
| 93 | 0.6965581 | 0.72055837 | 15.6575773 | 0.799211 | 0.669125 | 15.65758 |
| 94 | 0.72414493 | 0.69095633 | 15.6575773 | 0.775547 | 0.686414 | 15.65758 |
| 95 | 0.56751768 | 0.82297723 | 15.6575773 | 0.771394 | 0.687734 | 15.65758 |
| 96 | 0.68384218 | 0.7419826 | 15.6575773 | 0.772536 | 0.694775 | 15.65758 |
| 97 | 0.55867087 | 0.83016121 | 15.6575773 | 0.739445 | 0.719632 | 15.65758 |
| 98 | 0.59189641 | 0.80927822 | 15.6575773 | 0.731612 | 0.725989 | 15.65758 |
| 99 | 0.66689221 | 0.74551067 | 15.6575773 | 0.762586 | 0.705917 | 15.65758 |
| 100 | 0.76408857 | 0.68936908 | 15.6575773 | 0.755818 | 0.682473 | 15.65758 |
| 101 | 0.78195749 | 0.65704796 | 15.6575773 | 0.781971 | 0.6349 | 15.65758 |
| 102 | 0.62157124 | 0.79616277 | 15.6575773 | 0.782796 | 0.679205 | 15.65758 |
| 103 | 0.69262931 | 0.73158418 | 15.6575773 | 0.711814 | 0.741561 | 15.65758 |
| 104 | 0.35999342 | 0.93722591 | 15.6575773 | 0.830347 | 0.667358 | 15.65758 |
| 105 | 0.57105776 | 0.82676047 | 15.6575773 | 0.742403 | 0.721198 | 15.65758 |
| 106 | 0.61157077 | 0.79751651 | 15.6575773 | 0.8256 | 0.632411 | 15.65758 |
| 107 | 0.62836027 | 0.78725474 | 15.6575773 | 0.723734 | 0.72605 | 15.65758 |
| 108 | 0.63730953 | 0.77172128 | 15.6575773 | 0.676773 | 0.758942 | 15.65758 |
| 109 | 0.55034452 | 0.8364782 | 15.6575773 | 0.788154 | 0.685592 | 15.65758 |
| 110 | 0.58071617 | 0.81963676 | 15.6575773 | 0.841142 | 0.578848 | 15.65758 |
| 111 | 0.64288144 | 0.76919284 | 15.6575773 | 0.74721 | 0.712274 | 15.65758 |
| 112 | 0.76962275 | 0.6741102 | 15.6575773 | 0.779528 | 0.681984 | 15.65758 |
| 113 | 0.74316718 | 0.6962846 | 15.6575773 | 0.815183 | 0.62824 | 15.65758 |
| 114 | 0.74932054 | 0.7188616 | 15.6575773 | 0.7221 | 0.740452 | 15.65758 |
| 115 | 0.70768548 | 0.7150471 | 15.6575773 | 0.74192 | 0.71447 | 15.65758 |
| 116 | 0.64318529 | 0.77146204 | 15.6575773 | 0.747699 | 0.711697 | 15.65758 |
| 117 | 0.50027396 | 0.86654572 | 15.6575773 | 0.76586 | 0.700596 | 15.65758 |
| 118 | 0.65288856 | 0.7635483 | 15.6575773 | 0.797885 | 0.666194 | 15.65758 |
| 119 | 0.5870206 | 0.81213873 | 15.6575773 | 0.798512 | 0.671222 | 15.65758 |
| 120 | 0.69896254 | 0.7171962 | 15.6575773 | 0.698032 | 0.744293 | 15.65758 |
| 121 | 0.77664185 | 0.6616216 | 15.6575773 | 0.760781 | 0.701755 | 15.65758 |
| 122 | 0.67996473 | 0.73349422 | 15.6575773 | 0.795132 | 0.666717 | 15.65758 |
| 123 | 0.37997051 | 0.92877873 | 15.6575773 | 0.757785 | 0.703435 | 15.65758 |
| 124 | 0.72629742 | 0.69520622 | 15.6575773 | 0.784242 | 0.679389 | 15.65758 |
| 125 | 0.63413228 | 0.77743163 | 15.6575773 | 0.82885 | 0.658733 | 15.65758 |
| 126 | 0.58253376 | 0.81290462 | 15.6575773 | 0.812851 | 0.663289 | 15.65758 |
| 127 | 0.4026485 | 0.9152936 | 15.6575773 | 0.773322 | 0.690661 | 15.65758 |
| 128 | 0.5887239 | 0.81019846 | 15.6575773 | 0.864425 | 0.539901 | 15.65758 |
| 129 | 0.60223864 | 0.80525946 | 15.6575773 | 0.801334 | 0.664347 | 15.65758 |
| 130 | 0.67338731 | 0.73945285 | 15.6575773 | 0.826393 | 0.650289 | 15.65758 |
| 131 | 0.6775215 | 0.74103379 | 15.6575773 | 0.768024 | 0.690069 | 15.65758 |
| 132 | 0.81529304 | 0.62630844 | 15.6575773 | 0.76203 | 0.706059 | 15.65758 |
| 133 | 0.80319014 | 0.65161174 | 15.6575773 | 0.774558 | 0.703944 | 15.65758 |
| 134 | 0.67283271 | 0.76493473 | 15.6575773 | 0.809132 | 0.65769 | 15.65758 |
| 135 | 0.60445736 | 0.7972835 | 15.6575773 | 0.799303 | 0.665812 | 15.65758 |
| 136 | 0.67770028 | 0.7593337 | 15.6575773 | 0.810735 | 0.658945 | 15.65758 |
| 137 | 0.73847589 | 0.72632252 | 15.6575773 | 0.739253 | 0.720143 | 15.65758 |
| 138 | 0.74601418 | 0.72300831 | 15.6575773 | 0.662395 | 0.771096 | 15.65758 |
| 139 | 0.70236846 | 0.73427486 | 15.6575773 | 0.758509 | 0.702472 | 15.65758 |
| 140 | 0.81865779 | 0.57914457 | 15.6575773 | 0.678185 | 0.757806 | 15.65758 |
| 141 | 0.74338605 | 0.72207604 | 15.6575773 | 0.42344 | 0.908166 | 15.65758 |
| 142 | 0.84727784 | 0.64446399 | 15.6575773 | 0.81911 | 0.64399 | 15.65758 |
| 143 | 0.81034984 | 0.66762845 | 15.6575773 | 0.669108 | 0.769303 | 15.65758 |
| 144 | 0.87262691 | 0.56850327 | 15.6575773 | 0.816395 | 0.660872 | 15.65758 |
| 145 | 0.73770041 | 0.71204382 | 15.6575773 | 0.793646 | 0.684506 | 15.65758 |
| 146 | 0.72007362 | 0.72191013 | 15.6575773 | 0.761751 | 0.693291 | 15.65758 |
| 147 | 0.35159383 | 0.95375304 | 15.6575773 | 0.735312 | 0.718459 | 15.65758 |
| 148 | 0.68133293 | 0.74550045 | 15.6575773 | 0.860572 | 0.577393 | 15.65758 |
| 149 | 0.64469881 | 0.78311107 | 15.6575773 | 0.738542 | 0.728356 | 15.65758 |
| 150 | 0.76699006 | 0.65770387 | 15.6575773 | 0.736961 | 0.715982 | 15.65758 |
| 151 | 0.65261002 | 0.77708712 | 15.6575773 | 0.852436 | 0.606723 | 15.65758 |
| 152 | 0.69092728 | 0.72802414 | 15.6575773 | 0.815421 | 0.643371 | 15.65758 |
| 153 | 0.66815765 | 0.75516493 | 15.6575773 | 0.852101 | 0.600422 | 15.65758 |
| 154 | 0.72338 | 0.72113486 | 15.6575773 | 0.823008 | 0.647346 | 15.65758 |
| 155 | 0.61294783 | 0.80524366 | 15.6575773 | 0.81933 | 0.649554 | 15.65758 |
| 156 | 0.68029826 | 0.76974972 | 15.6575773 | 0.824095 | 0.637462 | 15.65758 |
| 157 | 0.6574042 | 0.77689403 | 15.6575773 | 0.793561 | 0.684478 | 15.65758 |
| 158 | 0.66517798 | 0.75270397 | 15.6575773 | 0.779435 | 0.670839 | 15.65758 |
| 159 | 0.60275426 | 0.79751146 | 15.6575773 | 0.761359 | 0.703594 | 15.65758 |
| 160 | 0.64954584 | 0.77630178 | 15.6575773 | 0.768137 | 0.696951 | 15.65758 |
| 161 | 0.33783197 | 0.9496664 | 15.6575773 | 0.826736 | 0.647421 | 15.65758 |
| 162 | 0.64145073 | 0.77269795 | 15.6575773 | 0.736982 | 0.71602 | 15.65758 |
| 163 | 0.57302665 | 0.82115977 | 15.6575773 | 0.762553 | 0.681568 | 15.65758 |
| 164 | 0.66410007 | 0.76511238 | 15.6575773 | 0.711566 | 0.737221 | 15.65758 |
| 165 | 0.69744735 | 0.72988507 | 15.6575773 | 0.74463 | 0.720925 | 15.65758 |
| 166 | 0.84479041 | 0.63148691 | 15.6575773 | 0.844184 | 0.609722 | 15.65758 |
| 167 | 0.65019929 | 0.75998999 | 15.6575773 | 0.737026 | 0.722095 | 15.65758 |
| 168 | 0.61418142 | 0.81586006 | 15.6575773 | 0.782789 | 0.677807 | 15.65758 |
| 169 | 0.24425278 | 0.99608952 | 15.6575773 | 0.724606 | 0.723865 | 15.65758 |
| 170 | 0.79420928 | 0.68015397 | 15.6575773 | 0.820238 | 0.638723 | 15.65758 |
| 171 | 0.69498389 | 0.72872033 | 15.6575773 | 0.677837 | 0.760919 | 15.65758 |
| 172 | 0.54730348 | 0.8376161 | 15.6575773 | 0.757639 | 0.709778 | 15.65758 |
| 173 | 0.59771908 | 0.80182968 | 15.6575773 | 0.752393 | 0.699098 | 15.65758 |
| 174 | 0.79089542 | 0.61525168 | 15.6575773 | 0.772723 | 0.691779 | 15.65758 |
| 175 | 0.73268118 | 0.71735237 | 15.6575773 | 0.789165 | 0.662873 | 15.65758 |
| 176 | 0.64544132 | 0.77395979 | 15.6575773 | 0.790608 | 0.678914 | 15.65758 |
| 177 | 0.72990485 | 0.73814333 | 15.6575773 | 0.755187 | 0.712886 | 15.65758 |
| 178 | 0.69403073 | 0.75451906 | 15.6575773 | 0.859342 | 0.61968 | 15.65758 |
| 179 | 0.71895773 | 0.70207397 | 15.6575773 | 0.787019 | 0.674599 | 15.65758 |
| 180 | 0.84347043 | 0.61851737 | 15.6575773 | 0.740603 | 0.718527 | 15.65758 |
| 181 | 0.70251848 | 0.75091921 | 15.6575773 | 0.77875 | 0.650124 | 15.65758 |
| 182 | 0.72978884 | 0.72399499 | 15.6575773 | 0.806435 | 0.669966 | 15.65758 |
| 183 | 0.59919488 | 0.81391231 | 15.6575773 | 0.658888 | 0.772346 | 15.65758 |
| 184 | 0.66052791 | 0.75134944 | 15.6575773 | 0.69686 | 0.749192 | 15.65758 |
| 185 | 0.65948202 | 0.77740456 | 15.6575773 | 0.753158 | 0.706553 | 15.65758 |
| 186 | 0.54895017 | 0.84003207 | 15.6575773 | 0.733403 | 0.714169 | 15.65758 |
| 187 | 0.76845936 | 0.66732278 | 15.6575773 | 0.774229 | 0.685554 | 15.65758 |
| 188 | 0.60680086 | 0.80270009 | 15.6575773 | 0.715689 | 0.736403 | 15.65758 |
| 189 | 0.54034402 | 0.84167912 | 15.6575773 | 0.818287 | 0.660603 | 15.65758 |
| 190 | 0.69362566 | 0.74127999 | 15.6575773 | 0.74269 | 0.723009 | 15.65758 |
| 191 | 0.34119349 | 0.95065248 | 15.6575773 | 0.838666 | 0.614078 | 15.65758 |
| 192 | 0.76556764 | 0.65038664 | 15.6575773 | 0.814702 | 0.660265 | 15.65758 |

| | | | | | | |
|---|---|---|---|---|---|---|
| 193 | 0.72599161 | 0.74095472 | 15.6575773 | 0.7571 | 0.715478 | 15.65758 |
| 194 | 0.66043078 | 0.75269024 | 15.6575773 | 0.75468 | 0.709712 | 15.65758 |
| 195 | 0.56001772 | 0.83338094 | 15.6575773 | 0.716521 | 0.732976 | 15.65758 |
| 196 | 0.81838675 | 0.60658809 | 15.6575773 | 0.821864 | 0.636654 | 15.65758 |
| 197 | 0.77302519 | 0.63590636 | 15.6575773 | 0.647836 | 0.784919 | 15.65758 |
| 198 | 0.54390926 | 0.84098442 | 15.6575773 | 0.747368 | 0.708886 | 15.65758 |
| 199 | 0.67961129 | 0.75558107 | 15.6575773 | 0.763616 | 0.689703 | 15.65758 |
| 200 | 0.60649215 | 0.7960565 | 15.6575773 | 0.812353 | 0.6419 | 15.65758 |
| 201 | 0.48456997 | 0.87433601 | 15.6575773 | 0.79388 | 0.651875 | 15.65758 |
| 202 | 0.81596176 | 0.64460334 | 15.6575773 | 0.761849 | 0.690674 | 15.65758 |
| 203 | 0.74966186 | 0.66777995 | 15.6575773 | 0.781606 | 0.663368 | 15.65758 |
| 204 | 0.75464334 | 0.65831696 | 15.6575773 | 0.821076 | 0.660814 | 15.65758 |
| 205 | 0.74968041 | 0.72670082 | 15.6575773 | 0.796044 | 0.666209 | 15.65758 |
| 206 | 0.81399703 | 0.58351606 | 15.6575773 | 0.791838 | 0.680987 | 15.65758 |
| 207 | 0.68506628 | 0.75996461 | 15.6575773 | 0.789881 | 0.689701 | 15.65758 |
| 208 | 0.6360702 | 0.78975549 | 15.6575773 | 0.809869 | 0.647442 | 15.65758 |
| 209 | 0.58242464 | 0.81682636 | 15.6575773 | 0.724377 | 0.731891 | 15.65758 |
| 210 | 0.68684771 | 0.75712965 | 15.6575773 | 0.804232 | 0.670361 | 15.65758 |
| 211 | 0.75432658 | 0.70731847 | 15.6575773 | 0.772383 | 0.700347 | 15.65758 |
| 212 | 0.74763336 | 0.70352207 | 15.6575773 | 0.79732 | 0.668725 | 15.65758 |
| 213 | 0.7345029 | 0.69096848 | 15.6575773 | 0.820374 | 0.643138 | 15.65758 |
| 214 | 0.38555341 | 0.92473624 | 15.6575773 | 0.786387 | 0.671845 | 15.65758 |
| 215 | 0.66480243 | 0.77317655 | 15.6575773 | 0.722954 | 0.725167 | 15.65758 |
| 216 | 0.53238055 | 0.8501063 | 15.6575773 | 0.661335 | 0.76966 | 15.65758 |
| 217 | 0.49614753 | 0.87117367 | 15.6575773 | 0.747917 | 0.713147 | 15.65758 |
| 218 | 0.62848809 | 0.78155805 | 15.6575773 | 0.754777 | 0.713618 | 15.65758 |
| 219 | 0.34756065 | 0.9461676 | 15.6575773 | 0.812003 | 0.672584 | 15.65758 |
| 220 | 0.71911963 | 0.70564645 | 15.6575773 | 0.752694 | 0.706264 | 15.65758 |
| 221 | 0.59834917 | 0.81128019 | 15.6575773 | 0.790308 | 0.685963 | 15.65758 |
| 222 | 0.47498616 | 0.87943216 | 15.6575773 | 0.836905 | 0.637796 | 15.65758 |
| 223 | 0.60689839 | 0.79634132 | 15.6575773 | 0.799052 | 0.669879 | 15.65758 |
| 224 | 0.50019681 | 0.86724342 | 15.6575773 | 0.793398 | 0.663917 | 15.65758 |
| 225 | 0.62729209 | 0.79058412 | 15.6575773 | 0.781364 | 0.688733 | 15.65758 |
| 226 | 0.46225126 | 0.88865426 | 15.6575773 | 0.780071 | 0.690821 | 15.65758 |
| 227 | 0.67231135 | 0.76832341 | 15.6575773 | 0.787926 | 0.675839 | 15.65758 |
| 228 | 0.64748332 | 0.76768051 | 15.6575773 | 0.82181 | 0.631713 | 15.65758 |
| 229 | 0.62251406 | 0.78420097 | 15.6575773 | 0.843974 | 0.604237 | 15.65758 |
| 230 | 0.8183653 | 0.67757687 | 15.6575773 | 0.731429 | 0.728452 | 15.65758 |
| 231 | 0.75270747 | 0.71496551 | 15.6575773 | 0.763006 | 0.705997 | 15.65758 |
| 232 | 0.6068207 | 0.79963664 | 15.6575773 | 0.74788 | 0.711903 | 15.65758 |
| 233 | 0.58879924 | 0.8080764 | 15.6575773 | 0.750129 | 0.707874 | 15.65758 |
| 234 | 0.19851404 | 1.02630196 | 2 | 0.796418 | 0.649542 | 15.65758 |
| 235 | 0.71390061 | 0.74582221 | 15.6575773 | 0.719688 | 0.729439 | 15.65758 |
| 236 | 0.76283788 | 0.68148595 | 15.6575773 | 0.76701 | 0.698827 | 15.65758 |
| 237 | 0.60732441 | 0.79707316 | 15.6575773 | 0.557045 | 0.836521 | 15.65758 |
| 238 | 0.54137454 | 0.84031461 | 15.6575773 | 0.786279 | 0.675397 | 15.65758 |
| 239 | 0.57237844 | 0.82061417 | 15.6575773 | 0.772489 | 0.694027 | 15.65758 |
| 240 | 0.89292037 | 0.45411744 | 15.6575773 | 0.791821 | 0.645896 | 15.65758 |
| 241 | 0.65583111 | 0.75916878 | 15.6575773 | 0.725282 | 0.728549 | 15.65758 |
| 242 | 0.774149 | 0.70954979 | 15.6575773 | 0.835758 | 0.615321 | 15.65758 |
| 243 | 0.67806547 | 0.76133731 | 15.6575773 | 0.749561 | 0.716408 | 15.65758 |
| 244 | 0.49364517 | 0.86885487 | 15.6575773 | 0.667615 | 0.773075 | 15.65758 |
| 245 | 0.536933 | 0.84322847 | 15.6575773 | 0.765581 | 0.683119 | 15.65758 |
| 246 | 0.70872418 | 0.7067122 | 15.6575773 | 0.734532 | 0.72122 | 15.65758 |
| 247 | 0.77589014 | 0.67296294 | 15.6575773 | 0.674216 | 0.766718 | 15.65758 |
| 248 | 0.80072619 | 0.61393943 | 15.6575773 | 0.803479 | 0.666537 | 15.65758 |
| 249 | 0.65988256 | 0.77526385 | 15.6575773 | 0.787501 | 0.678435 | 15.65758 |
| 250 | 0.68068467 | 0.74091501 | 15.6575773 | 0.76172 | 0.703398 | 15.65758 |
| 251 | 0.73244551 | 0.70928926 | 15.6575773 | 0.795979 | 0.678793 | 15.65758 |
| 252 | 0.75354693 | 0.67422913 | 15.6575773 | 0.776025 | 0.688462 | 15.65758 |
| 253 | 0.63254677 | 0.79478404 | 15.6575773 | 0.798442 | 0.680254 | 15.65758 |
| 254 | 0.6153526 | 0.79113624 | 15.6575773 | 0.738534 | 0.71802 | 15.65758 |
| 255 | 0.56034604 | 0.82753621 | 15.6575773 | 0.787161 | 0.676819 | 15.65758 |
| 256 | 0.77369824 | 0.65969904 | 15.6575773 | 0.744086 | 0.717224 | 15.65758 |
| 257 | 0.85411372 | 0.56176758 | 15.6575773 | 0.786093 | 0.665727 | 15.65758 |
| 258 | 0.57687458 | 0.81685974 | 15.6575773 | 0.801868 | 0.644505 | 15.65758 |
| 259 | 0.69854543 | 0.73007923 | 15.6575773 | 0.735325 | 0.72027 | 15.65758 |
| 260 | 0.26945451 | 0.97889818 | 15.6575773 | 0.820929 | 0.652595 | 15.65758 |
| 261 | 0.84318789 | 0.58198357 | 15.6575773 | 0.742597 | 0.714365 | 15.65758 |
| 262 | 0.69618896 | 0.75954336 | 15.6575773 | 0.81396 | 0.632136 | 15.65758 |
| 263 | 0.44164111 | 0.89711877 | 15.6575773 | 0.825774 | 0.638587 | 15.65758 |
| 264 | 0.63749278 | 0.78479043 | 15.6575773 | 0.705999 | 0.744879 | 15.65758 |
| 265 | 0.86949088 | 0.57450335 | 15.6575773 | 0.817806 | 0.658609 | 15.65758 |
| 266 | 0.8281639 | 0.5636942 | 15.6575773 | 0.768556 | 0.680677 | 15.65758 |
| 267 | 0.66002965 | 0.75860091 | 15.6575773 | 0.849624 | 0.608813 | 15.65758 |
| 268 | 0.75685307 | 0.71151029 | 15.6575773 | 0.737617 | 0.723191 | 15.65758 |
| 269 | 0.57332907 | 0.82288263 | 15.6575773 | 0.710483 | 0.723503 | 15.65758 |
| 270 | 0.84539899 | 0.59464806 | 15.6575773 | 0.784225 | 0.68118 | 15.65758 |
| 271 | 0.76104507 | 0.65077789 | 15.6575773 | 0.728 | 0.728922 | 15.65758 |
| 272 | 0.6640011 | 0.76063509 | 15.6575773 | 0.740465 | 0.716417 | 15.65758 |
| 273 | 0.66142851 | 0.75573128 | 15.6575773 | 0.792563 | 0.679757 | 15.65758 |
| 274 | 0.09561775 | 1.06392067 | 0.92081875 | 0.804311 | 0.656906 | 15.65758 |
| 275 | 0.59691428 | 0.81195173 | 15.6575773 | 0.764963 | 0.695617 | 15.65758 |
| 276 | 0.66214351 | 0.77044357 | 15.6575773 | 0.796255 | 0.672442 | 15.65758 |
| 277 | 0.77865783 | 0.65298519 | 15.6575773 | 0.827325 | 0.657303 | 15.65758 |
| 278 | 0.54618294 | 0.84078972 | 15.6575773 | 0.662821 | 0.768781 | 15.65758 |
| 279 | 0.6933973 | 0.73413878 | 15.6575773 | 0.762835 | 0.700561 | 15.65758 |
| 280 | 0.68076046 | 0.7351417 | 15.6575773 | 0.751751 | 0.695009 | 15.65758 |
| 281 | 0.64173496 | 0.7808481 | 15.6575773 | 0.738474 | 0.711002 | 15.65758 |
| 282 | 0.62576569 | 0.80110413 | 15.6575773 | 0.816107 | 0.653323 | 15.65758 |
| 283 | 0.7198404 | 0.69761934 | 15.6575773 | 0.775223 | 0.682029 | 15.65758 |
| 284 | 0.55056818 | 0.83413069 | 15.6575773 | 0.836076 | 0.628981 | 15.65758 |
| 285 | 0.70713266 | 0.71949422 | 15.6575773 | 0.751681 | 0.701574 | 15.65758 |
| 286 | 0.42489912 | 0.90477606 | 15.6575773 | 0.783174 | 0.692962 | 15.65758 |
| 287 | 0.731716 | 0.71285487 | 15.6575773 | 0.768599 | 0.690812 | 15.65758 |
| 288 | 0.81458549 | 0.62653519 | 15.6575773 | 0.741483 | 0.706972 | 15.65758 |
| 289 | 0.66883619 | 0.74993427 | 15.6575773 | 0.779969 | 0.693146 | 15.65758 |
| 290 | 0.59842802 | 0.80157266 | 15.6575773 | 0.68451 | 0.759285 | 15.65758 |
| 291 | 0.65410643 | 0.77468509 | 15.6575773 | 0.818814 | 0.645443 | 15.65758 |
| 292 | 0.71982128 | 0.70302377 | 15.6575773 | 0.748352 | 0.713861 | 15.65758 |
| 293 | 0.84054467 | 0.58340764 | 15.6575773 | 0.772643 | 0.689538 | 15.65758 |
| 294 | 0.65200613 | 0.78014444 | 15.6575773 | 0.682554 | 0.758181 | 15.65758 |
| 295 | 0.43740953 | 0.8993488 | 15.6575773 | 0.764508 | 0.673325 | 15.65758 |

| | | | | | | |
|---|---|---|---|---|---|---|
| 296 | 0.69769203 | 0.74067558 | 15.6575773 | 0.73447 | 0.722677 | 15.65758 |
| 297 | 0.62942763 | 0.79526233 | 15.6575773 | 0.740265 | 0.722765 | 15.65758 |
| 298 | 0.34273059 | 0.95188053 | 15.6575773 | 0.757274 | 0.714485 | 15.65758 |
| 299 | 0.86048091 | 0.54256793 | 15.6575773 | 0.841756 | 0.592278 | 15.65758 |
| 300 | 0.61614358 | 0.79170506 | 15.6575773 | 0.800507 | 0.654171 | 15.65758 |
| 301 | 0.67335919 | 0.75445336 | 15.6575773 | 0.770732 | 0.701699 | 15.65758 |
| 302 | 0.35345743 | 0.94275013 | 15.6575773 | 0.768785 | 0.698409 | 15.65758 |
| 303 | 0.71042911 | 0.74494089 | 15.6575773 | 0.767138 | 0.701982 | 15.65758 |
| 304 | 0.8477898 | 0.53334989 | 15.6575773 | 0.864484 | 0.58774 | 15.65758 |
| 305 | 0.67903314 | 0.73579422 | 15.6575773 | 0.731853 | 0.725057 | 15.65758 |
| 306 | 0.63507411 | 0.79044464 | 15.6575773 | 0.737426 | 0.715749 | 15.65758 |
| 307 | 0.64005978 | 0.78176292 | 15.6575773 | 0.77896 | 0.686587 | 15.65758 |
| 308 | 0.66719365 | 0.75715937 | 15.6575773 | 0.671694 | 0.759826 | 15.65758 |
| 309 | 0.68957512 | 0.75765528 | 15.6575773 | 0.753502 | 0.716508 | 15.65758 |
| 310 | 0.48493149 | 0.88565917 | 15.6575773 | 0.756836 | 0.700051 | 15.65758 |
| 311 | 0.39856453 | 0.92223475 | 15.6575773 | 0.758049 | 0.707652 | 15.65758 |
| 312 | 0.78137492 | 0.66483924 | 15.6575773 | 0.790423 | 0.68897 | 15.65758 |
| 313 | 0.78942621 | 0.64359881 | 15.6575773 | 0.814714 | 0.667971 | 15.65758 |
| 314 | 0.76627041 | 0.66710032 | 15.6575773 | 0.824011 | 0.654317 | 15.65758 |
| 315 | 0.825058 | 0.58586794 | 15.6575773 | 0.811953 | 0.63457 | 15.65758 |
| 316 | 0.69565437 | 0.72924025 | 15.6575773 | 0.736519 | 0.721321 | 15.65758 |
| 317 | 0.61540866 | 0.79928767 | 15.6575773 | 0.817248 | 0.628069 | 15.65758 |
| 318 | 0.67771209 | 0.75088031 | 15.6575773 | 0.70467 | 0.746934 | 15.65758 |
| 319 | 0.52363313 | 0.85124364 | 15.6575773 | 0.75058 | 0.71488 | 15.65758 |
| 320 | 0.87085539 | 0.54369011 | 15.6575773 | 0.673283 | 0.761646 | 15.65758 |
| 321 | 0.81630204 | 0.61709377 | 15.6575773 | 0.816021 | 0.658796 | 15.65758 |
| 322 | 0.64594725 | 0.7857347 | 15.6575773 | 0.745291 | 0.710347 | 15.65758 |
| 323 | 0.44286124 | 0.89777843 | 15.6575773 | 0.762003 | 0.70251 | 15.65758 |
| 324 | 0.56375425 | 0.82688696 | 15.6575773 | 0.743416 | 0.704938 | 15.65758 |
| 325 | 0.66011937 | 0.77486623 | 15.6575773 | 0.730243 | 0.715486 | 15.65758 |
| 326 | 0.64587618 | 0.7778332 | 15.6575773 | 0.76491 | 0.691397 | 15.65758 |
| 327 | 0.81350517 | 0.63650257 | 15.6575773 | 0.83337 | 0.653997 | 15.65758 |
| 328 | 0.67892547 | 0.74199618 | 15.6575773 | 0.7595 | 0.706742 | 15.65758 |
| 329 | 0.67768412 | 0.75595947 | 15.6575773 | 0.783379 | 0.683794 | 15.65758 |
| 330 | 0.57931722 | 0.82302225 | 15.6575773 | 0.823863 | 0.638998 | 15.65758 |
| 331 | 0.8277393 | 0.60774943 | 15.6575773 | 0.727557 | 0.728405 | 15.65758 |
| 332 | 0.63872241 | 0.78657921 | 15.6575773 | 0.85217 | 0.641538 | 15.65758 |
| 333 | 0.79485073 | 0.61411864 | 15.6575773 | 0.783887 | 0.686954 | 15.65758 |
| 334 | 0.67857081 | 0.75136256 | 15.6575773 | 0.822722 | 0.645722 | 15.65758 |
| 335 | 0.63648825 | 0.78721863 | 15.6575773 | 0.742859 | 0.718157 | 15.65758 |
| 336 | 0.81332719 | 0.61821189 | 15.6575773 | 0.733941 | 0.722593 | 15.65758 |
| 337 | 0.57055958 | 0.82872707 | 15.6575773 | 0.825853 | 0.627373 | 15.65758 |
| 338 | 0.63713995 | 0.78837872 | 15.6575773 | 0.758108 | 0.704373 | 15.65758 |
| 339 | 0.68086742 | 0.75679822 | 15.6575773 | 0.71921 | 0.721512 | 15.65758 |
| 340 | 0.72934189 | 0.72734226 | 15.6575773 | 0.757114 | 0.710346 | 15.65758 |
| 341 | 0.83192681 | 0.64921726 | 15.6575773 | 0.75686 | 0.717091 | 15.65758 |
| 342 | 0.78246491 | 0.63642252 | 15.6575773 | 0.829994 | 0.634412 | 15.65758 |
| 343 | 0.60465147 | 0.80669847 | 15.6575773 | 0.79388 | 0.683538 | 15.65758 |
| 344 | 0.53641883 | 0.84358496 | 15.6575773 | 0.780861 | 0.675445 | 15.65758 |
| 345 | 0.68972396 | 0.74980173 | 15.6575773 | 0.731879 | 0.718151 | 15.65758 |
| 346 | 0.75841901 | 0.68770193 | 15.6575773 | 0.77734 | 0.681967 | 15.65758 |
| 347 | 0.80725538 | 0.64476958 | 15.6575773 | 0.752977 | 0.706004 | 15.65758 |
| 348 | 0.5775612 | 0.86913574 | 15.6575773 | 0.79143 | 0.662081 | 15.65758 |
| 349 | 0.69500163 | 0.75067582 | 15.6575773 | 0.722789 | 0.727697 | 15.65758 |
| 350 | 0.66147917 | 0.7569634 | 15.6575773 | 0.829693 | 0.633217 | 15.65758 |
| 351 | 0.61676661 | 0.79240063 | 15.6575773 | 0.781361 | 0.685954 | 15.65758 |
| 352 | 0.82856033 | 0.60122032 | 15.6575773 | 0.654186 | 0.7776 | 15.65758 |
| 353 | 0.70617192 | 0.72990498 | 15.6575773 | 0.712173 | 0.734328 | 15.65758 |
| 354 | 0.73457706 | 0.68664734 | 15.6575773 | 0.824215 | 0.647241 | 15.65758 |
| 355 | 0.31359627 | 0.9629867 | 15.6575773 | 0.769599 | 0.669602 | 15.65758 |
| 356 | 0.45186184 | 0.89129695 | 15.6575773 | 0.805541 | 0.659896 | 15.65758 |
| 357 | 0.59299302 | 0.81308118 | 15.6575773 | 0.79006 | 0.69253 | 15.65758 |
| 358 | 0.53800577 | 0.84725377 | 15.6575773 | 0.736027 | 0.732259 | 15.65758 |
| 359 | 0.76431043 | 0.70604349 | 15.6575773 | 0.714312 | 0.735142 | 15.65758 |
| 360 | 0.17025679 | 1.03808476 | 2 | 0.761171 | 0.70684 | 15.65758 |
| 361 | 0.68547218 | 0.75350995 | 15.6575773 | 0.688967 | 0.757115 | 15.65758 |
| 362 | 0.66379789 | 0.75864982 | 15.6575773 | 0.827252 | 0.637705 | 15.65758 |
| 363 | 0.73369752 | 0.71775636 | 15.6575773 | 0.818534 | 0.667334 | 15.65758 |
| 364 | 0.64338608 | 0.77553813 | 15.6575773 | 0.835611 | 0.609172 | 15.65758 |
| 365 | 0.69103689 | 0.7409776 | 15.6575773 | 0.776392 | 0.696328 | 15.65758 |
| 366 | 0.64556511 | 0.76519122 | 15.6575773 | 0.766746 | 0.691208 | 15.65758 |
| 367 | 0.69142432 | 0.75020049 | 15.6575773 | 0.824394 | 0.627898 | 15.65758 |
| 368 | 0.71035602 | 0.70416429 | 15.6575773 | 0.74401 | 0.696828 | 15.65758 |
| 369 | 0.57393374 | 0.82158152 | 15.6575773 | 0.80995 | 0.653886 | 15.65758 |
| 370 | 0.7680132 | 0.64222566 | 15.6575773 | 0.822591 | 0.612494 | 15.65758 |
| 371 | 0.51849301 | 0.85746835 | 15.6575773 | 0.728594 | 0.726031 | 15.65758 |
| 372 | 0.74486079 | 0.68363681 | 15.6575773 | 0.768782 | 0.709896 | 15.65758 |
| 373 | 0.71912939 | 0.72351411 | 15.6575773 | 0.7362 | 0.723434 | 15.65758 |
| 374 | 0.66855253 | 0.75030359 | 15.6575773 | 0.802292 | 0.64371 | 15.65758 |
| 375 | 0.71500557 | 0.7087689 | 15.6575773 | 0.728628 | 0.710208 | 15.65758 |
| 376 | 0.57674964 | 0.82619145 | 15.6575773 | 0.759353 | 0.693681 | 15.65758 |
| 377 | 0.75240956 | 0.69466386 | 15.6575773 | 0.763747 | 0.704828 | 15.65758 |
| 378 | 0.73022512 | 0.73899983 | 15.6575773 | 0.772411 | 0.695916 | 15.65758 |
| 379 | 0.71941842 | 0.7101646 | 15.6575773 | 0.73949 | 0.724477 | 15.65758 |
| 380 | 0.67653551 | 0.747337 | 15.6575773 | 0.795691 | 0.664514 | 15.65758 |
| 381 | 0.80205415 | 0.63943573 | 15.6575773 | 0.729668 | 0.723158 | 15.65758 |
| 382 | 0.67812514 | 0.74188468 | 15.6575773 | 0.789552 | 0.671616 | 15.65758 |
| 383 | 0.70053945 | 0.73368547 | 15.6575773 | 0.81552 | 0.665196 | 15.65758 |
| 384 | 0.57307317 | 0.81988861 | 15.6575773 | 0.788056 | 0.68082 | 15.65758 |
| 385 | 0.66552644 | 0.76345452 | 15.6575773 | 0.775621 | 0.694257 | 15.65758 |
| 386 | 0.65402125 | 0.76145584 | 15.6575773 | 0.798758 | 0.641243 | 15.65758 |
| 387 | 0.6513516 | 0.76530441 | 15.6575773 | 0.72543 | 0.73445 | 15.65758 |
| 388 | 0.31307892 | 0.95940365 | 15.6575773 | 0.786517 | 0.681791 | 15.65758 |
| 389 | 0.61140943 | 0.79425297 | 15.6575773 | 0.755092 | 0.696874 | 15.65758 |
| 390 | 0.73779991 | 0.67695028 | 15.6575773 | 0.740136 | 0.722908 | 15.65758 |
| 391 | 0.26437683 | 0.98097019 | 15.6575773 | 0.78094 | 0.683336 | 15.65758 |
| 392 | 0.49064682 | 0.8706901 | 15.6575773 | 0.823799 | 0.640414 | 15.65758 |
| 393 | 0.53421455 | 0.84814431 | 15.6575773 | 0.786727 | 0.677893 | 15.65758 |
| 394 | 0.76957242 | 0.64990945 | 15.6575773 | 0.80014 | 0.675833 | 15.65758 |
| 395 | 0.76838601 | 0.69168466 | 15.6575773 | 0.855299 | 0.613982 | 15.65758 |
| 396 | 0.74465165 | 0.72117884 | 15.6575773 | 0.731772 | 0.717993 | 15.65758 |
| 397 | 0.82607666 | 0.56872887 | 15.6575773 | 0.760158 | 0.703722 | 15.65758 |
| 398 | 0.54116533 | 0.84420808 | 15.6575773 | 0.823615 | 0.648303 | 15.65758 |

| | | | | | | |
|---|---|---|---|---|---|---|
| 399 | 0.75380773 | 0.69201404 | 15.6575773 | 0.754296 | 0.710553 | 15.65758 |
| 400 | 0.70056472 | 0.73641256 | 15.6575773 | 0.828156 | 0.62062 | 15.65758 |
| 401 | 0.30317843 | 0.97100511 | 15.6575773 | 0.744932 | 0.720282 | 15.65758 |
| 402 | 0.42704837 | 0.90400651 | 15.6575773 | 0.760689 | 0.697853 | 15.65758 |
| 403 | 0.73250295 | 0.68569706 | 15.6575773 | 0.743454 | 0.716956 | 15.65758 |
| 404 | 0.63998157 | 0.77148253 | 15.6575773 | 0.797208 | 0.674443 | 15.65758 |
| 405 | 0.18982386 | 1.02679717 | 2 | 0.803261 | 0.64979 | 15.65758 |
| 406 | 0.83707936 | 0.55007538 | 15.6575773 | 0.767848 | 0.693432 | 15.65758 |
| 407 | 0.74339517 | 0.67975992 | 15.6575773 | 0.833875 | 0.634358 | 15.65758 |
| 408 | 0.74793287 | 0.67889893 | 15.6575773 | 0.820188 | 0.623863 | 15.65758 |
| 409 | 0.70750911 | 0.71679886 | 15.6575773 | 0.824295 | 0.654846 | 15.65758 |
| 410 | 0.77867297 | 0.67583158 | 15.6575773 | 0.700145 | 0.749118 | 15.65758 |
| 411 | 0.69289092 | 0.7567346 | 15.6575773 | 0.787082 | 0.681237 | 15.65758 |
| 412 | 0.83317722 | 0.56288769 | 15.6575773 | 0.789655 | 0.679567 | 15.65758 |
| 413 | 0.70542758 | 0.74607318 | 15.6575773 | 0.818843 | 0.651949 | 15.65758 |
| 414 | 0.60418303 | 0.81035751 | 15.6575773 | 0.788503 | 0.683418 | 15.65758 |
| 415 | 0.77263732 | 0.68624395 | 15.6575773 | 0.749337 | 0.713422 | 15.65758 |
| 416 | 0.77999501 | 0.64430522 | 15.6575773 | 0.857768 | 0.596827 | 15.65758 |
| 417 | 0.65819066 | 0.76349394 | 15.6575773 | 0.721292 | 0.738618 | 15.65758 |
| 418 | 0.84534771 | 0.61412027 | 15.6575773 | 0.742444 | 0.719647 | 15.65758 |
| 419 | 0.72107122 | 0.74360552 | 15.6575773 | 0.756215 | 0.697051 | 15.65758 |
| 420 | 0.69186243 | 0.76327487 | 15.6575773 | 0.823636 | 0.661281 | 15.65758 |
| 421 | 0.67704186 | 0.7557326 | 15.6575773 | 0.824067 | 0.644177 | 15.65758 |
| 422 | 0.61801931 | 0.79213051 | 15.6575773 | 0.838228 | 0.606988 | 15.65758 |
| 423 | 0.39774019 | 0.92016505 | 15.6575773 | 0.826148 | 0.66437 | 15.65758 |
| 424 | 0.66725127 | 0.76451426 | 15.6575773 | 0.764849 | 0.688124 | 15.65758 |
| 425 | 0.86750801 | 0.54605993 | 15.6575773 | 0.750045 | 0.692775 | 15.65758 |
| 426 | 0.65224792 | 0.76255366 | 15.6575773 | 0.71395 | 0.738439 | 15.65758 |
| 427 | 0.70968908 | 0.70488503 | 15.6575773 | 0.812222 | 0.649952 | 15.65758 |
| 428 | 0.8144281 | 0.61899063 | 15.6575773 | 0.794581 | 0.670675 | 15.65758 |
| 429 | 0.55162367 | 0.8352793 | 15.6575773 | 0.711564 | 0.739971 | 15.65758 |
| 430 | 0.81451033 | 0.65313459 | 15.6575773 | 0.774304 | 0.678878 | 15.65758 |
| 431 | 0.64322032 | 0.76959099 | 15.6575773 | 0.815513 | 0.651896 | 15.65758 |
| 432 | 0.76139424 | 0.6525384 | 15.6575773 | 0.699841 | 0.747594 | 15.65758 |
| 433 | 0.70921862 | 0.72550022 | 15.6575773 | 0.811825 | 0.637925 | 15.65758 |
| 434 | 0.57747588 | 0.82639924 | 15.6575773 | 0.759339 | 0.701199 | 15.65758 |
| 435 | 0.71591986 | 0.72409077 | 15.6575773 | 0.72731 | 0.729214 | 15.65758 |
| 436 | 0.47779155 | 0.880142 | 15.6575773 | 0.834849 | 0.618359 | 15.65758 |
| 437 | 0.49645427 | 0.86737501 | 15.6575773 | 0.791193 | 0.687659 | 15.65758 |
| 438 | 0.57743264 | 0.82305894 | 15.6575773 | 0.640255 | 0.78536 | 15.65758 |
| 439 | 0.72408992 | 0.73562376 | 15.6575773 | 0.747989 | 0.71532 | 15.65758 |
| 440 | 0.65438578 | 0.76223506 | 15.6575773 | 0.775596 | 0.682084 | 15.65758 |
| 441 | 0.77561601 | 0.673137 | 15.6575773 | 0.664528 | 0.77254 | 15.65758 |
| 442 | 0.65915061 | 0.75561091 | 15.6575773 | 0.744109 | 0.702744 | 15.65758 |
| 443 | 0.64163782 | 0.78081529 | 15.6575773 | 0.823738 | 0.662318 | 15.65758 |
| 444 | 0.63293546 | 0.78185073 | 15.6575773 | 0.728052 | 0.732315 | 15.65758 |
| 445 | 0.47748474 | 0.87907839 | 15.6575773 | 0.704574 | 0.745085 | 15.65758 |
| 446 | 0.79135335 | 0.64989083 | 15.6575773 | 0.723887 | 0.731429 | 15.65758 |
| 447 | 0.78444278 | 0.66513336 | 15.6575773 | 0.794666 | 0.671867 | 15.65758 |
| 448 | 0.66124032 | 0.75088453 | 15.6575773 | 0.836126 | 0.613719 | 15.65758 |
| 449 | 0.84725654 | 0.57327395 | 15.6575773 | 0.755826 | 0.700347 | 15.65758 |
| 450 | 0.6443371 | 0.78954395 | 15.6575773 | 0.75978 | 0.715384 | 15.65758 |
| 451 | 0.64475131 | 0.76588908 | 15.6575773 | 0.770362 | 0.669441 | 15.65758 |
| 452 | 0.74418936 | 0.67489822 | 15.6575773 | 0.855937 | 0.585868 | 15.65758 |
| 453 | 0.72247041 | 0.72688859 | 15.6575773 | 0.762756 | 0.700346 | 15.65758 |
| 454 | 0.51252129 | 0.86008798 | 15.6575773 | 0.743388 | 0.722329 | 15.65758 |
| 455 | 0.22879971 | 1.04332886 | 15.6575773 | 0.758161 | 0.713607 | 15.65758 |
| 456 | 0.49451615 | 0.86879374 | 15.6575773 | 0.784977 | 0.667649 | 15.65758 |
| 457 | 0.61730463 | 0.79382221 | 15.6575773 | 0.809956 | 0.658432 | 15.65758 |
| 458 | 0.72501316 | 0.74525514 | 15.6575773 | 0.803999 | 0.652058 | 15.65758 |
| 459 | 0.69454045 | 0.74772783 | 15.6575773 | 0.788514 | 0.685062 | 15.65758 |
| 460 | 0.63909089 | 0.77040152 | 15.6575773 | 0.765208 | 0.706371 | 15.65758 |
| 461 | 0.66181163 | 0.75262712 | 15.6575773 | 0.787266 | 0.667723 | 15.65758 |
| 462 | 0.55251219 | 0.83822132 | 15.6575773 | 0.829424 | 0.64137 | 15.65758 |
| 463 | 0.80513944 | 0.68464999 | 15.6575773 | 0.744724 | 0.711264 | 15.65758 |
| 464 | 0.70250859 | 0.73723424 | 15.6575773 | 0.768015 | 0.701385 | 15.65758 |
| 465 | 0.59414622 | 0.80920074 | 15.6575773 | 0.848811 | 0.622095 | 15.65758 |
| 466 | 0.61966274 | 0.78486906 | 15.6575773 | 0.796293 | 0.662787 | 15.65758 |
| 467 | 0.52473404 | 0.85249113 | 15.6575773 | 0.808454 | 0.671259 | 15.65758 |
| 468 | 0.60496308 | 0.80896415 | 15.6575773 | 0.742548 | 0.721366 | 15.65758 |
| 469 | 0.61949589 | 0.79070222 | 15.6575773 | 0.811742 | 0.669353 | 15.65758 |
| 470 | 0.69907747 | 0.73108434 | 15.6575773 | 0.73915 | 0.723346 | 15.65758 |
| 471 | 0.74682731 | 0.72107425 | 15.6575773 | 0.754893 | 0.717161 | 15.65758 |
| 472 | 0.68695287 | 0.72899135 | 15.6575773 | 0.782761 | 0.691515 | 15.65758 |
| 473 | 0.70027547 | 0.74764463 | 15.6575773 | 0.76757 | 0.691957 | 15.65758 |
| 474 | 0.67665174 | 0.74817816 | 15.6575773 | 0.78047 | 0.692034 | 15.65758 |
| 475 | 0.73865186 | 0.68108967 | 15.6575773 | 0.859199 | 0.559392 | 15.65758 |
| 476 | 0.52289529 | 0.85180319 | 15.6575773 | 0.668443 | 0.768252 | 15.65758 |
| 477 | 0.33180736 | 0.95022672 | 15.6575773 | 0.716968 | 0.732784 | 15.65758 |
| 478 | 0.59534322 | 0.81261156 | 15.6575773 | 0.815329 | 0.652696 | 15.65758 |
| 479 | 0.702301 | 0.72743886 | 15.6575773 | 0.803292 | 0.672427 | 15.65758 |
| 480 | 0.76350165 | 0.70133905 | 15.6575773 | 0.790684 | 0.678166 | 15.65758 |
| 481 | 0.77611628 | 0.7048954 | 15.6575773 | 0.80878 | 0.667171 | 15.65758 |
| 482 | 0.71592386 | 0.69943891 | 15.6575773 | 0.797176 | 0.647981 | 15.65758 |
| 483 | 0.7804117 | 0.67984787 | 15.6575773 | 0.75695 | 0.702238 | 15.65758 |
| 484 | 0.85593272 | 0.61143133 | 15.6575773 | 0.770794 | 0.690627 | 15.65758 |
| 485 | 0.74228279 | 0.7278532 | 15.6575773 | 0.828818 | 0.628046 | 15.65758 |
| 486 | 0.60588402 | 0.80759028 | 15.6575773 | 0.763802 | 0.697486 | 15.65758 |
| 487 | 0.71823562 | 0.70427794 | 15.6575773 | 0.801204 | 0.669957 | 15.65758 |
| 488 | 0.6636924 | 0.77504323 | 15.6575773 | 0.759127 | 0.700359 | 15.65758 |
| 489 | 0.69713735 | 0.73683862 | 15.6575773 | 0.759708 | 0.706338 | 15.65758 |
| 490 | 0.67121011 | 0.77080842 | 15.6575773 | 0.710417 | 0.741314 | 15.65758 |
| 491 | 0.78699172 | 0.64169254 | 15.6575773 | 0.797238 | 0.676997 | 15.65758 |
| 492 | 0.60100456 | 0.81047883 | 15.6575773 | 0.761262 | 0.706335 | 15.65758 |
| 493 | 0.81389203 | 0.64419667 | 15.6575773 | 0.764715 | 0.697881 | 15.65758 |
| 494 | 0.65987459 | 0.77273028 | 15.6575773 | 0.770325 | 0.700429 | 15.65758 |
| 495 | 0.699245 | 0.74799964 | 15.6575773 | 0.851678 | 0.599122 | 15.65758 |
| 496 | 0.79823767 | 0.65839407 | 15.6575773 | 0.760197 | 0.694588 | 15.65758 |
| 497 | 0.29238768 | 0.97692468 | 15.6575773 | 0.72815 | 0.720757 | 15.65758 |
| 498 | 0.44636892 | 0.89441426 | 15.6575773 | 0.756012 | 0.713311 | 15.65758 |
| 499 | 0.73933841 | 0.70345159 | 15.6575773 | 0.797833 | 0.682087 | 15.65758 |
| 500 | 0.58807227 | 0.81453685 | 15.6575773 | 0.714215 | 0.734489 | 15.65758 |
| 501 | 0.77527752 | 0.71043088 | 15.6575773 | 0.710306 | 0.745301 | 15.65758 |

| | | | | | | |
|---|---|---|---|---|---|---|
| 502 | 0.68434553 | 0.74872971 | 15.6575773 | 0.709174 | 0.745781 | 15.65758 |
| 503 | 0.83047442 | 0.65821388 | 15.6575773 | 0.872401 | 0.59495 | 15.65758 |
| 504 | 0.6976008 | 0.72715378 | 15.6575773 | 0.756366 | 0.70927 | 15.65758 |
| 505 | 0.45165395 | 0.89565841 | 15.6575773 | 0.795147 | 0.685827 | 15.65758 |
| 506 | 0.79364162 | 0.68147898 | 15.6575773 | 0.821229 | 0.585546 | 15.65758 |
| 507 | 0.72811617 | 0.69173956 | 15.6575773 | 0.735752 | 0.722935 | 15.65758 |
| 508 | 0.3902897 | 0.92240038 | 15.6575773 | 0.735647 | 0.712962 | 15.65758 |
| 509 | 0.67516731 | 0.76343468 | 15.6575773 | 0.731546 | 0.696748 | 15.65758 |
| 510 | 0.75947913 | 0.70850583 | 15.6575773 | 0.647702 | 0.782054 | 15.65758 |
| 511 | 0.68924142 | 0.75321705 | 15.6575773 | 0.786892 | 0.686214 | 15.65758 |
| 512 | 0.6297176 | 0.785607 | 15.6575773 | 0.719129 | 0.72712 | 15.65758 |
| 513 | 0.72722106 | 0.72063911 | 15.6575773 | 0.695228 | 0.754684 | 15.65758 |
| 514 | 0.68145715 | 0.74036507 | 15.6575773 | 0.74899 | 0.716309 | 15.65758 |
| 515 | 0.51438259 | 0.85866594 | 15.6575773 | 0.796234 | 0.651528 | 15.65758 |
| 516 | 0.43424789 | 0.9026355 | 15.6575773 | 0.795587 | 0.655116 | 15.65758 |
| 517 | 0.71897431 | 0.7125138 | 15.6575773 | 0.754364 | 0.699159 | 15.65758 |
| 518 | 0.71162204 | 0.73098453 | 15.6575773 | 0.805436 | 0.654452 | 15.65758 |
| 519 | 0.67510939 | 0.74560581 | 15.6575773 | 0.716585 | 0.739683 | 15.65758 |
| 520 | 0.55766011 | 0.83863379 | 15.6575773 | 0.882547 | 0.567297 | 15.65758 |
| 521 | 0.59129059 | 0.80753152 | 15.6575773 | 0.856421 | 0.604516 | 15.65758 |
| 522 | 0.74657949 | 0.66745387 | 15.6575773 | 0.828337 | 0.626874 | 15.65758 |
| 523 | 0.7467179 | 0.6935483 | 15.6575773 | 0.678775 | 0.763916 | 15.65758 |
| 524 | 0.5065352 | 0.86178731 | 15.6575773 | 0.847128 | 0.638282 | 15.65758 |
| 525 | 0.78742553 | 0.69741061 | 15.6575773 | 0.733981 | 0.721383 | 15.65758 |
| 526 | 0.66099005 | 0.75171893 | 15.6575773 | 0.737659 | 0.722559 | 15.65758 |
| 527 | 0.56290276 | 0.83293325 | 15.6575773 | 0.772807 | 0.712841 | 15.65758 |
| 528 | 0.611358 | 0.8006772 | 15.6575773 | 0.683897 | 0.747429 | 15.65758 |
| 529 | 0.74761565 | 0.708675 | 15.6575773 | 0.63868 | 0.789178 | 15.65758 |
| 530 | 0.65065973 | 0.78088125 | 15.6575773 | 0.834756 | 0.639608 | 15.65758 |
| 531 | 0.73842592 | 0.73296533 | 15.6575773 | 0.74666 | 0.707866 | 15.65758 |
| 532 | 0.63274813 | 0.79526962 | 15.6575773 | 0.712008 | 0.733241 | 15.65758 |
| 533 | 0.47611626 | 0.87863921 | 15.6575773 | 0.760208 | 0.706667 | 15.65758 |
| 534 | 0.6269829 | 0.78209819 | 15.6575773 | 0.790969 | 0.682922 | 15.65758 |
| 535 | 0.60912433 | 0.80380809 | 15.6575773 | 0.73324 | 0.729745 | 15.65758 |
| 536 | 0.74499571 | 0.70205367 | 15.6575773 | 0.832417 | 0.653584 | 15.65758 |
| 537 | 0.27125903 | 0.99280928 | 15.6575773 | 0.719435 | 0.731626 | 15.65758 |
| 538 | 0.61933423 | 0.79427226 | 15.6575773 | 0.758461 | 0.694801 | 15.65758 |
| 539 | 0.59298449 | 0.81821617 | 15.6575773 | 0.723226 | 0.724692 | 15.65758 |
| 540 | 0.69632199 | 0.74780338 | 15.6575773 | 0.759046 | 0.701491 | 15.65758 |
| 541 | 0.8432606 | 0.57369159 | 15.6575773 | 0.802775 | 0.672218 | 15.65758 |
| 542 | 0.67268601 | 0.7613017 | 15.6575773 | 0.743187 | 0.721746 | 15.65758 |
| 543 | 0.33926101 | 0.94618681 | 15.6575773 | 0.8015 | 0.671635 | 15.65758 |
| 544 | 0.68111123 | 0.76874461 | 15.6575773 | 0.701943 | 0.74554 | 15.65758 |
| 545 | 0.67881553 | 0.74609584 | 15.6575773 | 0.74564 | 0.721689 | 15.65758 |
| 546 | 0.54046712 | 0.84097136 | 15.6575773 | 0.828306 | 0.638664 | 15.65758 |
| 547 | 0.74375653 | 0.7035178 | 15.6575773 | 0.760761 | 0.709859 | 15.65758 |
| 548 | 0.62636949 | 0.78886823 | 15.6575773 | 0.741058 | 0.720378 | 15.65758 |
| 549 | 0.61438182 | 0.80033816 | 15.6575773 | 0.805449 | 0.600432 | 15.65758 |
| 550 | 0.67019108 | 0.75087283 | 15.6575773 | 0.721638 | 0.732072 | 15.65758 |
| 551 | 0.7569947 | 0.71983827 | 15.6575773 | 0.647006 | 0.784101 | 15.65758 |
| 552 | 0.58326934 | 0.81337943 | 15.6575773 | 0.738943 | 0.71594 | 15.65758 |
| 553 | 0.68913881 | 0.74352379 | 15.6575773 | 0.752531 | 0.662123 | 15.65758 |
| 554 | 0.78771451 | 0.68555313 | 15.6575773 | 0.855221 | 0.539384 | 15.65758 |
| 555 | 0.69969 | 0.75047493 | 15.6575773 | 0.780485 | 0.695053 | 15.65758 |
| 556 | 0.76775235 | 0.64835877 | 15.6575773 | 0.780038 | 0.673303 | 15.65758 |
| 557 | 0.69798146 | 0.72025493 | 15.6575773 | 0.848988 | 0.593368 | 15.65758 |
| 558 | 0.66441686 | 0.7613705 | 15.6575773 | 0.831556 | 0.561812 | 15.65758 |
| 559 | 0.74960655 | 0.68920277 | 15.6575773 | 0.708012 | 0.747469 | 15.65758 |
| 560 | 0.65651583 | 0.76636283 | 15.6575773 | 0.76714 | 0.701602 | 15.65758 |
| 561 | 0.76831415 | 0.69572604 | 15.6575773 | 0.793772 | 0.672402 | 15.65758 |
| 562 | 0.60516233 | 0.80489271 | 15.6575773 | 0.764574 | 0.64737 | 15.65758 |
| 563 | 0.73098764 | 0.72198431 | 15.6575773 | 0.81134 | 0.654706 | 15.65758 |
| 564 | 0.7850624 | 0.64925151 | 15.6575773 | 0.858014 | 0.629467 | 15.65758 |
| 565 | 0.714201 | 0.70662617 | 15.6575773 | 0.857895 | 0.617429 | 15.65758 |
| 566 | 0.1812916 | 1.01924653 | 2 | 0.765667 | 0.668773 | 15.65758 |
| 567 | 0.17227393 | 1.03528054 | 2 | 0.741609 | 0.725371 | 15.65758 |
| 568 | 0.79890394 | 0.66154572 | 15.6575773 | 0.718383 | 0.724958 | 15.65758 |
| 569 | 0.76437784 | 0.6876607 | 15.6575773 | 0.776485 | 0.699374 | 15.65758 |
| 570 | 0.40525942 | 0.91632942 | 15.6575773 | 0.785652 | 0.674369 | 15.65758 |
| 571 | 0.62821331 | 0.790657 | 15.6575773 | 0.79425 | 0.673281 | 15.65758 |
| 572 | 0.54054335 | 0.84392697 | 15.6575773 | 0.748109 | 0.7239 | 15.65758 |
| 573 | 0.76135566 | 0.71534245 | 15.6575773 | 0.764618 | 0.701386 | 15.65758 |
| 574 | 0.68841508 | 0.74224981 | 15.6575773 | 0.700809 | 0.750011 | 15.65758 |
| 575 | 0.5650588 | 0.82783287 | 15.6575773 | 0.873118 | 0.513347 | 15.65758 |
| 576 | 0.61925186 | 0.78776 | 15.6575773 | 0.778786 | 0.645726 | 15.65758 |
| 577 | 0.75747844 | 0.66199282 | 15.6575773 | 0.736532 | 0.728779 | 15.65758 |
| 578 | 0.52336018 | 0.85151132 | 15.6575773 | 0.747596 | 0.714281 | 15.65758 |
| 579 | | | | 0.829477 | 0.607059 | 15.65758 |
| 580 | | | | 0.833785 | 0.659942 | 15.65758 |
| 581 | | | | 0.797244 | 0.663176 | 15.65758 |
| 582 | | | | 0.807191 | 0.666704 | 15.65758 |
| 583 | | | | 0.654101 | 0.769864 | 15.65758 |
| 584 | | | | 0.787908 | 0.686874 | 15.65758 |
| 585 | | | | 0.831786 | 0.653351 | 15.65758 |
| 586 | | | | 0.792458 | 0.677405 | 15.65758 |
| 587 | | | | 0.799621 | 0.676005 | 15.65758 |
| 588 | | | | 0.804854 | 0.659336 | 15.65758 |
| 589 | | | | 0.7692 | 0.68541 | 15.65758 |
| 590 | | | | 0.819506 | 0.645388 | 15.65758 |
| 591 | | | | 0.739765 | 0.716171 | 15.65758 |
| 592 | | | | 0.703424 | 0.728333 | 15.65758 |
| 593 | | | | 0.880527 | 0.599636 | 15.65758 |
| 594 | | | | 0.74548 | 0.717302 | 15.65758 |
| 595 | | | | 0.802857 | 0.672605 | 15.65758 |
| 596 | | | | 0.809635 | 0.592931 | 15.65758 |
| 597 | | | | 0.836957 | 0.646949 | 15.65758 |
| 598 | | | | 0.839056 | 0.6416 | 15.65758 |
| 599 | | | | 0.81157 | 0.680155 | 15.65758 |
| 600 | | | | 0.800692 | 0.655562 | 15.65758 |
| 601 | | | | 0.780288 | 0.698029 | 15.65758 |
| 602 | | | | 0.755929 | 0.689811 | 15.65758 |
| 603 | | | | 0.81974 | 0.663588 | 15.65758 |
| 604 | | | | 0.723193 | 0.731441 | 15.65758 |

| | | | |
|---|---|---|---|
| 605 | 0.836401 | 0.628687 | 15.65758 |
| 606 | 0.911251 | 0.419262 | 15.65758 |
| 607 | 0.818817 | 0.6423 | 15.65758 |
| 608 | 0.815959 | 0.587867 | 15.65758 |
| 609 | 0.775377 | 0.701169 | 15.65758 |
| 610 | 0.810637 | 0.676703 | 15.65758 |
| 611 | 0.783418 | 0.686293 | 15.65758 |
| 612 | 0.827488 | 0.651619 | 15.65758 |
| 613 | 0.830243 | 0.633429 | 15.65758 |
| 614 | 0.76299 | 0.706539 | 15.65758 |
| 615 | 0.777326 | 0.683108 | 15.65758 |
| 616 | 0.792992 | 0.688547 | 15.65758 |
| 617 | 0.791698 | 0.667083 | 15.65758 |
| 618 | 0.746328 | 0.709129 | 15.65758 |
| 619 | 0.824635 | 0.653013 | 15.65758 |
| 620 | 0.780386 | 0.695657 | 15.65758 |
| 621 | 0.890327 | 0.548828 | 15.65758 |
| 622 | 0.765789 | 0.695786 | 15.65758 |
| 623 | 0.793323 | 0.67593 | 15.65758 |
| 624 | 0.799253 | 0.685113 | 15.65758 |
| 625 | 0.707269 | 0.713864 | 15.65758 |
| 626 | 0.815809 | 0.659695 | 15.65758 |
| 627 | 0.756838 | 0.708618 | 15.65758 |
| 628 | 0.758913 | 0.712994 | 15.65758 |
| 629 | 0.806806 | 0.670664 | 15.65758 |
| 630 | 0.725551 | 0.702166 | 15.65758 |
| 631 | 0.789219 | 0.677406 | 15.65758 |
| 632 | 0.798177 | 0.671401 | 15.65758 |
| 633 | 0.771027 | 0.67813 | 15.65758 |
| 634 | 0.809229 | 0.666031 | 15.65758 |
| 635 | 0.752316 | 0.7117 | 15.65758 |
| 636 | 0.745579 | 0.711261 | 15.65758 |
| 637 | 0.745375 | 0.717513 | 15.65758 |
| 638 | 0.828201 | 0.622256 | 15.65758 |
| 639 | 0.819101 | 0.644291 | 15.65758 |
| 640 | 0.783208 | 0.690717 | 15.65758 |
| 641 | 0.639238 | 0.786195 | 15.65758 |
| 642 | 0.728835 | 0.728735 | 15.65758 |
| 643 | 0.5941 | 0.810263 | 15.65758 |
| 644 | 0.743587 | 0.716656 | 15.65758 |
| 645 | 0.819951 | 0.659432 | 15.65758 |
| 646 | 0.71859 | 0.736019 | 15.65758 |
| 647 | 0.817048 | 0.659215 | 15.65758 |
| 648 | 0.814462 | 0.666707 | 15.65758 |
| 649 | 0.737011 | 0.71207 | 15.65758 |
| 650 | 0.751921 | 0.716585 | 15.65758 |
| 651 | 0.771256 | 0.690252 | 15.65758 |
| 652 | 0.779024 | 0.688401 | 15.65758 |
| 653 | 0.784727 | 0.669316 | 15.65758 |
| 654 | 0.763146 | 0.688507 | 15.65758 |
| 655 | 0.761684 | 0.700649 | 15.65758 |
| 656 | 0.744078 | 0.718388 | 15.65758 |
| 657 | 0.839238 | 0.631888 | 15.65758 |
| 658 | 0.712365 | 0.70875 | 15.65758 |
| 659 | 0.781958 | 0.696883 | 15.65758 |
| 660 | 0.764964 | 0.708114 | 15.65758 |
| 661 | 0.781655 | 0.694766 | 15.65758 |
| 662 | 0.88273 | 0.513199 | 15.65758 |
| 663 | 0.708519 | 0.743325 | 15.65758 |
| 664 | 0.78099 | 0.685178 | 15.65758 |
| 665 | 0.705255 | 0.723169 | 15.65758 |
| 666 | 0.78502 | 0.685646 | 15.65758 |
| 667 | 0.696065 | 0.734747 | 15.65758 |
| 668 | 0.832355 | 0.645889 | 15.65758 |
| 669 | 0.687384 | 0.756173 | 15.65758 |
| 670 | 0.778759 | 0.689232 | 15.65758 |
| 671 | 0.739144 | 0.727058 | 15.65758 |
| 672 | 0.766936 | 0.700403 | 15.65758 |
| 673 | 0.831052 | 0.656697 | 15.65758 |
| 674 | 0.76816 | 0.703657 | 15.65758 |
| 675 | 0.80481 | 0.656479 | 15.65758 |
| 676 | 0.830248 | 0.662945 | 15.65758 |
| 677 | 0.799644 | 0.675043 | 15.65758 |
| 678 | 0.755421 | 0.682801 | 15.65758 |
| 679 | 0.807969 | 0.663481 | 15.65758 |
| 680 | 0.778601 | 0.703357 | 15.65758 |
| 681 | 0.678404 | 0.759408 | 15.65758 |
| 682 | 0.844275 | 0.562663 | 15.65758 |
| 683 | 0.771829 | 0.692486 | 15.65758 |
| 684 | 0.827057 | 0.658848 | 15.65758 |
| 685 | 0.8889 | 0.520816 | 15.65758 |
| 686 | 0.779557 | 0.685926 | 15.65758 |
| 687 | 0.826954 | 0.656704 | 15.65758 |
| 688 | 0.769125 | 0.699903 | 15.65758 |
| 689 | 0.881961 | 0.496594 | 15.65758 |
| 690 | 0.710025 | 0.745151 | 15.65758 |
| 691 | 0.815741 | 0.637419 | 15.65758 |
| 692 | 0.864562 | 0.58562 | 15.65758 |
| 693 | 0.731239 | 0.725962 | 15.65758 |
| 694 | 0.725731 | 0.733802 | 15.65758 |
| 695 | 0.809398 | 0.650864 | 15.65758 |
| 696 | 0.844813 | 0.629186 | 15.65758 |
| 697 | 0.772659 | 0.696847 | 15.65758 |
| 698 | 0.785803 | 0.684959 | 15.65758 |
| 699 | 0.875638 | 0.559683 | 15.65758 |
| 700 | 0.745452 | 0.708584 | 15.65758 |
| 701 | 0.629989 | 0.794424 | 15.65758 |
| 702 | 0.814794 | 0.658299 | 15.65758 |
| 703 | 0.755485 | 0.715518 | 15.65758 |
| 704 | 0.831513 | 0.634615 | 15.65758 |
| 705 | 0.852114 | 0.583726 | 15.65758 |
| 706 | 0.849181 | 0.608123 | 15.65758 |
| 707 | 0.766739 | 0.672542 | 15.65758 |
| 708 | 0.687113 | 0.751077 | 15.65758 |
| 709 | 0.775649 | 0.668987 | 15.65758 |
| 710 | 0.73186 | 0.728451 | 15.65758 |
| 711 | 0.794779 | 0.668587 | 15.65758 |
| 712 | 0.746058 | 0.714136 | 15.65758 |
| 713 | 0.783579 | 0.684422 | 15.65758 |
| 714 | 0.793903 | 0.623058 | 15.65758 |
| 715 | 0.797154 | 0.664462 | 15.65758 |
| 716 | 0.829359 | 0.654034 | 15.65758 |
| 717 | 0.794461 | 0.67782 | 15.65758 |
| 718 | 0.746753 | 0.713377 | 15.65758 |
| 719 | 0.764302 | 0.709912 | 15.65758 |
| 720 | 0.821783 | 0.669849 | 15.65758 |
| 721 | 0.600036 | 0.803218 | 15.65758 |
| 722 | 0.773492 | 0.697938 | 15.65758 |
| 723 | 0.748952 | 0.708535 | 15.65758 |
| 724 | 0.802202 | 0.669293 | 15.65758 |
| 725 | 0.872055 | 0.608634 | 15.65758 |
| 726 | 0.821294 | 0.652607 | 15.65758 |
| 727 | 0.726352 | 0.722836 | 15.65758 |
| 728 | 0.659818 | 0.765166 | 15.65758 |
| 729 | 0.782503 | 0.684321 | 15.65758 |
| 730 | 0.688529 | 0.744634 | 15.65758 |
| 731 | 0.773486 | 0.705232 | 15.65758 |
| 732 | 0.691429 | 0.749325 | 15.65758 |
| 733 | 0.71483 | 0.729752 | 15.65758 |
| 734 | 0.78104 | 0.684301 | 15.65758 |
| 735 | 0.75907 | 0.690635 | 15.65758 |
| 736 | 0.610114 | 0.800979 | 15.65758 |
| 737 | 0.756082 | 0.695542 | 15.65758 |
| 738 | 0.76771 | 0.686997 | 15.65758 |
| 739 | 0.804666 | 0.665904 | 15.65758 |
| 740 | 0.664061 | 0.753398 | 15.65758 |
| 741 | 0.810006 | 0.663316 | 15.65758 |
| 742 | 0.749778 | 0.721999 | 15.65758 |
| 743 | 0.810336 | 0.665186 | 15.65758 |
| 744 | 0.712186 | 0.735793 | 15.65758 |
| 745 | 0.783865 | 0.679325 | 15.65758 |
| 746 | 0.796486 | 0.682023 | 15.65758 |
| 747 | 0.796407 | 0.671123 | 15.65758 |
| 748 | 0.804208 | 0.675282 | 15.65758 |
| 749 | 0.800447 | 0.654541 | 15.65758 |
| 750 | 0.767746 | 0.693829 | 15.65758 |
| 751 | 0.732253 | 0.728506 | 15.65758 |
| 752 | 0.751563 | 0.699475 | 15.65758 |
| 753 | 0.81538 | 0.675123 | 15.65758 |
| 754 | 0.823838 | 0.657871 | 15.65758 |
| 755 | 0.758944 | 0.710429 | 15.65758 |
| 756 | 0.695817 | 0.739976 | 15.65758 |
| 757 | 0.826663 | 0.628033 | 15.65758 |
| 758 | 0.853517 | 0.595221 | 15.65758 |
| 759 | 0.797715 | 0.681695 | 15.65758 |
| 760 | 0.785589 | 0.691973 | 15.65758 |
| 761 | 0.795076 | 0.679921 | 15.65758 |
| 762 | 0.768809 | 0.719103 | 15.65758 |
| 763 | 0.84491 | 0.646007 | 15.65758 |
| 764 | 0.773846 | 0.684647 | 15.65758 |
| 765 | 0.84135 | 0.614592 | 15.65758 |
| 766 | 0.808888 | 0.672955 | 15.65758 |
| 767 | 0.782672 | 0.682618 | 15.65758 |
| 768 | 0.7018 | 0.747612 | 15.65758 |
| 769 | 0.820353 | 0.624411 | 15.65758 |
| 770 | 0.763721 | 0.692248 | 15.65758 |
| 771 | 0.770611 | 0.680024 | 15.65758 |
| 772 | 0.725137 | 0.731628 | 15.65758 |
| 773 | 0.739857 | 0.718038 | 15.65758 |
| 774 | 0.855136 | 0.583444 | 15.65758 |
| 775 | 0.752022 | 0.712958 | 15.65758 |
| 776 | 0.736903 | 0.721651 | 15.65758 |
| 777 | 0.793606 | 0.622588 | 15.65758 |
| 778 | 0.822497 | 0.644963 | 15.65758 |
| 779 | 0.825582 | 0.619915 | 15.65758 |
| 780 | 0.772236 | 0.692406 | 15.65758 |
| 781 | 0.791914 | 0.653098 | 15.65758 |
| 782 | 0.808445 | 0.672087 | 15.65758 |
| 783 | 0.793581 | 0.687456 | 15.65758 |
| 784 | 0.679075 | 0.760631 | 15.65758 |
| 785 | 0.775771 | 0.674573 | 15.65758 |
| 786 | 0.855273 | 0.603479 | 15.65758 |
| 787 | 0.766861 | 0.699522 | 15.65758 |
| 788 | 0.833812 | 0.63632 | 15.65758 |
| 789 | 0.671904 | 0.767981 | 15.65758 |
| 790 | 0.627788 | 0.780373 | 15.65758 |
| 791 | 0.691161 | 0.754252 | 15.65758 |
| 792 | 0.800067 | 0.633382 | 15.65758 |
| 793 | 0.681481 | 0.747214 | 15.65758 |
| 794 | 0.896852 | 0.506732 | 15.65758 |
| 795 | 0.826931 | 0.646884 | 15.65758 |
| 796 | 0.81027 | 0.685558 | 15.65758 |
| 797 | 0.833139 | 0.601492 | 15.65758 |
| 798 | 0.866313 | 0.595174 | 15.65758 |
| 799 | 0.761802 | 0.70396 | 15.65758 |
| 800 | 0.708014 | 0.73629 | 15.65758 |
| 801 | 0.777637 | 0.693333 | 15.65758 |
| 802 | 0.820679 | 0.64775 | 15.65758 |
| 803 | 0.783761 | 0.667992 | 15.65758 |
| 804 | 0.745293 | 0.71143 | 15.65758 |
| 805 | 0.722151 | 0.743919 | 15.65758 |
| 806 | 0.761253 | 0.710745 | 15.65758 |
| 807 | 0.73444 | 0.683018 | 15.65758 |
| 808 | 0.768755 | 0.704782 | 15.65758 |
| 809 | 0.746894 | 0.717701 | 15.65758 |
| 810 | 0.707796 | 0.743212 | 15.65758 |

| | | | |
|---|---|---|---|
| 811 | 0.691524 | 0.75594 | 15.65758 |
| 812 | 0.78693 | 0.666508 | 15.65758 |
| 813 | 0.745628 | 0.698497 | 15.65758 |
| 814 | 0.74929 | 0.713002 | 15.65758 |
| 815 | 0.768514 | 0.698464 | 15.65758 |
| 816 | 0.842632 | 0.584736 | 15.65758 |
| 817 | 0.7051 | 0.730224 | 15.65758 |
| 818 | 0.76721 | 0.699879 | 15.65758 |
| 819 | 0.787893 | 0.671389 | 15.65758 |
| 820 | 0.780374 | 0.682974 | 15.65758 |
| 821 | 0.785363 | 0.692493 | 15.65758 |
| 822 | 0.859342 | 0.549345 | 15.65758 |
| 823 | 0.767217 | 0.704088 | 15.65758 |
| 824 | 0.739796 | 0.720221 | 15.65758 |
| 825 | 0.728577 | 0.733938 | 15.65758 |
| 826 | 0.733453 | 0.726341 | 15.65758 |
| 827 | 0.732387 | 0.72961 | 15.65758 |
| 828 | 0.815105 | 0.630482 | 15.65758 |
| 829 | 0.839441 | 0.65636 | 15.65758 |
| 830 | 0.834698 | 0.650292 | 15.65758 |
| 831 | 0.771269 | 0.704258 | 15.65758 |
| 832 | 0.789611 | 0.686827 | 15.65758 |
| 833 | 0.735719 | 0.72393 | 15.65758 |
| 834 | 0.814591 | 0.616849 | 15.65758 |
| 835 | 0.790419 | 0.689815 | 15.65758 |
| 836 | 0.78473 | 0.682878 | 15.65758 |
| 837 | 0.813936 | 0.656387 | 15.65758 |
| 838 | 0.829929 | 0.611358 | 15.65758 |
| 839 | 0.772189 | 0.694661 | 15.65758 |
| 840 | 0.707238 | 0.720096 | 15.65758 |
| 841 | 0.685376 | 0.754889 | 15.65758 |
| 842 | 0.769271 | 0.691431 | 15.65758 |
| 843 | 0.744316 | 0.714659 | 15.65758 |
| 844 | 0.880236 | 0.542133 | 15.65758 |
| 845 | 0.760082 | 0.702195 | 15.65758 |
| 846 | 0.763591 | 0.702477 | 15.65758 |
| 847 | 0.809324 | 0.672497 | 15.65758 |
| 848 | 0.822625 | 0.612721 | 15.65758 |
| 849 | 0.766008 | 0.712739 | 15.65758 |
| 850 | 0.785518 | 0.69913 | 15.65758 |
| 851 | 0.675955 | 0.765904 | 15.65758 |
| 852 | 0.800859 | 0.602692 | 15.65758 |
| 853 | 0.858401 | 0.561668 | 15.65758 |
| 854 | 0.769692 | 0.692131 | 15.65758 |
| 855 | 0.816124 | 0.659429 | 15.65758 |
| 856 | 0.771042 | 0.692388 | 15.65758 |
| 857 | 0.771223 | 0.686221 | 15.65758 |
| 858 | 0.670844 | 0.76087 | 15.65758 |
| 859 | 0.737992 | 0.727044 | 15.65758 |
| 860 | 0.672762 | 0.762582 | 15.65758 |
| 861 | 0.887975 | 0.483904 | 15.65758 |
| 862 | 0.73406 | 0.723293 | 15.65758 |
| 863 | 0.816208 | 0.655008 | 15.65758 |
| 864 | 0.841274 | 0.657643 | 15.65758 |
| 865 | 0.799198 | 0.687463 | 15.65758 |
| 866 | 0.835788 | 0.629502 | 15.65758 |
| 867 | 0.777863 | 0.689189 | 15.65758 |
| 868 | 0.695276 | 0.752255 | 15.65758 |
| 869 | 0.712226 | 0.736885 | 15.65758 |
| 870 | 0.773715 | 0.697803 | 15.65758 |
| 871 | 0.79545 | 0.687114 | 15.65758 |
| 872 | 0.765891 | 0.688616 | 15.65758 |
| 873 | 0.795922 | 0.683263 | 15.65758 |
| 874 | 0.769179 | 0.682906 | 15.65758 |
| 875 | 0.760249 | 0.70093 | 15.65758 |
| 876 | 0.819652 | 0.651266 | 15.65758 |
| 877 | 0.70846 | 0.709759 | 15.65758 |
| 878 | 0.803284 | 0.649877 | 15.65758 |
| 879 | 0.80757 | 0.666417 | 15.65758 |
| 880 | 0.775111 | 0.697829 | 15.65758 |
| 881 | 0.763048 | 0.704407 | 15.65758 |
| 882 | 0.768287 | 0.701009 | 15.65758 |
| 883 | 0.767751 | 0.694695 | 15.65758 |
| 884 | 0.822764 | 0.634876 | 15.65758 |
| 885 | 0.850516 | 0.640262 | 15.65758 |
| 886 | 0.591747 | 0.816696 | 15.65758 |
| 887 | 0.73863 | 0.725974 | 15.65758 |
| 888 | 0.748382 | 0.720216 | 15.65758 |
| 889 | 0.804458 | 0.656342 | 15.65758 |
| 890 | 0.769842 | 0.692604 | 15.65758 |
| 891 | 0.778384 | 0.691461 | 15.65758 |
| 892 | 0.728208 | 0.728898 | 15.65758 |
| 893 | 0.81855 | 0.672021 | 15.65758 |
| 894 | 0.704846 | 0.746881 | 15.65758 |
| 895 | 0.72821 | 0.730615 | 15.65758 |
| 896 | 0.77736 | 0.658292 | 15.65758 |
| 897 | 0.701517 | 0.744183 | 15.65758 |
| 898 | 0.784194 | 0.643621 | 15.65758 |
| 899 | 0.802241 | 0.66083 | 15.65758 |
| 900 | 0.825347 | 0.655826 | 15.65758 |
| 901 | 0.783879 | 0.671634 | 15.65758 |
| 902 | 0.812159 | 0.651685 | 15.65758 |
| 903 | 0.787115 | 0.68212 | 15.65758 |
| 904 | 0.765787 | 0.695202 | 15.65758 |
| 905 | 0.789955 | 0.691247 | 15.65758 |
| 906 | 0.743388 | 0.691064 | 15.65758 |
| 907 | 0.657245 | 0.76763 | 15.65758 |
| 908 | 0.379121 | 0.93263 | 15.65758 |
| 909 | 0.778152 | 0.68659 | 15.65758 |
| 910 | 0.708266 | 0.741077 | 15.65758 |
| 911 | 0.741949 | 0.703251 | 15.65758 |
| 912 | 0.732645 | 0.709869 | 15.65758 |
| 913 | 0.733462 | 0.719083 | 15.65758 |

| | | | |
|---|---|---|---|
| 914 | 0.716453 | 0.742462 | 15.65758 |
| 915 | 0.654978 | 0.773194 | 15.65758 |
| 916 | 0.625046 | 0.795415 | 15.65758 |
| 917 | 0.714263 | 0.730403 | 15.65758 |
| 918 | 0.770429 | 0.693096 | 15.65758 |
| 919 | 0.749573 | 0.693034 | 15.65758 |
| 920 | 0.772131 | 0.695857 | 15.65758 |
| 921 | 0.766798 | 0.691671 | 15.65758 |
| 922 | 0.756766 | 0.698513 | 15.65758 |
| 923 | 0.640414 | 0.779451 | 15.65758 |
| 924 | 0.750113 | 0.706342 | 15.65758 |
| 925 | 0.719802 | 0.728077 | 15.65758 |
| 926 | 0.777063 | 0.679308 | 15.65758 |
| 927 | 0.663746 | 0.766772 | 15.65758 |
| 928 | 0.719106 | 0.712363 | 15.65758 |
| 929 | 0.751487 | 0.692049 | 15.65758 |
| 930 | 0.790472 | 0.674228 | 15.65758 |
| 931 | 0.605095 | 0.802587 | 15.65758 |
| 932 | 0.755214 | 0.715396 | 15.65758 |
| 933 | 0.697897 | 0.74966 | 15.65758 |
| 934 | 0.785667 | 0.692032 | 15.65758 |
| 935 | 0.739355 | 0.719973 | 15.65758 |
| 936 | 0.741586 | 0.708798 | 15.65758 |
| 937 | 0.771365 | 0.688013 | 15.65758 |
| 938 | 0.740556 | 0.698675 | 15.65758 |
| 939 | 0.696002 | 0.751329 | 15.65758 |
| 940 | 0.806964 | 0.656185 | 15.65758 |
| 941 | 0.681813 | 0.755193 | 15.65758 |
| 942 | 0.668765 | 0.764835 | 15.65758 |
| 943 | 0.677381 | 0.757851 | 15.65758 |
| 944 | 0.751058 | 0.663902 | 15.65758 |
| 945 | 0.732774 | 0.689789 | 15.65758 |
| 946 | 0.67583 | 0.757583 | 15.65758 |
| 947 | 0.771558 | 0.680492 | 15.65758 |
| 948 | 0.697133 | 0.727558 | 15.65758 |
| 949 | 0.810621 | 0.659512 | 15.65758 |
| 950 | 0.784512 | 0.665114 | 15.65758 |
| 951 | 0.840039 | 0.615174 | 15.65758 |
| 952 | 0.694421 | 0.744816 | 15.65758 |
| 953 | 0.712999 | 0.7328 | 15.65758 |
| 954 | 0.838536 | 0.616874 | 15.65758 |
| 955 | 0.65724 | 0.771597 | 15.65758 |
| 956 | 0.744268 | 0.710347 | 15.65758 |
| 957 | 0.729235 | 0.72432 | 15.65758 |
| 958 | 0.70442 | 0.729031 | 15.65758 |
| 959 | 0.775332 | 0.695943 | 15.65758 |
| 960 | 0.630596 | 0.789718 | 15.65758 |
| 961 | 0.701197 | 0.723258 | 15.65758 |
| 962 | 0.806464 | 0.633677 | 15.65758 |
| 963 | 0.703745 | 0.725434 | 15.65758 |
| 964 | 0.582657 | 0.822075 | 15.65758 |
| 965 | 0.700899 | 0.717601 | 15.65758 |
| 966 | 0.696693 | 0.742772 | 15.65758 |
| 967 | 0.720849 | 0.714114 | 15.65758 |
| 968 | 0.716702 | 0.729774 | 15.65758 |
| 969 | 0.756091 | 0.676041 | 15.65758 |
| 970 | 0.761053 | 0.694696 | 15.65758 |
| 971 | 0.825623 | 0.619598 | 15.65758 |
| 972 | 0.760872 | 0.685556 | 15.65758 |
| 973 | 0.767353 | 0.701338 | 15.65758 |
| 974 | 0.751818 | 0.716972 | 15.65758 |
| 975 | 0.672787 | 0.758048 | 15.65758 |
| 976 | 0.70122 | 0.735332 | 15.65758 |
| 977 | 0.548242 | 0.841881 | 15.65758 |
| 978 | 0.653347 | 0.778939 | 15.65758 |
| 979 | 0.657986 | 0.758886 | 15.65758 |
| 980 | 0.693729 | 0.739627 | 15.65758 |
| 981 | 0.75958 | 0.68512 | 15.65758 |
| 982 | 0.787659 | 0.682513 | 15.65758 |
| 983 | 0.783072 | 0.673346 | 15.65758 |
| 984 | 0.852235 | 0.565936 | 15.65758 |
| 985 | 0.756683 | 0.704229 | 15.65758 |
| 986 | 0.776682 | 0.70108 | 15.65758 |
| 987 | 0.744442 | 0.717364 | 15.65758 |
| 988 | 0.705559 | 0.729741 | 15.65758 |
| 989 | 0.691645 | 0.735575 | 15.65758 |
| 990 | 0.766992 | 0.682072 | 15.65758 |
| 991 | 0.723486 | 0.723949 | 15.65758 |
| 992 | 0.624424 | 0.793921 | 15.65758 |
| 993 | 0.224089 | 1.000564 | 15.65758 |
| 994 | 0.742459 | 0.701235 | 15.65758 |
| 995 | 0.669515 | 0.762791 | 15.65758 |
| 996 | 0.800951 | 0.650713 | 15.65758 |
| 997 | 0.688341 | 0.743621 | 15.65758 |
| 998 | 0.680654 | 0.742041 | 15.65758 |
| 999 | 0.750087 | 0.70924 | 15.65758 |
| 1000 | 0.781958 | 0.684516 | 15.65758 |
| 1001 | 0.837911 | 0.624383 | 15.65758 |
| 1002 | 0.771139 | 0.686127 | 15.65758 |
| 1003 | 0.735688 | 0.722035 | 15.65758 |
| 1004 | 0.747783 | 0.686443 | 15.65758 |
| 1005 | 0.727815 | 0.690349 | 15.65758 |
| 1006 | 0.762345 | 0.700186 | 15.65758 |
| 1007 | 0.718939 | 0.72389 | 15.65758 |
| 1008 | 0.746073 | 0.719766 | 15.65758 |
| 1009 | 0.705781 | 0.738709 | 15.65758 |
| 1010 | 0.793995 | 0.65613 | 15.65758 |
| 1011 | 0.723311 | 0.723929 | 15.65758 |
| 1012 | 0.629387 | 0.786832 | 15.65758 |
| 1013 | 0.77112 | 0.669028 | 15.65758 |
| 1014 | 0.77787 | 0.66907 | 15.65758 |
| 1015 | 0.669405 | 0.76766 | 15.65758 |
| 1016 | 0.705858 | 0.742072 | 15.65758 |

| | | | |
|---|---|---|---|
| 1017 | 0.680278 | 0.753871 | 15.65758 |
| 1018 | 0.753652 | 0.662044 | 15.65758 |
| 1019 | 0.820061 | 0.647687 | 15.65758 |
| 1020 | 0.72079 | 0.723806 | 15.65758 |
| 1021 | 0.776586 | 0.688046 | 15.65758 |
| 1022 | 0.687348 | 0.754433 | 15.65758 |
| 1023 | 0.690358 | 0.739867 | 15.65758 |
| 1024 | 0.783328 | 0.661857 | 15.65758 |
| 1025 | 0.506323 | 0.861537 | 15.65758 |
| 1026 | 0.67595 | 0.754128 | 15.65758 |
| 1027 | 0.757751 | 0.710844 | 15.65758 |
| 1028 | 0.75452 | 0.709793 | 15.65758 |
| 1029 | 0.757314 | 0.696604 | 15.65758 |
| 1030 | 0.815508 | 0.612907 | 15.65758 |
| 1031 | 0.71877 | 0.729066 | 15.65758 |
| 1032 | 0.743899 | 0.715105 | 15.65758 |
| 1033 | 0.790251 | 0.658635 | 15.65758 |
| 1034 | 0.732677 | 0.729678 | 15.65758 |
| 1035 | 0.35503 | 0.964937 | 15.65758 |
| 1036 | 0.709782 | 0.76099 | 15.65758 |
| 1037 | 0.531547 | 0.847224 | 15.65758 |
| 1038 | 0.732107 | 0.723587 | 15.65758 |
| 1039 | 0.744643 | 0.674368 | 15.65758 |
| 1040 | 0.627713 | 0.793194 | 15.65758 |
| 1041 | 0.756276 | 0.688384 | 15.65758 |
| 1042 | 0.653043 | 0.780371 | 15.65758 |
| 1043 | 0.560027 | 0.833088 | 15.65758 |
| 1044 | 0.711066 | 0.725508 | 15.65758 |
| 1045 | 0.756527 | 0.710687 | 15.65758 |
| 1046 | 0.818695 | 0.634769 | 15.65758 |
| 1047 | 0.695607 | 0.735414 | 15.65758 |
| 1048 | 0.714009 | 0.72878 | 15.65758 |
| 1049 | 0.712248 | 0.741944 | 15.65758 |
| 1050 | 0.775953 | 0.672263 | 15.65758 |
| 1051 | 0.744133 | 0.703175 | 15.65758 |
| 1052 | 0.772121 | 0.684686 | 15.65758 |
| 1053 | 0.713474 | 0.728495 | 15.65758 |
| 1054 | 0.771891 | 0.685924 | 15.65758 |
| 1055 | 0.809628 | 0.654636 | 15.65758 |
| 1056 | 0.760103 | 0.692118 | 15.65758 |
| 1057 | 0.78302 | 0.678274 | 15.65758 |
| 1058 | 0.745836 | 0.714558 | 15.65758 |
| 1059 | 0.768741 | 0.697168 | 15.65758 |
| 1060 | 0.704826 | 0.749205 | 15.65758 |
| 1061 | 0.775161 | 0.683603 | 15.65758 |
| 1062 | 0.732497 | 0.702145 | 15.65758 |
| 1063 | 0.794772 | 0.653777 | 15.65758 |
| 1064 | 0.704785 | 0.741416 | 15.65758 |
| 1065 | 0.610918 | 0.803309 | 15.65758 |
| 1066 | 0.73975 | 0.703612 | 15.65758 |
| 1067 | 0.578734 | 0.819566 | 15.65758 |
| 1068 | 0.755852 | 0.712753 | 15.65758 |
| 1069 | 0.757024 | 0.697733 | 15.65758 |
| 1070 | 0.790732 | 0.687048 | 15.65758 |
| 1071 | 0.63718 | 0.77523 | 15.65758 |
| 1072 | 0.672559 | 0.764661 | 15.65758 |
| 1073 | 0.551601 | 0.837156 | 15.65758 |
| 1074 | 0.838523 | 0.610499 | 15.65758 |
| 1075 | 0.78146 | 0.649677 | 15.65758 |
| 1076 | 0.757695 | 0.682712 | 15.65758 |
| 1077 | 0.765066 | 0.691627 | 15.65758 |
| 1078 | 0.798581 | 0.656941 | 15.65758 |
| 1079 | 0.66402 | 0.761724 | 15.65758 |
| 1080 | 0.720704 | 0.730526 | 15.65758 |
| 1081 | 0.660943 | 0.768078 | 15.65758 |
| 1082 | 0.819114 | 0.637775 | 15.65758 |
| 1083 | 0.74674 | 0.695437 | 15.65758 |
| 1084 | 0.709397 | 0.741257 | 15.65758 |
| 1085 | 0.711678 | 0.738512 | 15.65758 |
| 1086 | 0.688208 | 0.753062 | 15.65758 |
| 1087 | 0.793762 | 0.656493 | 15.65758 |
| 1088 | 0.708538 | 0.730728 | 15.65758 |
| 1089 | 0.801037 | 0.669775 | 15.65758 |
| 1090 | 0.641376 | 0.776633 | 15.65758 |
| 1091 | 0.736028 | 0.720062 | 15.65758 |
| 1092 | 0.674931 | 0.768255 | 15.65758 |
| 1093 | 0.776621 | 0.659614 | 15.65758 |
| 1094 | 0.767389 | 0.680831 | 15.65758 |
| 1095 | 0.790682 | 0.632699 | 15.65758 |
| 1096 | 0.841569 | 0.63514 | 15.65758 |
| 1097 | 0.743903 | 0.704644 | 15.65758 |
| 1098 | 0.805389 | 0.618343 | 15.65758 |
| 1099 | 0.696449 | 0.748044 | 15.65758 |
| 1100 | 0.719927 | 0.715891 | 15.65758 |
| 1101 | 0.69704 | 0.73399 | 15.65758 |
| 1102 | 0.651679 | 0.771676 | 15.65758 |
| 1103 | 0.755278 | 0.670244 | 15.65758 |
| 1104 | 0.720316 | 0.722533 | 15.65758 |
| 1105 | 0.737407 | 0.719694 | 15.65758 |
| 1106 | 0.662154 | 0.766654 | 15.65758 |
| 1107 | 0.830692 | 0.631771 | 15.65758 |
| 1108 | 0.791845 | 0.666042 | 15.65758 |
| 1109 | 0.649861 | 0.777237 | 15.65758 |
| 1110 | 0.686171 | 0.760505 | 15.65758 |
| 1111 | 0.816981 | 0.643317 | 15.65758 |
| 1112 | 0.764978 | 0.679219 | 15.65758 |
| 1113 | 0.614046 | 0.795592 | 15.65758 |
| 1114 | 0.639398 | 0.78214 | 15.65758 |
| 1115 | 0.704238 | 0.733455 | 15.65758 |
| 1116 | 0.755563 | 0.695235 | 15.65758 |
| 1117 | 0.785704 | 0.692678 | 15.65758 |
| 1118 | 0.758935 | 0.708937 | 15.65758 |
| 1119 | 0.781297 | 0.676508 | 15.65758 |
| 1120 | 0.71271 | 0.741053 | 15.65758 |
| 1121 | 0.590683 | 0.814181 | 15.65758 |
| 1122 | 0.717742 | 0.722819 | 15.65758 |
| 1123 | 0.692886 | 0.739174 | 15.65758 |
| 1124 | 0.630878 | 0.791091 | 15.65758 |
| 1125 | 0.735303 | 0.720045 | 15.65758 |
| 1126 | 0.631029 | 0.780588 | 15.65758 |
| 1127 | 0.722456 | 0.724475 | 15.65758 |
| 1128 | 0.531931 | 0.848575 | 15.65758 |
| 1129 | 0.70148 | 0.746213 | 15.65758 |
| 1130 | 0.763469 | 0.685513 | 15.65758 |
| 1131 | 0.639119 | 0.773494 | 15.65758 |
| 1132 | 0.772997 | 0.687042 | 15.65758 |
| 1133 | 0.828557 | 0.623948 | 15.65758 |
| 1134 | 0.695512 | 0.755683 | 15.65758 |
| 1135 | 0.742055 | 0.690035 | 15.65758 |
| 1136 | 0.6752 | 0.765561 | 15.65758 |
| 1137 | 0.813241 | 0.638919 | 15.65758 |
| 1138 | 0.800103 | 0.673278 | 15.65758 |
| 1139 | 0.778431 | 0.663085 | 15.65758 |
| 1140 | 0.665075 | 0.758563 | 15.65758 |
| 1141 | 0.239762 | 0.998118 | 15.65758 |
| 1142 | 0.66556 | 0.757436 | 15.65758 |
| 1143 | 0.700001 | 0.735559 | 15.65758 |
| 1144 | 0.678126 | 0.762227 | 15.65758 |
| 1145 | 0.747102 | 0.705114 | 15.65758 |
| 1146 | 0.803634 | 0.660102 | 15.65758 |
| 1147 | 0.783473 | 0.669712 | 15.65758 |
| 1148 | 0.747155 | 0.708985 | 15.65758 |
| 1149 | 0.725198 | 0.702334 | 15.65758 |
| 1150 | 0.685829 | 0.75145 | 15.65758 |
| 1151 | 0.502274 | 0.864317 | 15.65758 |
| 1152 | 0.782728 | 0.672239 | 15.65758 |
| 1153 | 0.72986 | 0.727101 | 15.65758 |
| 1154 | 0.773939 | 0.697421 | 15.65758 |
| 1155 | 0.687333 | 0.743767 | 15.65758 |
| 1156 | 0.707543 | 0.731405 | 15.65758 |
| 1157 | 0.845588 | 0.611515 | 15.65758 |
| 1158 | 0.710151 | 0.732487 | 15.65758 |
| 1159 | 0.641018 | 0.772645 | 15.65758 |
| 1160 | 0.661364 | 0.776176 | 15.65758 |
| 1161 | 0.808727 | 0.653302 | 15.65758 |
| 1162 | 0.766915 | 0.69704 | 15.65758 |
| 1163 | 0.686071 | 0.753808 | 15.65758 |
| 1164 | 0.599673 | 0.805603 | 15.65758 |
| 1165 | 0.794448 | 0.629683 | 15.65758 |
| 1166 | 0.802811 | 0.644151 | 15.65758 |
| 1167 | 0.651216 | 0.772093 | 15.65758 |
| 1168 | 0.795941 | 0.674203 | 15.65758 |
| 1169 | 0.72548 | 0.711849 | 15.65758 |
| 1170 | 0.739106 | 0.712871 | 15.65758 |
| 1171 | 0.730707 | 0.71683 | 15.65758 |
| 1172 | 0.698956 | 0.743106 | 15.65758 |
| 1173 | 0.809202 | 0.661838 | 15.65758 |
| 1174 | 0.821564 | 0.578432 | 15.65758 |
| 1175 | 0.632834 | 0.784984 | 15.65758 |
| 1176 | 0.577107 | 0.821497 | 15.65758 |
| 1177 | 0.707338 | 0.741701 | 15.65758 |
| 1178 | 0.715981 | 0.736117 | 15.65758 |
| 1179 | 0.764327 | 0.675078 | 15.65758 |
| 1180 | 0.807443 | 0.655474 | 15.65758 |
| 1181 | 0.681317 | 0.739627 | 15.65758 |
| 1182 | 0.73531 | 0.704112 | 15.65758 |
| 1183 | 0.66009 | 0.755459 | 15.65758 |
| 1184 | 0.740376 | 0.714956 | 15.65758 |
| 1185 | 0.776033 | 0.672729 | 15.65758 |
| 1186 | 0.740137 | 0.69061 | 15.65758 |
| 1187 | 0.63157 | 0.787747 | 15.65758 |
| 1188 | 0.754081 | 0.697833 | 15.65758 |
| 1189 | 0.74844 | 0.685375 | 15.65758 |
| 1190 | 0.727209 | 0.735502 | 15.65758 |
| 1191 | 0.656472 | 0.77443 | 15.65758 |
| 1192 | 0.796227 | 0.658925 | 15.65758 |
| 1193 | 0.772166 | 0.672837 | 15.65758 |
| 1194 | 0.715671 | 0.736194 | 15.65758 |
| 1195 | 0.52654 | 0.849773 | 15.65758 |
| 1196 | 0.732145 | 0.726056 | 15.65758 |
| 1197 | 0.705633 | 0.734931 | 15.65758 |
| 1198 | 0.768547 | 0.702023 | 15.65758 |
| 1199 | 0.71499 | 0.738296 | 15.65758 |
| 1200 | 0.764234 | 0.694914 | 15.65758 |
| 1201 | 0.673909 | 0.756292 | 15.65758 |
| 1202 | 0.682181 | 0.744902 | 15.65758 |
| 1203 | 0.774757 | 0.684742 | 15.65758 |
| 1204 | 0.680445 | 0.757292 | 15.65758 |
| 1205 | 0.69218 | 0.741157 | 15.65758 |
| 1206 | 0.76709 | 0.67628 | 15.65758 |
| 1207 | 0.736914 | 0.707589 | 15.65758 |
| 1208 | 0.708526 | 0.729553 | 15.65758 |
| 1209 | 0.708515 | 0.728573 | 15.65758 |
| 1210 | 0.740751 | 0.717606 | 15.65758 |
| 1211 | 0.623889 | 0.794416 | 15.65758 |
| 1212 | 0.807316 | 0.656808 | 15.65758 |
| 1213 | 0.762856 | 0.706352 | 15.65758 |
| 1214 | 0.712044 | 0.736508 | 15.65758 |
| 1215 | 0.588046 | 0.817042 | 15.65758 |
| 1216 | 0.755743 | 0.686084 | 15.65758 |
| 1217 | 0.757567 | 0.666973 | 15.65758 |
| 1218 | 0.278226 | 0.974871 | 15.65758 |
| 1219 | 0.725258 | 0.723067 | 15.65758 |
| 1220 | 0.70005 | 0.736196 | 15.65758 |
| 1221 | 0.58577 | 0.814576 | 15.65758 |
| 1222 | 0.723187 | 0.720456 | 15.65758 |

| | | | |
|---|---|---|---|
| 1223 | 0.714422 | 0.730269 | 15.65758 |
| 1224 | 0.577502 | 0.822202 | 15.65758 |
| 1225 | 0.693312 | 0.751261 | 15.65758 |
| 1226 | 0.791282 | 0.685651 | 15.65758 |
| 1227 | 0.564387 | 0.829423 | 15.65758 |
| 1228 | 0.747661 | 0.704331 | 15.65758 |
| 1229 | 0.720655 | 0.713909 | 15.65758 |
| 1230 | 0.662961 | 0.769072 | 15.65758 |
| 1231 | 0.72137 | 0.721816 | 15.65758 |
| 1232 | 0.686161 | 0.748134 | 15.65758 |
| 1233 | 0.809834 | 0.661821 | 15.65758 |
| 1234 | 0.76412 | 0.698151 | 15.65758 |
| 1235 | 0.689511 | 0.733263 | 15.65758 |
| 1236 | 0.792592 | 0.657457 | 15.65758 |
| 1237 | 0.766558 | 0.712442 | 15.65758 |
| 1238 | 0.761543 | 0.706918 | 15.65758 |
| 1239 | 0.550016 | 0.837194 | 15.65758 |
| 1240 | 0.730906 | 0.726227 | 15.65758 |
| 1241 | 0.793114 | 0.681477 | 15.65758 |
| 1242 | 0.784768 | 0.672592 | 15.65758 |
| 1243 | 0.747936 | 0.716881 | 15.65758 |
| 1244 | 0.725548 | 0.721379 | 15.65758 |
| 1245 | 0.629535 | 0.794369 | 15.65758 |
| 1246 | 0.839963 | 0.619946 | 15.65758 |
| 1247 | 0.630581 | 0.786691 | 15.65758 |
| 1248 | 0.802012 | 0.65231 | 15.65758 |
| 1249 | 0.704737 | 0.732707 | 15.65758 |
| 1250 | 0.684682 | 0.75004 | 15.65758 |
| 1251 | 0.751803 | 0.681922 | 15.65758 |
| 1252 | 0.727949 | 0.725403 | 15.65758 |
| 1253 | 0.695998 | 0.751598 | 15.65758 |
| 1254 | 0.484333 | 0.874303 | 15.65758 |
| 1255 | 0.698134 | 0.743205 | 15.65758 |
| 1256 | 0.714551 | 0.727829 | 15.65758 |
| 1257 | 0.66066 | 0.782367 | 15.65758 |
| 1258 | 0.785997 | 0.641922 | 15.65758 |
| 1259 | 0.784753 | 0.691787 | 15.65758 |
| 1260 | 0.779707 | 0.685682 | 15.65758 |
| 1261 | 0.825568 | 0.627843 | 15.65758 |
| 1262 | 0.72802 | 0.711189 | 15.65758 |
| 1263 | 0.75862 | 0.696339 | 15.65758 |
| 1264 | 0.747344 | 0.700294 | 15.65758 |
| 1265 | 0.694342 | 0.746211 | 15.65758 |
| 1266 | 0.860389 | 0.599314 | 15.65758 |
| 1267 | 0.759755 | 0.703841 | 15.65758 |
| 1268 | 0.642002 | 0.777851 | 15.65758 |
| 1269 | 0.776608 | 0.633207 | 15.65758 |
| 1270 | 0.744324 | 0.704364 | 15.65758 |
| 1271 | 0.741291 | 0.712387 | 15.65758 |
| 1272 | 0.744685 | 0.694415 | 15.65758 |
| 1273 | 0.702378 | 0.745417 | 15.65758 |
| 1274 | 0.779596 | 0.6634 | 15.65758 |
| 1275 | 0.756676 | 0.701942 | 15.65758 |
| 1276 | 0.677364 | 0.771027 | 15.65758 |
| 1277 | 0.714177 | 0.733026 | 15.65758 |
| 1278 | 0.722679 | 0.730909 | 15.65758 |
| 1279 | 0.702159 | 0.743194 | 15.65758 |
| 1280 | 0.74375 | 0.702985 | 15.65758 |
| 1281 | 0.812572 | 0.642327 | 15.65758 |
| 1282 | 0.720098 | 0.735015 | 15.65758 |
| 1283 | 0.749565 | 0.69603 | 15.65758 |
| 1284 | 0.75384 | 0.709555 | 15.65758 |
| 1285 | 0.779544 | 0.686506 | 15.65758 |
| 1286 | 0.585673 | 0.817444 | 15.65758 |
| 1287 | 0.62267 | 0.793432 | 15.65758 |
| 1288 | 0.75075 | 0.679543 | 15.65758 |
| 1289 | 0.769074 | 0.714154 | 15.65758 |
| 1290 | 0.672913 | 0.755533 | 15.65758 |
| 1291 | 0.791189 | 0.677605 | 15.65758 |
| 1292 | 0.687371 | 0.748525 | 15.65758 |
| 1293 | 0.728507 | 0.727564 | 15.65758 |
| 1294 | 0.712912 | 0.744501 | 15.65758 |
| 1295 | 0.770852 | 0.664634 | 15.65758 |
| 1296 | 0.722733 | 0.720011 | 15.65758 |
| 1297 | 0.779218 | 0.666609 | 15.65758 |
| 1298 | 0.647898 | 0.772769 | 15.65758 |
| 1299 | 0.796565 | 0.6778 | 15.65758 |
| 1300 | 0.777649 | 0.685544 | 15.65758 |
| 1301 | 0.697163 | 0.750899 | 15.65758 |
| 1302 | 0.723201 | 0.736601 | 15.65758 |
| 1303 | 0.663783 | 0.771957 | 15.65758 |
| 1304 | 0.79649 | 0.632939 | 15.65758 |
| 1305 | 0.716883 | 0.734674 | 15.65758 |
| 1306 | 0.826048 | 0.635109 | 15.65758 |
| 1307 | 0.81099 | 0.665067 | 15.65758 |
| 1308 | 0.721699 | 0.720644 | 15.65758 |
| 1309 | 0.779726 | 0.691522 | 15.65758 |
| 1310 | 0.707097 | 0.750393 | 15.65758 |
| 1311 | 0.768244 | 0.65827 | 15.65758 |
| 1312 | 0.693818 | 0.742331 | 15.65758 |
| 1313 | 0.654357 | 0.774034 | 15.65758 |
| 1314 | 0.731014 | 0.727412 | 15.65758 |
| 1315 | 0.820342 | 0.606148 | 15.65758 |
| 1316 | 0.719218 | 0.72763 | 15.65758 |
| 1317 | 0.792234 | 0.649942 | 15.65758 |
| 1318 | 0.607372 | 0.801541 | 15.65758 |
| 1319 | 0.767538 | 0.694009 | 15.65758 |
| 1320 | 0.672924 | 0.771797 | 15.65758 |
| 1321 | 0.700652 | 0.744471 | 15.65758 |
| 1322 | 0.704809 | 0.723656 | 15.65758 |
| 1323 | 0.814738 | 0.629158 | 15.65758 |
| 1324 | 0.790582 | 0.628781 | 15.65758 |
| 1325 | 0.718023 | 0.713256 | 15.65758 |
| 1326 | 0.783174 | 0.692256 | 15.65758 |
| 1327 | 0.675715 | 0.755211 | 15.65758 |
| 1328 | 0.659093 | 0.769057 | 15.65758 |
| 1329 | 0.668308 | 0.758882 | 15.65758 |
| 1330 | 0.669046 | 0.752132 | 15.65758 |
| 1331 | 0.744776 | 0.714642 | 15.65758 |
| 1332 | 0.670585 | 0.764935 | 15.65758 |
| 1333 | 0.68995 | 0.740489 | 15.65758 |
| 1334 | 0.607847 | 0.796431 | 15.65758 |
| 1335 | 0.754045 | 0.70841 | 15.65758 |
| 1336 | 0.611958 | 0.805104 | 15.65758 |
| 1337 | 0.746985 | 0.705477 | 15.65758 |
| 1338 | 0.753129 | 0.69097 | 15.65758 |
| 1339 | 0.825727 | 0.661501 | 15.65758 |
| 1340 | 0.697593 | 0.745985 | 15.65758 |
| 1341 | 0.621878 | 0.792016 | 15.65758 |
| 1342 | 0.760842 | 0.704345 | 15.65758 |
| 1343 | 0.771142 | 0.677164 | 15.65758 |
| 1344 | 0.775522 | 0.678941 | 15.65758 |
| 1345 | 0.488443 | 0.872785 | 15.65758 |
| 1346 | 0.687731 | 0.754729 | 15.65758 |
| 1347 | 0.72357 | 0.736731 | 15.65758 |
| 1348 | 0.756148 | 0.677644 | 15.65758 |
| 1349 | 0.727772 | 0.709451 | 15.65758 |

**Table S2. Phenotypic characteristics of GTEx lung tissue donors**

| Feature | Number of individuals (%) | Median (range) |
|---|---|---|
| Sex | | |
| Female | 183 (31.7) | |
| Male | 395 (68.3) | |
| Age | | |
| <40 | 73 (12.6) | 56 (21 - 70) |
| 40-49 | 93 (16.1) | |
| 50-59 | 200 (34.6) | |
| 60-69 | 190 (32.9) | |
| >69 | 22 (3.8) | |
| Body mass index | | |
| <25 | 155 (26.8) | 27.7 (17.0 - 35.4) |
| 25-30 | 246 (42.6) | |
| >30 | 177 (30.6) | |
| Race | | |
| Caucasian | 493 (85.9) | |
| Black/African American | 70 (12.2) | |
| Asian | 10 (1.7) | |
| American Indian/Alaska Native | 1 (0.2) | |
| Smoking status | | |
| Non-smoker | 180 (32.0) | |
| Smoker | 382 (68.0) | |

**Table S3. Multivariate analysis of ACE2 expression in human lung tissue**

| Variable | $\beta$ value | S.E. | $t$ value | $p$ value | |
|---|---|---|---|---|---|
| CD8+ T cells | -2.719 | 0.584 | -4.656 | **4.02e-06** | **** |
| NK cells resting | -3.128 | 0.583 | -5.366 | **1.18e-07** | **** |
| NK cells activated | -2.281 | 0.704 | -3.241 | **0.0013** | ** |
| M1 macrophages | -4.963 | 1.339 | -3.707 | **0.0002** | *** |
| CD4+ T cells | 0.453 | 0.513 | 0.882 | 0.3781 | |
| Dendritic cells | -12.793 | 6.688 | -1.913 | 0.0563 | |
| Neutrophils | -0.398 | 0.469 | -0.848 | 0.3971 | |
| Sex = male (vs. female) | 0.065 | 0.056 | 1.16 | 0.2466 | |
| Age | 0.004 | 0.002 | 1.799 | 0.0726 | |
| BMI | -0.004 | 0.006 | -0.556 | 0.5787 | |
| Race = Caucasian (vs. non-Caucasian) | -0.096 | 0.073 | -1.313 | 0.1899 | |
| Smoking status = yes (vs. no) | -0.025 | 0.058 | -0.433 | 0.6650 | |

Abbreviations: S.E., Standard Error.

**Table S4. Meta-analysis of ACE2 mRNA and protein levels in 17 human tissues**
*Source data of Fig. 2f.*

| Source data type | Source data method | Adipose tissue | Airway epithelium | Brain | Cardiac muscle | Colorectal tissue | Endometrium | Gall bladder | Kidney | Liver | Lung | Oral mucosa/esophagus | Skin | Small intestine | Smooth muscle | Spleen | Stomach | Testis | Ref. |
|---|---|---|---|---|---|---|---|---|---|---|---|---|---|---|---|---|---|---|---|
| mRNA | NB | 0 | N | N | 3 | 2 | 0 | N | 3 | 0 | 0 | N | 0 | 2 | 0 | N | N | 3 | [1] |
| mRNA | qRT-PCR | 1 | 1 | 0 | 2 | 2 | 1 | 2 | 2 | 1 | 1 | 1 | 0 | 3 | 1 | 0 | 1 | 3 | [2] |
| mRNA | MA | 2 | 2 | 2 | 2 | 2 | 1 | N | 3 | 2 | 2 | N | 1 | 3 | 2 | N | N | 3 | [3] |
| mRNA | RNAseq | 2 | N | 2 | 2 | 1 | 1 | 3 | 3 | 0 | 0 | 0 | 1 | 3 | 2 | N | 0 | 3 | [4] |
| mRNA | RNAseq | N | N | N | N | N | N | N | N | N | N | 1 | N | N | N | N | N | N | [5] |
| mRNA | CAGE | 0 | N | 0 | 1 | 2 | 0 | 1 | 1 | 0 | 0 | 0 | N | 3 | 0 | 0 | N | 2 | [6] |
| Protein | MS | N | N | 0 | 1 | 0 | N | 1 | 3 | 1 | 0 | 0 | N | N | N | N | N | 3 | [7] |
| Protein | IHC | 3 | 1 | 1 | N | 1 | N | N | 3 | 1 | 3 | 2 | 3 | 3 | 3 | 3 | 0 | N | [8] |
| Protein | IHC | 0 | 0 | 0 | 1 | 2 | 0 | 3 | 3 | 0 | 0 | 0 | 0 | 3 | 0 | 0 | 0 | 3 | [9] |
| **Scores (averages)** | | | | | | | | | | | | | | | | | | | |
| **mRNA score** | | 1.0 | 1.5 | 1.0 | 2.0 | 1.8 | 0.6 | 2.0 | 2.4 | 0.6 | 0.6 | 0.5 | 0.5 | 2.8 | 1.0 | 0.0 | 0.5 | 2.8 | |
| **Protein score** | | 1.5 | 0.5 | 0.3 | 1.0 | 1.0 | 0.0 | 2.0 | 3.0 | 0.7 | 1.0 | 0.7 | 1.5 | 3.0 | 1.5 | 1.5 | 0.0 | 3.0 | |

*Abbreviations:* CAGE, cap analysis of gene expression; IHC, immunohistochemistry; MA, microarray; MS, mass spectrometry; N, not determined; NB Northern blot.
